# The Blood RNA Stability Atlas: defining temporal structure and trait-state programs in the human whole-blood transcriptome

**DOI:** 10.64898/2026.07.07.737044

**Authors:** Will C. Baltazar, Robert O. Messing, Laura B. Ferguson

**Author notes:** Corresponding Author: Laura Ferguson, Ph.D. Assistant Professor, Department of Pharmacology and Department of Psychiatry Bowles Center for Alcohol Studies, University of North Carolina School of Medicine, 104 Manning Dr, Chapel Hill, NC 27599.

## Abstract

Whole-blood RNA biomarkers are widely used for diagnosis and disease monitoring, but their utility depends not only on abundance but also on temporal stability, a property that is not routinely incorporated into biomarker design. We analyzed 968 longitudinal whole-blood transcriptomes from 165 healthy individuals across eight independent studies spanning diverse platforms, time scales (50 minutes to 16 weeks), and common environmental exposures. Using a cross-study analytical framework integrating variance partitioning, repeatability, and time-associated differential expression, we quantified temporal stability for 6,064 RNAs and classified transcripts into “trait” (stable) and “state” (dynamic) categories representing the extremes of longitudinal changes in transcript abundance. We identified 1,118 trait RNAs exhibiting stable within-individual levels of abundance but substantial inter-individual variability, enriched for whole-blood eGenes (P = 6.0 × 10⁻²⁰), supporting a genetic basis for stability. In contrast, 1,504 state RNAs showed context-dependent temporal variation and were enriched for translation and RNA-binding pathways. Integration with genetic datasets revealed that 4,395 (72%) blood transcripts were linked to at least one whole-blood eQTL, collectively associated with 18,358 GWAS trait relationships, providing disease-relevant context for transcript stability. We developed the Blood RNA Stability Atlas to integrate these features and demonstrate both top-down (disease-to-gene) and bottom-up (gene-to-context) applications for biomarker prioritization and interpretation. These findings establish temporal stability as a defining property of the blood transcriptome and provide a practical, publicly accessible framework for distinguishing stable baseline abundance levels from context-dependent transcriptional responses, informing biomarker selection, study design, and hypothesis generation.

## Introduction

Progress in medical genomics and personalized medicine has been accelerated in recent decades with the advent of -omics methodologies including Next Generation Sequencing (NGS), the establishment of large repositories for -omics and clinical datasets, and the development of computational methods to extract meaningful information from these high-dimensional datasets [1, 2]. The transcriptome comprises the complete repertoire of RNA molecules expressed within a biological sample and provides a dynamic readout of cellular function, making transcriptomic profiling a powerful tool for studying biology and discovering biomarkers [3]. It can reflect the extent of transcript abundance in different cell types, tissues, individuals, and environmental contexts, making it a rich source of inter-individual and context-dependent biological variation that researchers have used to explain differences in source sample phenotypes.

Biomarkers are measurements that indicate the biological state of an organism. While a broad range of measurements qualify as biomarkers, molecular-scale biomarkers such as RNA have made headway in the -omics revolution due to their technological accessibility, comprehensiveness, and specificity for an individual [4]. Biomarkers can be broadly conceptualized as trait-like or state-like depending on whether they capture relatively stable biological characteristics or dynamic physiological changes. To establish clear definitions for biomarkers and their endpoints, a joint FDA-NIH commission created the Biomarkers, EndpointS, and other Tools (BEST) resource [5]. The BEST delineates seven types of biomarkers reviewed in detail by Califf [6]: diagnostic, monitoring, pharmacodynamic/response (herein referred to as response), predictive, prognostic, safety, and susceptibility/risk (herein referred to as risk). In terms of the broad classification as trait-like or state-like biomarkers, trait-like biomarkers include diagnostic, predictive, prognostic, and risk biomarkers, and state-like biomarkers include monitoring, response, predictive, and safety biomarkers. Collectively, these efforts reflect the continued evolution of biomarker science toward increasingly multimodal, data-driven, and clinically actionable approaches for precision medicine [7].

Among accessible tissues, peripheral blood has emerged as an attractive source of transcriptomic biomarkers because it can be sampled repeatedly with minimal invasiveness and reflects diverse physiological and pathological processes [8–10]. These factors make blood a popular target for biomarker discovery and preferable to tissues that require invasive methods to obtain, such as cerebrospinal fluid or solid-tissue biopsies. Blood is routinely collected in clinics and fractionated into its constituent parts, making it a promising candidate for the advancement of personalized medicine, and its rapid time to storage using dedicated blood RNA tubes can make whole blood a more advantageous tissue than its cellular and cell-free fractions [11]. The COVID-19 pandemic accelerated the expansion of molecular diagnostic tools in the clinical environment, including in low-resource settings, which has further elevated RNA to the forefront of clinical biomarker research globally [12, 13]. As the clinical use of blood RNA biomarkers expands, foundational knowledge of transcript behavior in healthy individuals becomes increasingly essential.

Most whole blood transcriptome studies seek to identify RNA signatures associated with disease, typically through cross-sectional comparisons between healthy and diseased individuals. These investigations rarely include repeated sampling of healthy controls to establish the baseline temporal variability of the healthy transcriptome. However, the recognition that peripheral blood contains reproducible, individual-specific transcriptomic signatures dates back more than two decades [14]. Recent work has begun to characterize this structure at genome scale. An eight-year longitudinal microarray study found that the most temporally stable whole-blood transcripts were enriched for immune and interferon-signaling pathways, whereas transcripts involved in cellular maintenance, signaling, apoptosis, and cell-cycle regulation exhibited greater long-term temporal change [15]. Longitudinal RNA-sequencing of platelets similarly demonstrated that many platelet transcripts remain remarkably stable over four years and showed that highly stable transcripts are enriched for expression and splice quantitative trait loci, linking temporal stability to underlying genetic regulation [16]. Most recently, a population-scale study of 333 healthy individuals sampled longitudinally reported that most genes exhibit greater within-individual than between-individual variation, whereas genes that remain stable within individuals while differing consistently between individuals are enriched for T- and B-cell–associated transcripts and show greater heritability [17]. Together, these studies demonstrate that although many blood transcripts fluctuate over time within individuals, a subset of immune-related genes remains remarkably stable within individuals while differing reproducibly between individuals, forming genetically influenced individual-specific expression signatures that provide an important foundation for blood biomarker discovery.

Despite these advances, researchers lack a general framework for evaluating transcript stability across studies or for assessing individual candidate biomarkers. Existing longitudinal references are each limited to a single cohort, platform, or sampling design, leaving it unclear whether transcript stability generalizes across experimental settings and they provide no practical resource for investigators to evaluate the temporal stability of candidate biomarkers. Here we addressed this gap by harmonizing eight longitudinal whole-blood transcriptomic studies spanning microarray and RNA sequencing platforms and sampling intervals from 50 minutes to 16 weeks. By integrating diverse study designs, we identified transcripts whose temporal behavior is reproducible across cohorts and technologies rather than specific to any individual dataset. We quantified temporal stability for 6,064 transcripts measured across all studies and identified transcripts exhibiting trait-like or state-like expression behavior. Our cross-platform consensus captured the immune-related stability and genetic regulation patterns reported in previous longitudinal studies, demonstrating that these signatures generalize across fundamentally different study designs. Finally, to facilitate biomarker discovery and interpretation we developed the Blood RNA Stability Atlas, a publicly available interactive resource that provides transcript classifications, stability metrics, longitudinal expression trajectories, and linked eQTL and GWAS associations, supporting both disease-to-transcript and transcript-to-context biomarker workflows.

## Results

### Datasets

To establish a framework for prioritizing RNA biomarkers based on temporal changes in abundance, we analyzed publicly available longitudinal whole-blood transcriptomic datasets from healthy individuals. Eight studies met our selection criteria, comprising 968 samples from 165 individuals (**Table 1**). The studies spanned multiple populations, experimental conditions, measurement platforms, and sampling intervals ranging from minutes to months. This diversity enabled us to evaluate temporal transcript behavior across multiple sources of biological and technical variation.

**Table 1.**
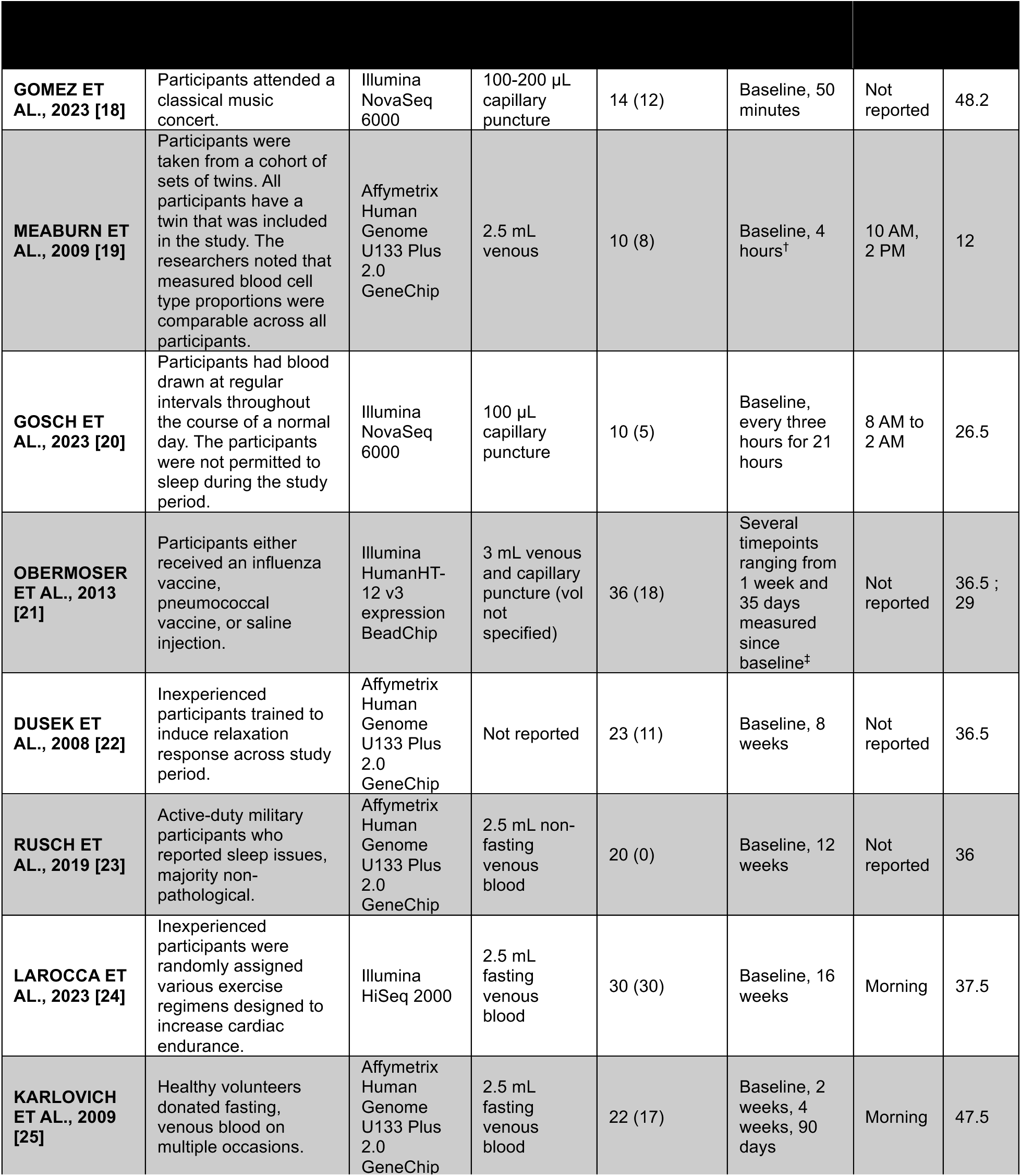
Characteristics of the Studies Used for Analysis. Blood was collected in each study via venipuncture or capillary puncture or was otherwise not reported. Whole blood was collected in blood tubes containing RNA-stabilizing media and RNA was extracted according to methods specified by each study. 968 samples were collected from 165 individuals. 61.2% of participants were female. † this study conducted across two identical experiments 10 months apart. ‡ this study conducted across four experiments using two cohorts – the first cohort was used for one experiment, and the second cohort was used for three.

### Categorizing Whole-Blood RNAs Based on Temporal Changes in Transcript Abundance

To assist RNA-based biomarker development, we identified groups of whole-blood RNAs with optimal temporal profiles for different types of biomarkers by following the steps outlined in **Figure 1A** and detailed in the Methods. These classifications represent operational definitions designed to identify the extremes of a continuum of temporal behavior rather than distinct biological classes. Using this approach, more than half of the transcripts in whole blood (3,448/6,064) were initially classified as trait or state before removing transcripts that met criteria for both categories. Overlapping transcripts were removed to ensure mutually exclusive gene sets for downstream analyses. The resulting trait and state gene sets used for subsequent analyses are shown in **Figure 1B**. See **Table S1** in Supplementary Data File 2 for the trait and state gene lists, or they can be downloaded from the Blood RNA Stability Atlas application (https://wcaar.shinyapps.io/blood-rna-stability-atlas/). Each category is described in detail in the following paragraphs.

**Figure 1.**
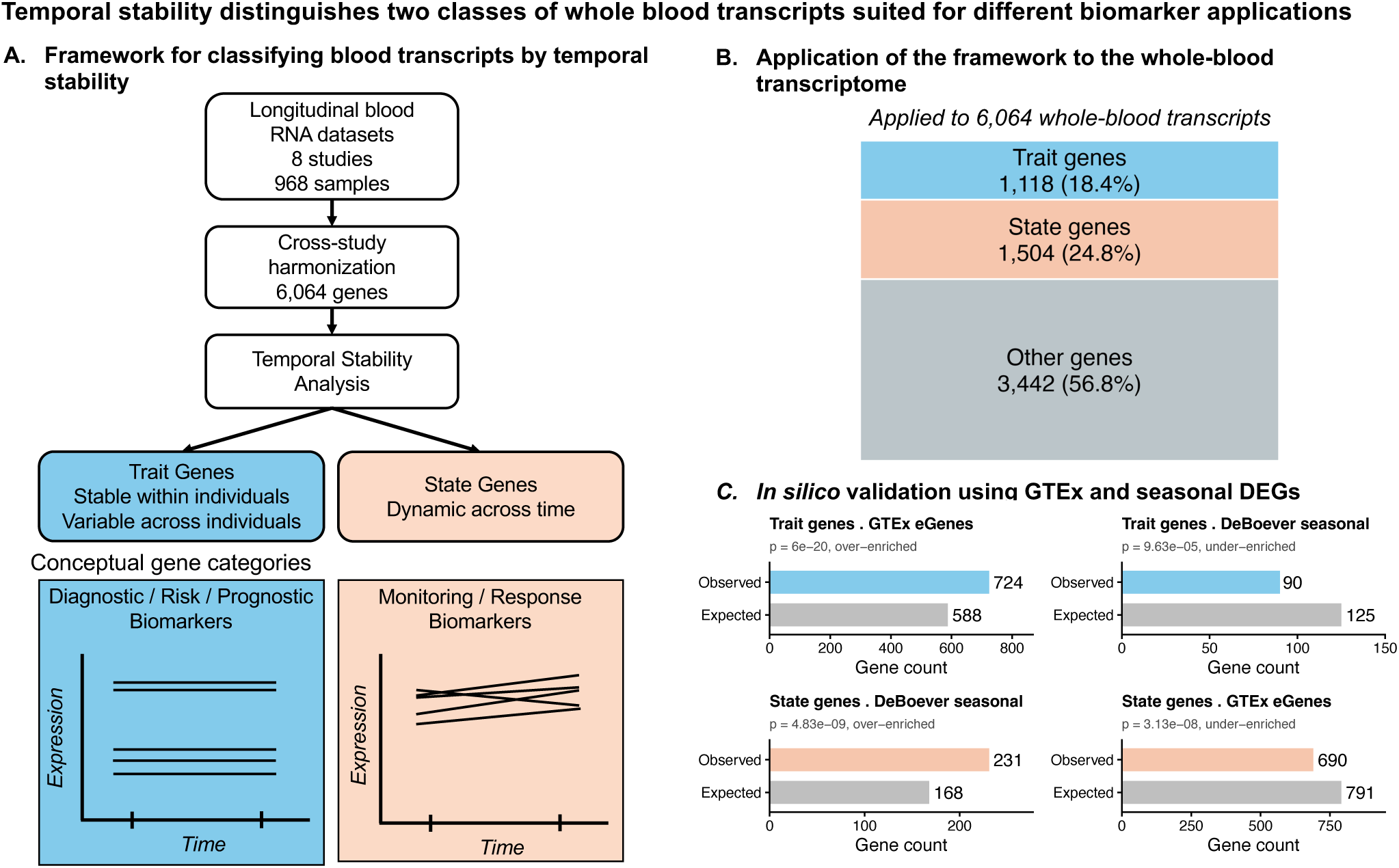
Temporal stability distinguishes two classes of whole blood transcripts suited for different biomarker applications. **A.)** Framework for classifying blood transcripts by temporal stability. Longitudinal whole blood RNA datasets (8 studies, 968 samples) were harmonized to identify 6,064 transcripts measured across studies. Temporal stability analysis classified transcripts as trait genes, which are stable within individuals but variable across individuals, or state genes, which show dynamic changes over time. Conceptual diagrams illustrate the expected temporal behavior of each class and potential biomarker applications associated with stable versus dynamic transcript behavior. **B.)** Application of the framework to the whole blood transcriptome. Among the 6,064 transcripts analyzed, 1,118 (18.4%) were classified as trait genes and 1,504 (24.8%) as state genes, while the remaining transcripts did not meet criteria for either class. **C.)** *In silico* validation analysis shows that trait and state genes align with previous findings. Trait genes are enriched for whole blood eQTL genes– overlap between trait genes and GTEx whole blood eGenes significant enrichment, while overlap with a set of genes shown to vary with season (DeBoever et al. [26]) showed significant under-enrichment of trait genes. The opposite pattern was observed for state genes. GTEx = Adult Genotype Tissue Expression; DEGs = differentially expressed genes

### Trait Genes: Transcripts with Stable Interindividual Levels

Diagnostic, predictive, prognostic, and risk biomarkers often depend on capturing stable biological differences between individuals. We therefore defined “trait genes” as transcripts exhibiting low within-individual variation across time but substantial variability between individuals. Using the thresholds described in the Methods, we identified 1,118 trait genes (**Figure 1B**). Notably, 88 transcripts exhibited trait-like behavior across all eight studies (**Supplementary Data File 2 - Table S2**). These findings indicate that a substantial fraction of blood transcripts show stable inter-individual differences in abundance levels. This suggests that trait transcripts may be less sensitive to environmental or physiological fluctuations such as sleep, exercise, diet, or vaccinations. Such stability is a property that could be advantageous for diagnostic or risk biomarker development.

Because trait genes exhibit stable transcript levels within individuals, we hypothesized that their variability would be driven in part by genetic regulation. Consistent with this expectation, trait genes were strongly enriched for significant whole blood eGenes (defined as having a Storey q-value < 0.001 for strongest eQTL) identified in the GTEx dataset (hypergeometric P = 6.0 × 10⁻²⁰; **Figure 1C**), supporting a genetic contribution to stable inter-individual differences in transcript abundance levels. Conversely, state genes, which showed variable transcript levels within individuals, were significantly under-enriched for eGenes (lower-tailed hypergeometric P = 3.1 × 10^−8^; **Figure 1C**).

To examine the biological processes represented by temporally stable transcripts, we performed functional enrichment analysis of the trait gene set. Trait genes were strongly enriched for immune and hematopoietic pathways, including innate immune response, leukocyte activation, and hematopoietic cell lineage (**Figure 2; Supplementary Data File 2 - Table S3–S4**).

**Figure 2.**
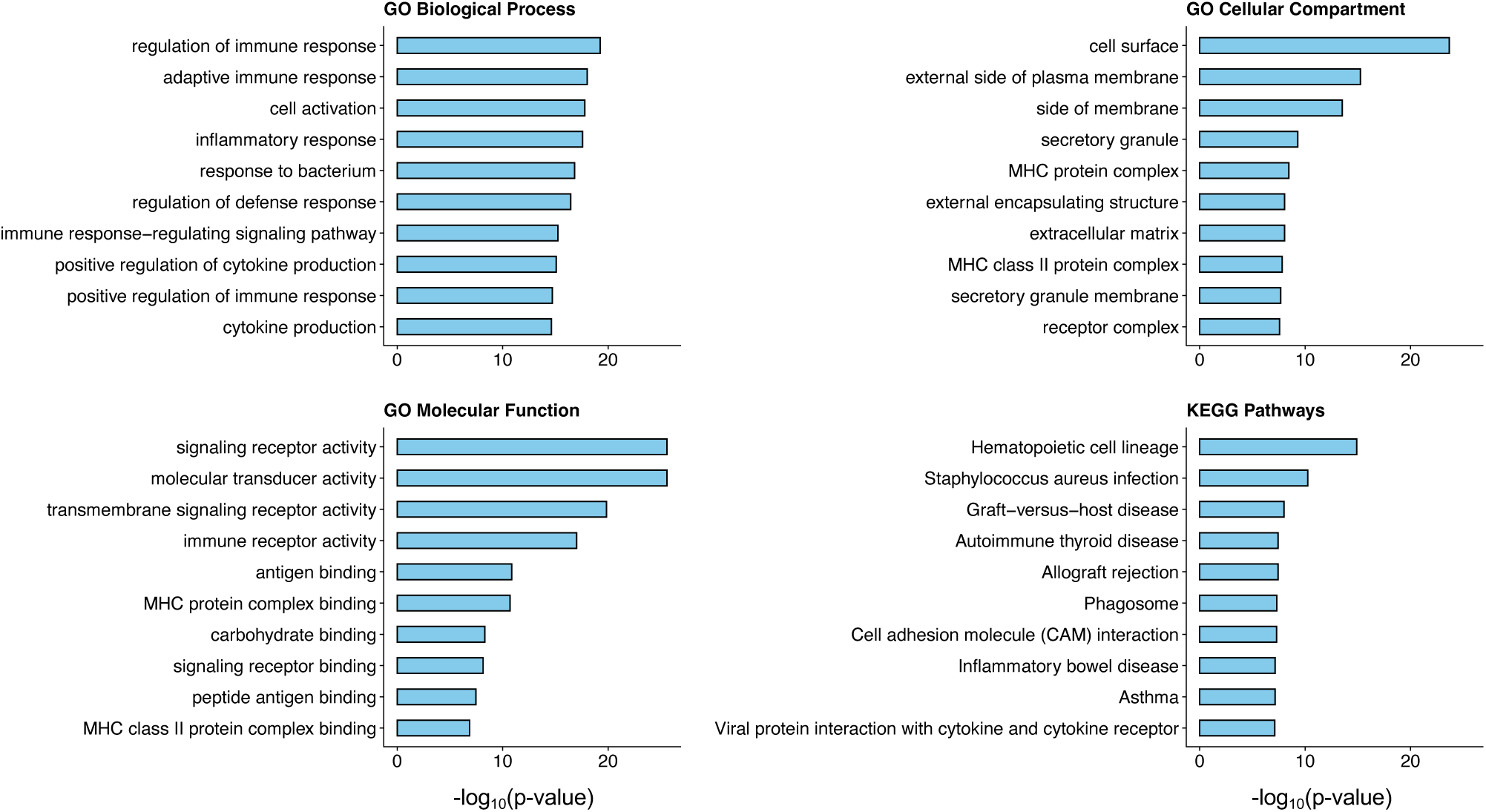
Functional enrichment analysis of trait genes. Top enriched functional categories for trait genes identified in the temporal stability analysis. Each panel shows the top ten significantly enriched terms for a given ontology. All displayed categories passed Benjamini-Hochberg multiple testing correction. Complete enrichment results for both trait and state genes are provided in **Supplementary Data File 2 - Tables S3–S4**. (GO = Gene Ontology, KEGG = Kyoto Encyclopedia of Genes and Genomes.)

Because whole blood transcript abundance reflects both transcriptional regulation and cellular composition, stable expression patterns may also capture stable inter-individual differences in leukocyte populations. Together, these findings indicate that trait-like transcripts represent reproducible, genetically influenced differences between individuals and may therefore be particularly useful for biomarker applications requiring stable measurements across time including diagnostic, predictive, and risk biomarkers.

### State Genes: Transcripts with Dynamic Temporal Modulation

Transcripts exhibiting dynamic changes in abundance across time were classified as “state genes,” reflecting temporally responsive transcriptional behavior that may be relevant for monitoring or response-oriented biomarker applications. Using the criteria described in the Methods for assessing temporal modulation, 1,504 transcripts met the threshold in at least four studies (**Figure 1B**).

In contrast to the highly reproducible trait signals, temporal transcriptional responses were more heterogeneous across datasets, consistent with context-dependent biological regulation. The number and identity of differentially expressed transcripts varied substantially across studies, and no gene was differentially expressed in all eight datasets. A list of the 44 transcripts called temporally modulated in six studies is available in **Table S5** of **Supplementary Data File 2**.

Examples of temporal responses within individual datasets, including vaccine responses and diurnal transcriptional variation, are shown in **Figures S1-S4** of **Supplementary Data File 3** and **Table S6** of **Supplementary Data File 2**.

To evaluate whether these transcripts capture previously reported temporal variation in the blood transcriptome, we compared our state gene set with transcripts reported by DeBoever et al. to exhibit temporal changes in abundance in whole blood [26]. Among the 679 DeBoever transcripts overlapping our 6,064-transcript universe, 231 were also classified as state genes, representing a significant enrichment (hypergeometric P = 4.8 × 10^−9^). Consistent with the conceptual distinction between the two classes, trait genes were significantly under-enriched in the DeBoever gene set (lower-tailed hypergeometric P = 9.6 × 10^−5^) (**Figure 1C**).

Functional enrichment analysis of the state gene set revealed five ontologies that passed multiple testing correction: translation, protein biosynthetic process, and cytoplasmic translation for GO Biological Process; mRNA binding for GO molecular function; and ribonucleoprotein complex for GO Cellular Compartment (**Figure 3; Supplementary Data File 2 - Table S7–S8**). These findings indicate that temporally dynamic transcripts can capture responsive gene expression programs that vary with environmental or experimental context. Having established these transcript classes, we next developed a resource to integrate temporal stability with genetic and disease association data and thereby facilitate biomarker prioritization.

**Figure 3.**
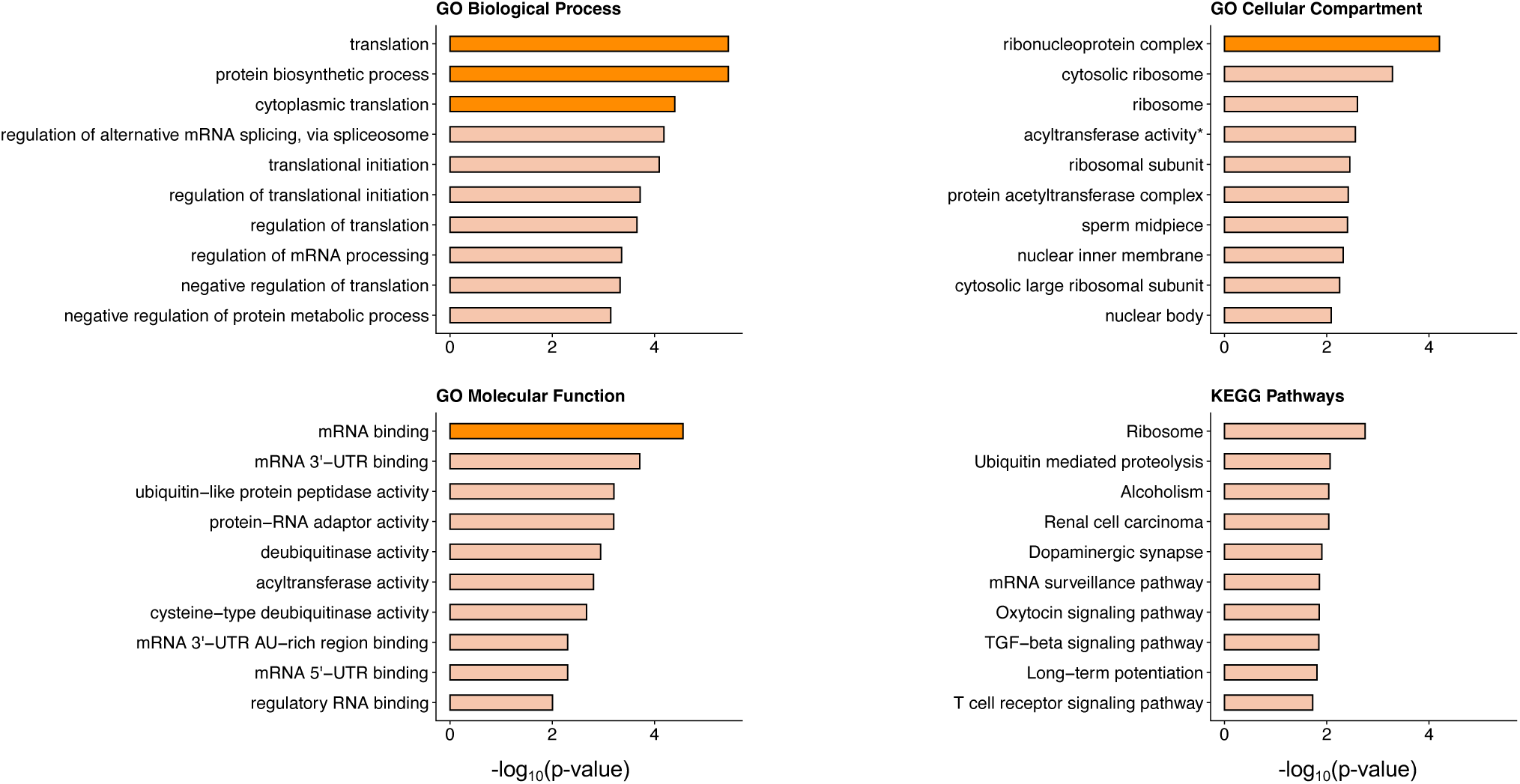
Functional enrichment analysis of state genes. Top enriched functional categories for state genes identified in the temporal stability analysis. Each panel shows the top ten significantly enriched terms for a given ontology. Categories in dark orange passed Benjamini-Hochberg multiple testing correction. Complete enrichment results for both trait and state genes are provided in **Supplementary Data File 2 - Tables S7–S8**. (GO = Gene Ontology, KEGG = Kyoto Encyclopedia of Genes and Genomes.)

### The Blood RNA Stability Atlas Integrates Temporal Stability with Genetic and Disease Associations

To enable systematic exploration of transcript stability in the context of human disease genetics, we developed the Blood RNA Stability Atlas, an interactive resource that integrates temporal stability classifications with whole blood eQTLs and GWAS associations. The Atlas allows users to download RNA stability classifications, visualize longitudinal expression patterns across the eight datasets in this study, and query disease associated genetic variants linked to whole blood expression. For example, querying individual genes in the “Investigate Blood RNA Temporal Expression” interface allows visualization of longitudinal expression trajectories and positioning of each gene within the distribution of stability metrics across all transcripts.

To extend the utility of the framework, we linked the 6,064 blood transcripts to whole-blood eQTL variants from the GTEx v11 database. In total, 4,395 blood transcripts were associated with at least one eQTL SNP. These variants were then linked to clinical phenotypes using the GWAS Catalog, resulting in 18,358 trait associations across the 4,395 genes. The application integrates these resources to connect disease-associated genetic variants with whole-blood transcript expression and temporal stability metrics. These results are displayed dynamically in the “Discover RNA Biomarker Candidates” interface, allowing users to prioritize candidate biomarkers based on genetic association, transcript stability, and expression characteristics. We next demonstrate the utility of this framework and resource by prioritizing candidate blood RNA biomarkers for coronary artery disease.

### Prioritizing Blood RNA Biomarker Candidates for Coronary Artery Disease Using the Blood RNA Stability Atlas

To demonstrate the utility of the Blood RNA Stability Atlas, we used the application to identify candidate blood RNA biomarkers for coronary artery disease (**Figure 4A**). CAD is highly heritable and extensively studied in genome wide association studies, making it an ideal test case for integrating genetic associations with transcript stability to prioritize candidate biomarkers [27].

**Figure 4.**
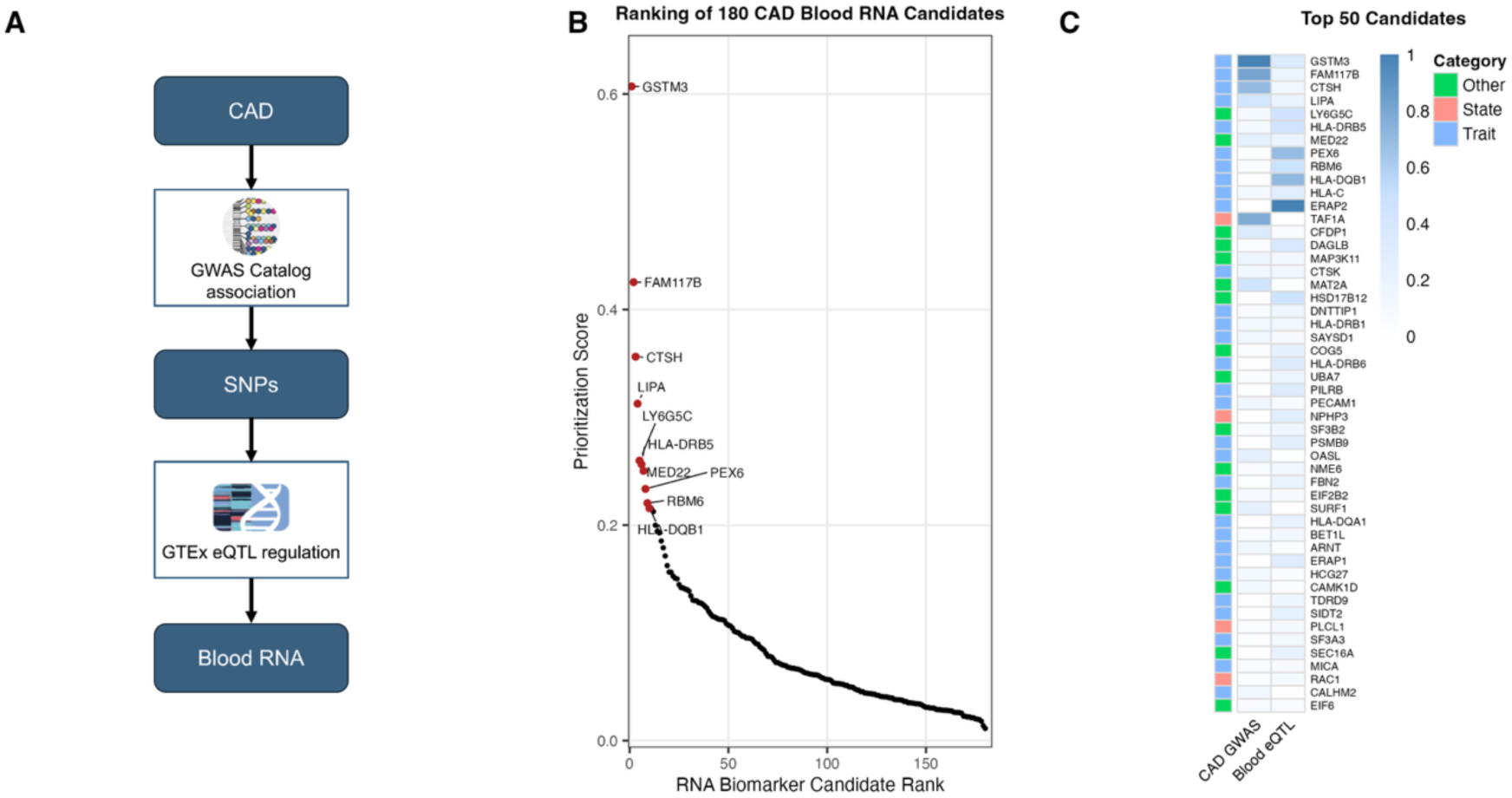
Identification and triage of candidate CAD blood RNA biomarkers using the Blood RNA Stability Atlas. **A.)** Schematic of the variant-based search framework linking SNPs associated with clinical phenotypes to blood RNA abundance levels. Candidate biomarkers for coronary artery disease (CAD) were identified by integrating GWAS associations with whole blood eQTL data. **B.)** Illustrative ranking of 180 CAD-associated blood RNAs using a composite score based on CAD GWAS association strength and blood eQTL support. The score is intended to support candidate triage rather than define a final biomarker panel. Each point represents a gene ranked by its combined score. Top 10 ranked transcripts are highlighted in red. The full list of 180 CAD-associated blood RNAs is provided in **Supplementary Data File 2 – Table S9**. **C.)** Heatmap of normalized feature values for the top 50 ranked candidates. Rows represent genes ordered by the composite score, and row annotations indicate temporal classification as trait, state, or other.

Querying the application for the GWAS trait “coronary artery disease” yielded 180 whole-blood RNAs linked to CAD-associated eQTL variants (**Supplementary Data File 2 - Table S9**). These genes represent a disease-linked candidate space rather than a definitive biomarker panel. To facilitate visualization and triage of candidates for downstream testing, we calculated an illustrative prioritization score based on CAD GWAS association strength, blood eQTL support, and expression level (**Figure 4B-C**). This score was used to rank candidates in Figure 4, but all 180 genes remain potentially relevant depending on the intended biomarker application, assay platform, and validation context.

Among the 180 candidates, 47 were classified as trait genes and 38 as state genes, demonstrating how the Atlas can annotate disease-linked candidates by temporal behavior. Trait genes were enriched among higher-ranked candidates, consistent with the representation of genetically-regulated transcripts among the disease-associated candidate set. This enrichment was statistically significant (Trait vs State and Trait vs Other, Wilcoxon P < 7 × 10^−6^), although the magnitude of the effect was moderate. Notably, three genes identified among the 13 whole blood cis-eGenes with Mendelian randomization evidence supporting causal relationships with CAD in the Framingham Heart Study [28], LIPA, CAMK1D, and NUDT5, were also present among the 180 CAD associated candidate transcripts identified here. This overlap was greater than expected by chance even when using all 6,064 analyzed transcripts as a conservative background set (hypergeometric P = 0.0059), providing orthogonal support that the Atlas can prioritize biologically relevant disease-associated blood transcripts.

This example illustrates how the Atlas can help investigators move from broad disease associated genetic signals to a refined set of blood RNA candidates with interpretable temporal stability and expression characteristics. As a complementary use case, we next demonstrate how the Atlas can be used to contextualize candidate biomarkers identified in independent studies.

### Application of the Blood RNA Stability Atlas to Published Sepsis Biomarkers

To demonstrate how the Blood RNA Stability Atlas can be used to interpret and contextualize candidate biomarkers, we examined two genes, BPGM and TAP2, identified as part of a minimal classifier for a high-risk sepsis endotype in a large whole blood transcriptomic study of critically ill patients [29]. In this study, transcript abundance ratios involving BPGM and TAP2 were used to identify patients belonging to the Mars1 endotype, which was consistently associated with increased mortality across cohorts.

Using the Blood RNA Stability Atlas, BPGM was classified as a trait gene and TAP2 was not classified as a trait or state gene although stability metrics indicated relatively high stability for TAP2 which had a median repeatability score in the 87^th^ percentile. Visualization of transcript abundance levels across a 24-hour time course showed minimal temporal fluctuation for both transcripts, supporting the conclusion that their abundance levels are robust to sampling time (**Figure 5A**). This property is particularly relevant in acute clinical settings such as ICU admission, where biomarker measurements must be reliable despite variability in collection timing.

**Figure 5.**
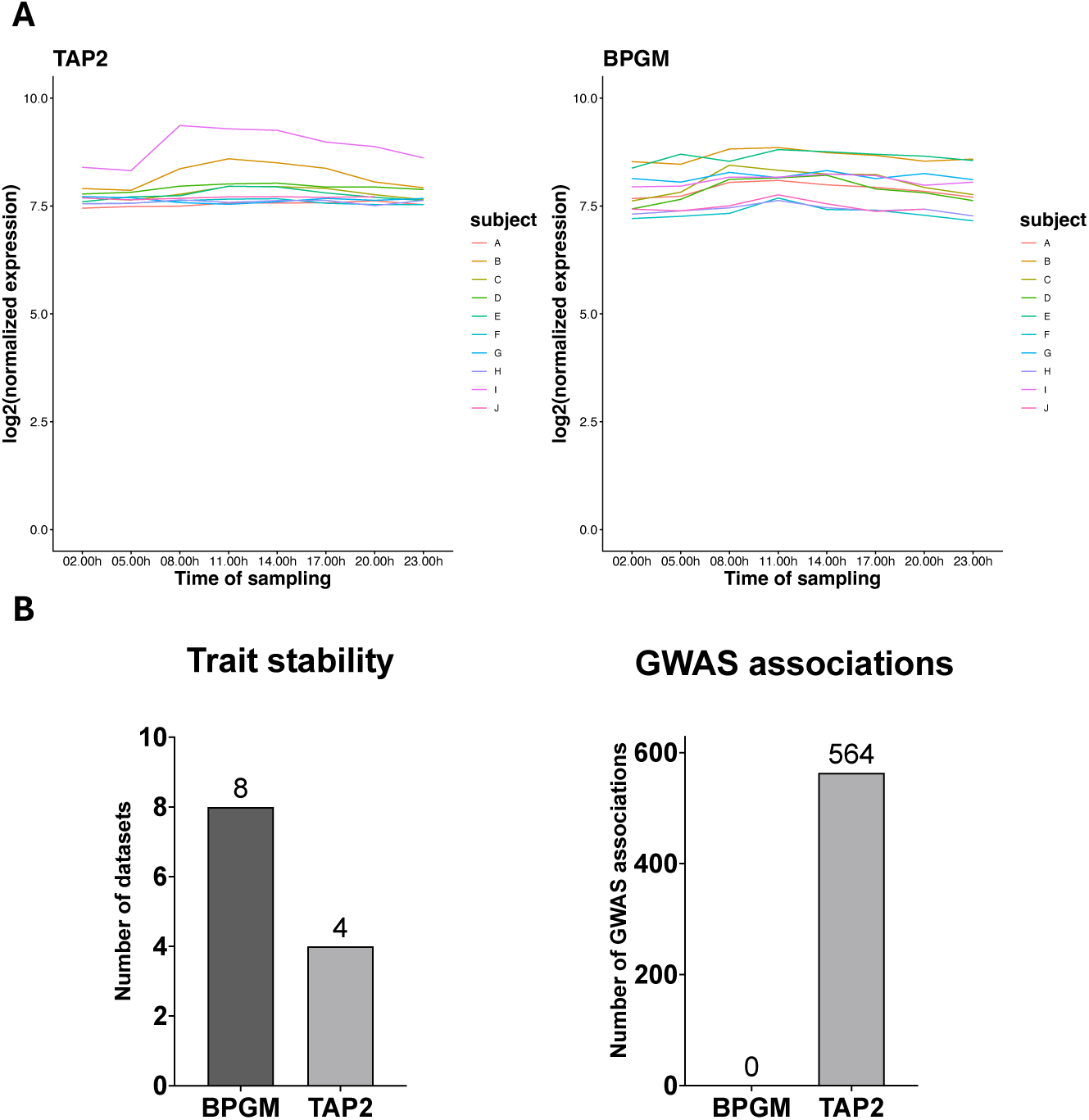
Temporal stability and genetic context of published sepsis biomarkers using the Blood RNA Stability Atlas. **A.)** Transcript abundance measurements of BPGM and TAP2 across a 24-hour time course [20] demonstrates minimal within-subject variation, indicating high temporal stability for both transcripts. **B.)** Number of datasets in which each transcript exhibits trait-like patterns and the number of GWAS trait associations linked to their blood eQTLs. BPGM exhibits strong trait-like behavior across all datasets but lacks known GWAS associations, whereas TAP2 shows slightly lower trait consistency and is associated with numerous disease-linked variants. These results illustrate how the Blood RNA Stability Atlas distinguishes between stable transcripts with different biological interpretations.

Despite shared temporal stability, BPGM and TAP2 differed markedly in their genetic and disease association profiles (**Figure 5B**). TAP2 transcript abundance was linked to multiple blood eQTLs that are associated with diverse immune related traits and diseases, providing a potential connection to antigen presentation pathways and immune function. In contrast, BPGM eQTLs showed no evidence of disease association in the Atlas, suggesting that its role in the classifier may reflect stable physiological variation rather than genetically mediated disease risk.

These differences illustrate how the Atlas can distinguish between candidate biomarkers that are similarly stable but differ in their potential biological interpretation and downstream utility.

Together, this example demonstrates how integrating temporal stability with genetic and disease association data can provide additional context for interpreting existing biomarker candidates, guiding prioritization and hypothesis generation for follow up studies.

## Discussion

Whole blood RNA biomarkers have been widely investigated for diagnosis, prognosis, risk prediction, and disease monitoring. However, most studies focus on associations between transcript abundance and phenotype, often using cross-sectional designs or longitudinal disease cohorts without corresponding evaluation of temporal behavior in healthy individuals. As a result, the temporal stability of candidate blood RNA biomarkers is rarely incorporated into biomarker prioritization and interpretation. Here, we address this gap by quantifying transcript stability across eight longitudinal healthy cohorts spanning diverse biological contexts, integrating these measurements with genetic regulation and disease associations, and making the resulting framework publicly accessible through the Blood RNA Stability Atlas. Our cross-platform consensus identified the immune-related stability and genetic-regulation patterns reported in previous longitudinal studies, demonstrating that these signatures generalize across heterogeneous study designs rather than reflecting artifacts of individual datasets [14–17].

Although previous longitudinal studies established that blood gene expression contains both stable and dynamic components, our principal contribution is to provide the first cross-platform consensus framework for transcript-level temporal stability, integrating diverse longitudinal datasets into a practical resource for biomarker discovery and interpretation.

Trait transcripts exhibit high within-individual stability alongside between-individual variability, consistent with a role in capturing baseline differences that are relatively insensitive to transient physiological or environmental fluctuations. Consistent with this interpretation, trait genes were strongly enriched for whole-blood eGenes, suggesting that their expression variability is likely influenced in part by inherited genetic differences. Similar enrichment of longitudinally stable transcripts for regulatory genetic variants has been reported in both platelet transcriptomes [16] and large population-based blood studies [17], supporting the conclusion that stable expression patterns frequently reflect genetically regulated transcriptional programs. Notably, ERAP2, PAM, and HLA-DQB1 exhibited strong trait-like behavior across datasets. ERAP2 is particularly illustrative because its expression is known to be highly genotype dependent [30], providing an independent example linking temporal stability with inherited regulation. Together, these findings suggest that trait transcripts represent reproducible biological programs that are well suited for applications requiring consistent measurements across time, including diagnostic, prognostic, and risk biomarkers.

One of the most striking observations was the strong enrichment of trait genes for immune and inflammatory pathways. Immune-related transcripts are often presumed to be poor candidates for diagnostic biomarkers because their expression can fluctuate in response to infection or environmental perturbation. However, despite integrating datasets that included immune stimulation, such as vaccination, many immune-related transcripts remained remarkably stable within individuals. These findings extend previous observations that immune and interferon-signaling genes are among the most temporally stable components of the blood transcriptome [15] and that T- and B-cell-associated transcripts form stable individual-specific expression signatures [17]. By reproducing these patterns across independent cohorts, measurement platforms, and sampling intervals ranging from less than one hour to several months, our results demonstrate that this immune-centered stable core represents a robust feature of healthy blood biology rather than a study-specific observation. The persistence of these stable immune transcriptional programs suggests that baseline immune state may reflect enduring biological characteristics despite ongoing physiological and environmental challenges, reinforcing their potential value as robust blood biomarkers.

In contrast to trait transcripts, state transcripts exhibited substantially greater temporal variability and were less consistently reproduced across the heterogeneous datasets analyzed here.

Rather than representing a single biological program, these transcripts likely reflect diverse physiological responses to environmental, behavioral, and technical influences that differ across cohorts. Nevertheless, state genes were enriched for pathways related to translation, mRNA binding, ribonucleoprotein complexes, and other components of gene-expression machinery, consistent with recent population-scale analyses showing that ribosomal proteins, RNA processing, and translational pathways are among the most temporally dynamic components of the blood transcriptome [17]. Although their greater temporal variability makes state transcripts less suitable for applications requiring stable baseline measurements, these dynamic genes are likely to be particularly informative in settings where transcriptomic responses reflect disease progression, treatment response, or environmental exposure. For example, approaches that aim to model transcriptomic dynamics or predict disease trajectories from single timepoint measurements require careful gene selection, as many blood transcripts do not vary with disease state and may obscure informative signals [31]. Rather than representing undesirable biological noise, temporal responsiveness may itself constitute the biological signal of interest for many monitoring and pharmacodynamic biomarkers.

The distinction between trait and state transcripts has important implications for biomarker development. However, these categories should not be viewed as competing biomarker classes or as a universal feature-selection criterion. Rather, they represent complementary sources of biological information along a continuum of temporal stability. In this study, transcripts were classified as trait or state by identifying the extremes of this continuum, but many transcripts outside these categories exhibit intermediate stability and may represent valuable biomarkers in specific contexts. Stable transcripts may provide robust measurements of baseline biological state, whereas dynamic transcripts may capture disease activity, treatment response, or environmental perturbation. Future biomarker panels may therefore benefit from integrating both classes, combining stable baseline markers with disease-responsive transcripts to improve diagnostic performance and biological interpretability. The Blood RNA Stability Atlas enables investigators to explore this continuum directly by evaluating transcript stability alongside disease relevance and other biological features. Importantly, temporal stability represents one dimension of biomarker robustness and complements, rather than replaces, careful validation across independent cohorts, populations, and measurement platforms.

Beyond providing transcript classifications, the Blood RNA Stability Atlas is intended as a practical framework for biomarker discovery and interpretation. We envision two complementary workflows. In a top-down approach, investigators begin with a disease or trait of interest and identify candidate blood RNA biomarkers linked to disease-associated genetic variants.

Conversely, in a bottom-up approach, investigators can evaluate candidate transcripts emerging from experimental studies in the context of their temporal stability, longitudinal expression behavior, and associated genetic evidence. Together, these complementary strategies support feature prioritization, interpretation of cross-study variability, study design, and hypothesis generation. Although the Atlas integrates blood RNA measurements with eQTL and GWAS associations, these relationships should be considered hypothesis-generating rather than causal. We did not perform colocalization or causal inference analyses and therefore do not interpret shared genetic associations as evidence that blood RNA abundance mediates disease risk. Instead, genetic associations are intended to provide complementary biological context for prioritizing candidate biomarkers, with experimental validation remaining essential for establishing causal mechanisms [32, 33].

A major source of variation in whole-blood transcriptomes is cellular composition [14, 34]. Consequently, a portion of the temporal variability observed here likely reflects fluctuations in circulating leukocyte populations in addition to transcriptional regulation within individual cell types. Previous studies have shown that changes in leukocyte abundance account for a substantial fraction of transcriptomic variation in blood biomarkers [35]. However, recent population-scale analyses demonstrated that although cell composition contributes to gene-expression variability, correcting for estimated leukocyte proportions has minimal impact on the overall ranking of temporally stable versus dynamic transcripts [17], indicating that compositional changes alone do not explain the temporal structure of the healthy blood transcriptome. Importantly, we intentionally evaluated stability in whole blood without correcting for cell composition because clinically deployed blood RNA assays are typically performed on bulk samples rather than cell-type-resolved preparations. Accordingly, the stability metrics presented here reflect the behavior of transcripts under conditions that most closely resemble real-world biomarker applications.

Several limitations define the scope of this framework. First, our analyses were restricted to healthy populations, and transcript stability may differ in disease states where both baseline expression and temporal dynamics are altered. Second, the datasets analyzed here were collected retrospectively and differ in cohort characteristics, sampling protocols, and measurement platforms. Although this heterogeneity strengthened our ability to identify transcripts exhibiting reproducible temporal behavior across diverse biological and technical settings, it also limited standardized control of experimental variables. Finally, our analyses were performed at the gene level and therefore do not capture transcript isoform dynamics. Recent work has shown that differential transcript usage contributes substantially to intra-individual variation, particularly among temporally dynamic genes [17], highlighting an additional layer of temporal regulation not addressed by the present framework.

This study establishes a framework that can expand as additional longitudinal transcriptomic datasets become available. Future studies using standardized longitudinal sampling across multiple timescales within the same individuals will help determine whether short-term stability predicts long-term stability or whether transcript stability itself depends on sampling interval.

Extending this framework to disease cohorts will be particularly important for determining whether transcripts identified as stable in healthy individuals retain their measurement robustness under pathological conditions. Future integration of transcript-level quantification, matched leukocyte measurements, and single-cell transcriptomics will further distinguish stable transcriptional programs from changes driven by cellular composition or isoform usage. As these resources continue to grow, they can be incorporated directly into the Blood RNA Stability Atlas, allowing the framework to evolve alongside advances in longitudinal transcriptomics and biomarker discovery.

## Conclusion

We identify temporal stability as a defining property of blood RNA expression and show that transcripts span a continuum that can be approximated as stable “trait” and dynamic “state” genes. This distinction provides a framework for interpreting blood transcriptomic variation by distinguishing stable baseline abundance levels from context-dependent responses. The Blood RNA Stability Atlas extends this framework by integrating transcript stability with genetic and disease-associated variants, enabling temporal patterns to be interpreted in the context of disease genetics to support candidate prioritization and hypothesis generation. More broadly, this work helps operationalize temporal variation as a practical dimension of biomarker discovery, transforming a long-recognized but underutilized concept into an accessible resource for transcriptomic research.

## Methods

Detailed preprocessing workflows, quality control procedures, statistical model specifications, and software packages and versions are described in the **Supplementary Methods**.

### Ethical Considerations

Each study acquired the consent of those involved in blood RNA experiments as well as the approval of an independent review board, as described in the study methodologies.

### Study Selection

We searched Google Scholar and the Gene Expression Omnibus for publicly available longitudinal whole blood transcriptomic datasets derived from human participants. Searches were performed using combinations of terms related to whole blood, longitudinal sampling, and gene expression. Studies were included if they sampled individuals at multiple timepoints under generally healthy conditions and provided sufficient metadata for longitudinal analysis.

Healthy populations were defined as individuals without diagnosed disease and not exposed to sustained or severe interventions expected to substantially alter systemic physiology. Studies involving acute or moderate environmental or behavioral perturbations (for example, vaccination, exercise, or sleep variation) were retained to capture typical sources of physiological variability. Studies and individual samples failing quality control criteria were excluded as described below.

### Data Processing

All datasets were processed from raw or count level data using Bioconductor workflows in RStudio. Microarray data from Affymetrix and Illumina platforms were background corrected, normalized, and filtered for expression using established affy and limma pipelines, including RMA [36] for Affymetrix arrays and NEQC [37] for Illumina arrays. Probes were retained if they were detected above background in a minimum number of samples defined by the smallest timepoint group within each study.

RNA sequencing count matrices were normalized using the trimmed mean of M values method using the edgeR package [38, 39] and filtered using the filterByExpr function, which requires genes to have greater than 10/median(CPM) in at least the smallest number of samples collected for any of the timepoints in an experiment. Precision weights were estimated using voom to account for the mean variance relationship prior to downstream modeling [40].

Quality control included evaluation of array level metrics, multidimensional scaling, and principal component analysis. Samples that failed to cluster with other samples from the same individual across timepoints were excluded. A summary of quality control procedures is provided in **Supplementary Data File 1**.

Gene identifiers were harmonized across platforms using the mapIds function from the AnnotationDbi package with associated database objects for mapping gene symbols [41]. Analyses were restricted to the 6,064 genes present across all datasets after filtering.

### Changes in Transcript Abundance over Time

Changes in transcript abundance across timepoints were evaluated using linear models implemented with limma [42], with time modeled as a fixed effect and repeated measures within individuals accounted for using duplicateCorrelation. Precision weights calculated using voom were incorporated into models built on RNA-seq data. Covariates such as age, sex, race, and ethnicity were included where available. Nominal p-values (unadjusted P < 0.05) were used to identify transcripts exhibiting temporal modulation, prioritizing sensitivity for downstream cross-study consensus rather than significance within individual datasets.

### Quantifying Temporal Stability and Defining Transcript Classes Trait RNAs

We hypothesized that ideal diagnostic, predictive, prognostic, and risk biomarkers would exhibit variability between individuals but stability within an individual over time. Transcripts exhibiting this pattern were classified as trait RNAs. To identify trait RNAs, we calculated complementary statistics for each transcript or probe using the processed data for each study as described in the following paragraph. We then used these statistics to create thresholds for longitudinal stability and interindividual variability.

For each transcript within each study, we quantified temporal stability and inter-individual variability using complementary statistics: within-individual standard deviation (σ_w_), total standard deviation (σ_t_), variance partitioning for subject (v_s_) and time effects (v_t_) using the variancePartition package [43], repeatability (r), and estimated genetic variance (v_g_) using the heritability package [44]. Repeatability, also called the intraclass correlation coefficient, is proportional to the consistency of a transcript’s measurement for a specific individual across multiple timepoints and is a complementary statistic to v_s_. The genetic variance, v_g_, estimates the variation in expression of a transcript due to an individual’s genetics.

Trait RNAs were defined using percentile-based thresholds selected to capture transcripts with relatively low within-individual variability and high between-individual variability within each study. The thresholds for trait RNAs were based on three pairs of statistics and calculated for each transcript per study: 1) σ_w_ less than or equal to the 25^th^ percentile and σ_t_ greater than or equal to the 75^th^ percentile, 2) v_s_ greater than or equal to the 75^th^ percentile and a v_t_ less than or equal to the 25^th^ percentile, 3) r and v_g_ greater than the 75^th^ percentile. These thresholds were applied to identify reproducible extremes of temporal stability within each dataset rather than to define universal cutoffs, enabling consistent cross-study comparison across heterogeneous cohorts and platforms.

### State RNAs

State RNAs were defined as transcripts exhibiting significant temporal modulation (unadjusted P < 0.05). For each study, differential expression across timepoints was assessed as described above, and transcripts were ranked by frequency of temporal association across datasets.

### Cross-Study Analysis

To define robust transcript categories, we performed a cross-study consensus analysis across all datasets. Transcripts were classified as trait or state based on their longitudinal abundance patterns within each study and then aggregated across studies. To prioritize reproducibility across heterogeneous cohorts and platforms, transcripts were required to meet category criteria in at least four independent studies, including representation in both microarray and RNA sequencing datasets. We selected this threshold to prioritize reproducibility across platforms while retaining adequate sensitivity to identify broadly stable or dynamic transcripts. Transcripts meeting criteria for both categories were excluded to ensure non overlapping sets.

### *In Silico* Validation

To evaluate the biological relevance of transcript categories, we tested for enrichment of trait and state gene sets in independent reference datasets. Trait genes were tested for overrepresentation among whole blood expression quantitative trait loci associated genes from GTEx [45] using genes with Storey q values less than 0.001. State genes were tested for overrepresentation among previously reported temporally variable blood transcripts [26]. Enrichment was assessed using one tailed hypergeometric tests.

### Functional Enrichment Analysis

To gain biological insight into the RNA categories we defined, we used the clusterProfiler enrichGO and enrichKEGG functions with the universe argument set to the 6,064 common gene symbols to conduct an overrepresentation analysis for Gene Ontology (GO) categories and KEGG pathways in the sets of RNAs making up each category using the one-tailed hypergeometric test [46].

### Blood RNA Stability Atlas

To enable exploration of transcript stability metrics derived from this study, we developed the Blood RNA Stability Atlas, an interactive web application built using the R Shiny framework. The application integrates gene level stability statistics, cross study category assignments, and associated annotations, including whole blood eQTLs from GTEx and disease associations from the GWAS Catalog. It allows users to query by individual genes or disease/phenotype of interest, visualize longitudinal expression patterns across studies, and download processed data.

All metrics and classifications presented in the application are derived directly from the analyses described above. The application, source data, and user documentation are publicly available at: https://wcaar.shinyapps.io/blood-rna-stability-atlas/ and https://github.com/wbaltazar/blood-rna-stability-atlas, and will be updated as additional datasets are incorporated.

### Coronary Artery Disease Example: Composite Biomarker Prioritization Score

To triage candidate biomarkers for coronary artery disease (CAD) identified through the Blood RNA Stability Atlas and aid in visualization, we developed a composite scoring framework integrating multiple biologically relevant features. For each transcript, two components were considered: (i) strength of association with CAD based on GWAS (-log10 P value) and (ii) strength of blood eQTL association (-log10 P value). Each component was rescaled to a 0–1 range using min-max normalization. To integrate these features into a single prioritization metric, we calculated the geometric mean of the rescaled values for each transcript. This approach emphasizes candidates with consistently strong support across all features while penalizing transcripts that perform poorly in any individual dimension. Transcripts were ranked based on the composite prioritization score. For visualization purposes, feature contributions for the top-ranked transcripts were displayed using a heatmap of normalized component scores.

### Sepsis Biomarker Example

To demonstrate application of the Atlas to externally derived candidate biomarkers, we evaluated transcripts identified in a published whole-blood transcriptomic study of sepsis that defined molecular endotypes associated with clinical outcomes [29]. Two transcripts from a minimal classifier for the high-risk endotype (BPGM and TAP2) were selected as representative examples and queried in the application. Temporal stability metrics, including repeatability and within-subject variation, longitudinal expression patterns across datasets, and associated whole-blood eQTL and GWAS annotations were evaluated using the Atlas interface. Trait classification and the number of datasets exhibiting trait-like temporal patterns were also examined to assess consistency of temporal behavior across studies.

## Supporting information

Datasets & Quality Control

## List of Abbreviations

BEST: Biomarkers, EndpointS, and other Tools
CAD: coronary artery disease
DEG: differentially expressed gene
eGene: a gene whose expression is associated with an
eQTL eQTL: expression quantitative trait locus
GO: Molecular Signature Database’s Gene Ontology
GTEx: Adult Genotype Tissue Expression (GTEx) Project
GWAS: genome-wide association study
KEGG: Kyoto Encyclopedia of Genes and Genomes
RNA: ribonucleic acid

## Declarations

### Ethics approval and consent to participate

The data collected and analyzed in this study is publicly available. Each dataset’s respective authors collected consent for each study participant’s participation and received ethical approval of their protocols from their institution’s review board. Specific details may be found in each dataset’s methodology section.

### Additional Information

The authors declare that they have no competing interests.

### Author Contributions

WCB conducted data collection, bioinformatic analyses, R Shiny code, and manuscript - original draft, review, editing. ROM was responsible for application testing and manuscript - review, editing. LBF was responsible for conceptualization, study design, bioinformatic analyses, application testing, and manuscript – draft, review, editing.

## Acknowledgements

We would like to thank the participants and researchers involved in generating the datasets that were analyzed in this study.

## Data Availability

The datasets analyzed during the current study are available in the GEO database (https://www.ncbi.nlm.nih.gov/geo/), the GTEx Portal (https://gtexportal.org/), and the GWAS Catalog (https://www.ebi.ac.uk/gwas/). These data were derived from the following resources available in the public domain: GEO accession numbers GSE239282, GSE14844, GSE223613, GSE30101, GSE10041, GSE16028, GSE81761, and GSE206505; GTEx dataset version 11; and GWAS Catalog All Associations v1.0, downloaded May 13, 2026. Additionally, the code and results used in this study are openly available on GitHub at https://github.com/wbaltazar/blood-rna-stability-atlas and the Blood RNA Stability Atlas application at https://wcaar.shinyapps.io/blood-rna-stability-atlas/.

## References

1. Hassan, M., et al., Innovations in Genomics and Big Data Analytics for Personalized Medicine and Health Care: A Review. Int J Mol Sci, 2022. 23(9).

2. Yurkovich, J.T., et al., The transition from genomics to phenomics in personalized population health. Nat Rev Genet, 2024. 25(4): p. 286–302.

3. Rosati, D., et al., Differential gene expression analysis pipelines and bioinformatic tools for the identification of specific biomarkers: A review. Comput Struct Biotechnol J, 2024. 23: p. 1154–1168.

4. Tebani, A., et al., Omics-Based Strategies in Precision Medicine: Toward a Paradigm Shift in Inborn Errors of Metabolism Investigations. Int J Mol Sci, 2016. 17(9).

5. in BEST (Biomarkers, EndpointS, and other Tools) Resource. 2016: Silver Spring (MD) Bethesda (MD).

6. Califf, R.M., Biomarker definitions and their applications. Exp Biol Med (Maywood), 2018. 243(3): p. 213–221.

7. Klein, A., et al., The Biomarkers Consortium: 20 years of advancing precision medicine. Nat Rev Drug Discov, 2026. 25(3): p. 155–156.

8. Mohr, S. and C.C. Liew, The peripheral-blood transcriptome: new insights into disease and risk assessment. Trends Mol Med, 2007. 13(10): p. 422–32.

9. Hoffmann, M., et al., Blood transcriptomics analysis offers insights into variant-specific immune response to SARS-CoV-2. Sci Rep, 2024. 14(1): p. 2808.

10. Balnis, J., et al., Peripheral Blood Omics and Other Multiplex-based Systems in Pulmonary and Critical Care Medicine. Am J Respir Cell Mol Biol, 2023. 69(4): p. 383–390.

11. Vanhaverbeke, M. and B. Izzi, Peripheral blood DNA and RNA biomarkers of cardiovascular disease in clinical practice, in Epigenetics in Cardiovascular Disease, Y. Devaux and E.L. Robinson, Editors. 2021, Academic Press: Translational Epigenetics. p. 261–281.

12. Zhu, H., et al., PCR past, present and future. Biotechniques, 2020. 69(4): p. 317–325.

13. Okeke, I.N. and C. Ihekweazu, The importance of molecular diagnostics for infectious diseases in low-resource settings. Nat Rev Microbiol, 2021. 19(9): p. 547–548.

14. Whitney, A.R., et al., Individuality and variation in gene expression patterns in human blood. Proc Natl Acad Sci U S A, 2003. 100(4): p. 1896–901.

15. Sakai, Y., et al., Eight-year longitudinal study of whole blood gene expression profiles in individuals undergoing long-term medical follow-up. Sci Rep, 2021. 11(1): p. 16564.

16. Rondina, M.T., et al., Longitudinal RNA-Seq Analysis of the Repeatability of Gene Expression and Splicing in Human Platelets Identifies a Platelet SELP Splice QTL. Circ Res, 2020. 126(4): p. 501–516.

17. Mishra, N., et al., Large-scale analysis of temporal gene expression variation in peripheral blood. Nat Commun, 2026.

18. Gomez-Carballa, A., et al., Music compensates for altered gene expression in age-related cognitive disorders. Sci Rep, 2023. 13(1): p. 21259.

19. Meaburn, E.L., et al., Assessing individual differences in genome-wide gene expression in human whole blood: reliability over four hours and stability over 10 months. Twin Res Hum Genet, 2009. 12(4): p. 372–80.

20. Gosch, A., A. Bhardwaj, and C. Courts, TrACES of time: Transcriptomic analyses for the contextualization of evidential stains - Identification of RNA markers for estimating time-of-day of bloodstain deposition. Forensic Sci Int Genet, 2023. 67: p. 102915.

21. Obermoser, G., et al., Systems scale interactive exploration reveals quantitative and qualitative differences in response to influenza and pneumococcal vaccines. Immunity, 2013. 38(4): p. 831–44.

22. Dusek, J.A., et al., Genomic counter-stress changes induced by the relaxation response. PLoS One, 2008. 3(7): p. e2576.

23. Rusch, H.L., et al., Gene expression differences in PTSD are uniquely related to the intrusion symptom cluster: A transcriptome-wide analysis in military service members. Brain Behav Immun, 2019. 80: p. 904–908.

24. LaRocca, T.J., et al., Novel whole blood transcriptome signatures of changes in maximal aerobic capacity in response to endurance exercise training in healthy women. Physiol Genomics, 2023. 55(8): p. 338–344.

25. Karlovich, C., et al., A longitudinal study of gene expression in healthy individuals. BMC Med Genomics, 2009. 2: p. 33.

26. De Boever, P., et al., Characterization of the peripheral blood transcriptome in a repeated measures design using a panel of healthy individuals. Genomics, 2014. 103(1): p. 31–9.

27. Schunkert, H., P. Natarajan, and N.J. Samani, The Inherited Basis of Coronary Artery Disease. N Engl J Med, 2026. 394(6): p. 576–587.

28. Liu, C., et al., Whole genome DNA and RNA sequencing of whole blood elucidates the genetic architecture of gene expression underlying a wide range of diseases. Sci Rep, 2022. 12(1): p. 20167.

29. Scicluna, B.P., et al., Classification of patients with sepsis according to blood genomic endotype: a prospective cohort study. Lancet Respir Med, 2017. 5(10): p. 816–826.

30. Gamazon, E.R., et al., A gene-based association method for mapping traits using reference transcriptome data. Nat Genet, 2015. 47(9): p. 1091–8.

31. Dunican, C., et al., Predicting trajectories of illness using RNA velocity of whole blood. Nat Commun, 2026. 17(1).

32. Umans, B.D., A. Battle, and Y. Gilad, Where Are the Disease-Associated eQTLs? Trends Genet, 2021. 37(2): p. 109–124.

33. Liu, B., et al., Abundant associations with gene expression complicate GWAS follow-up. Nat Genet, 2019. 51(5): p. 768–769.

34. Chomczynski, P., et al., Inter-Individual Differences in RNA Levels in Human Peripheral Blood. PLoS One, 2016. 11(2): p. e0148260.

35. O’Connell, G.C., et al., Shifts in Leukocyte Counts Drive the Differential Expression of Transcriptional Stroke Biomarkers in Whole Blood. Transl Stroke Res, 2019. 10(1): p. 26–35.

36. Gautier, L., et al., affy--analysis of Affymetrix GeneChip data at the probe level. Bioinformatics, 2004. 20(3): p. 307–15.

37. Shi, W., A. Oshlack, and G.K. Smyth, Optimizing the noise versus bias trade-off for Illumina whole genome expression BeadChips. Nucleic Acids Res, 2010. 38(22): p. e204.

38. Chen, Y., A.T. Lun, and G.K. Smyth, From reads to genes to pathways: differential expression analysis of RNA-Seq experiments using Rsubread and the edgeR quasi-likelihood pipeline. F1000Res, 2016. 5: p. 1438.

39. Robinson, M.D. and A. Oshlack, A scaling normalization method for differential expression analysis of RNA-seq data. Genome Biol, 2010. 11(3): p. R25.

40. Law, C.W., et al., voom: Precision weights unlock linear model analysis tools for RNA-seq read counts. Genome Biol, 2014. 15(2): p. R29.

41. Pagès, H., et al. AnnotationDbi: Manipulation of SQLite-based annotations in Bioconductor. 2024 [cited 2024 Jul 9]; Available from: https://bioconductor.org/packages/AnnotationDbi.

42. Phipson, B., et al., Robust Hyperparameter Estimation Protects against Hypervariable Genes and Improves Power to Detect Differential Expression. Ann Appl Stat, 2016. 10(2): p. 946–963.

43. Hoffman, G.E. and E.E. Schadt, variancePartition: interpreting drivers of variation in complex gene expression studies. BMC Bioinformatics, 2016. 17(1): p. 483.

44. Kruijer, W., et al., Marker-based estimation of heritability in immortal populations. Genetics, 2015. 199(2): p. 379–98.

45. Consortium, G.T., The GTEx Consortium atlas of genetic regulatory effects across human tissues. Science, 2020. 369(6509): p. 1318–1330.

46. Wu, T., et al., clusterProfiler 4.0: A universal enrichment tool for interpreting omics data. Innovation (Camb), 2021. 2(3): p. 100141.

