## Supplementary material for "The Blood RNA Stability Atlas: defining temporal structure and trait-state programs in the human whole-blood transcriptome": Datasets & Quality Control

The University of Texas at Austin
Waggoner Center for
Alcohol & Addiction Research

### Datasets & Quality Control

Whole blood datasets used for stability and variability analysis

Will Baltazar

### General Overview

Summary plots were generated for each dataset according to the platform they were on. For each dataset, normalization and gene filters were applied as described in the Methods. Then, Principal Component Analysis was used to visualize variation among samples. Samples which had faulty metrics as measured by summary statistics or deviated significantly from other samples in PCA after visual inspection were excluded from analysis.

This PowerPoint includes all plots and visualizations used for our quality control analysis and describes which samples from each dataset were removed from the analysis.

### RNA-seq Studies

The University of Texas at Austin
Waggoner Center for
Alcohol & Addiction Research

### Gómez-Carballa et al. (2023)

RNA-seq study, before and after 50 minutes of listening to classical music.

This study measured music's effect on the transcriptomes of cognitively healthy and those experiencing age-related cognitive decline (ACD). More prominent changes were observed in the ACD group than the healthy controls.

Gómez-Carballa, Alberto, Laura Navarro, Jacobo Pardo-Seco, Xabier Bello, Sara Pischedda, Sandra Viz-Lasheras, Alba Camino-Mera, et al. "Music Compensates for Altered Gene Expression in Age-Related Cognitive Disorders." *Scientific Reports* 13, no. 1 (December 2, 2023): 21259. https://doi.org/10.1038/s41598-023-48094-5.

### Gomez et al. (2023) - summary

- No samples were removed

The University of Texas at Austin
Waggoner Center for
Alcohol & Addiction Research

### Gómez-Carballa: filterByExpr and library sizes

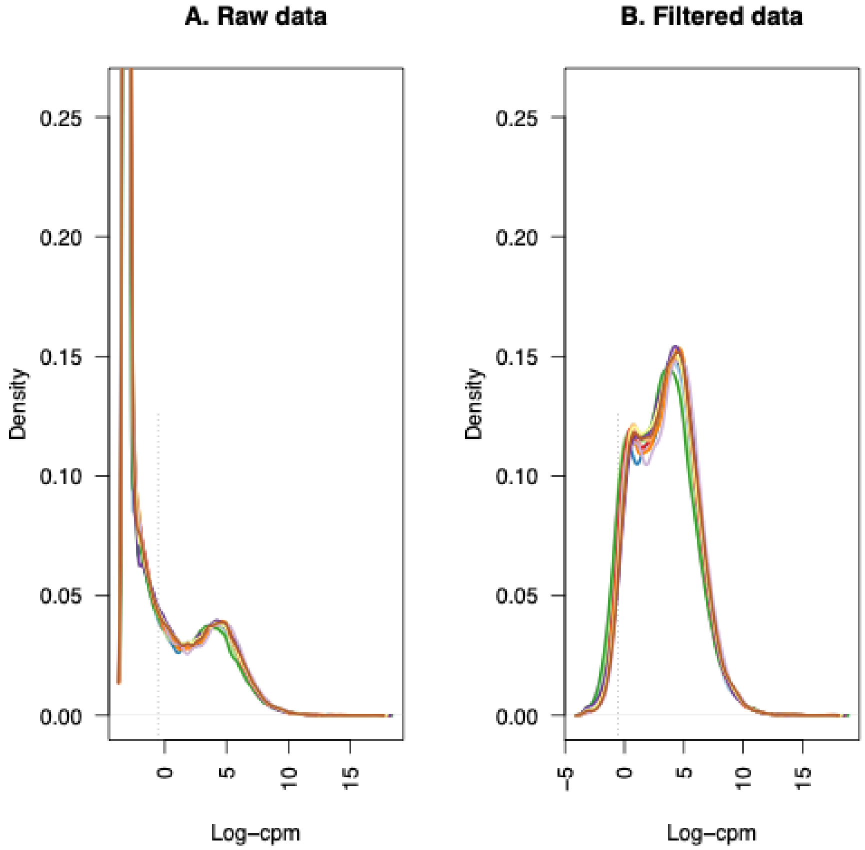


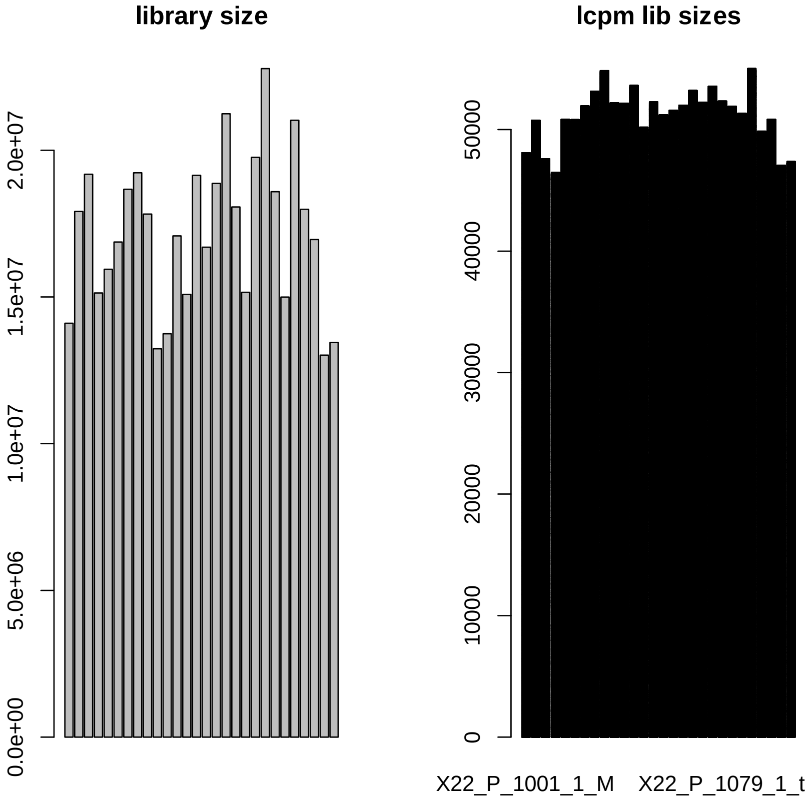

### Gómez-Carballa: PCA

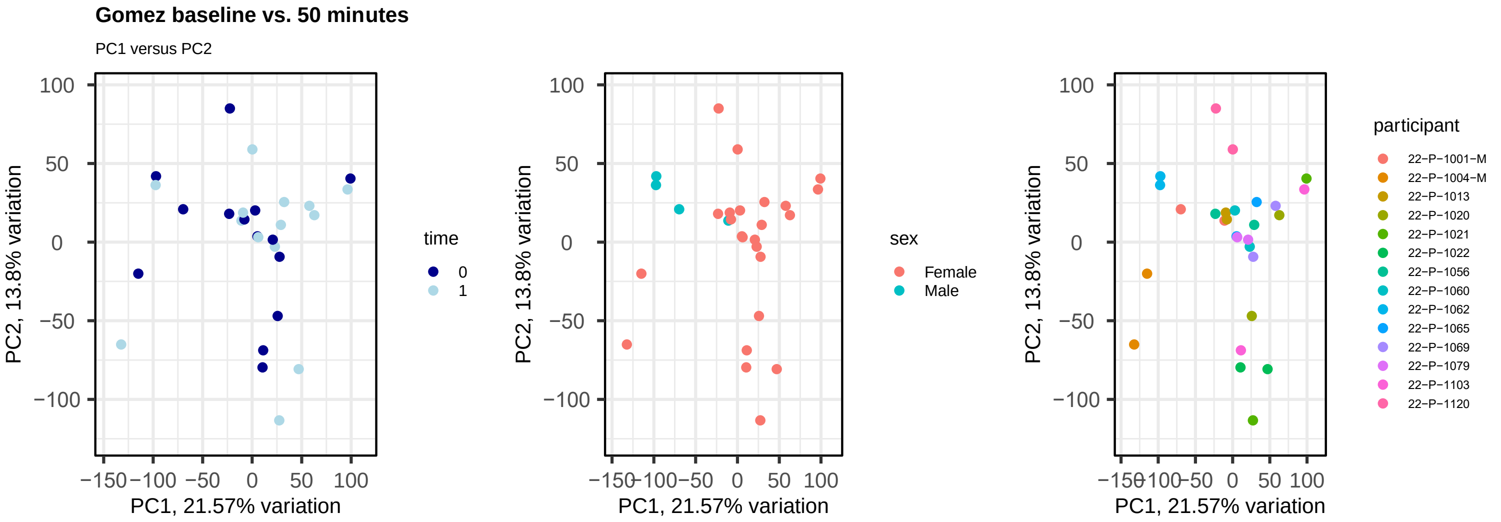


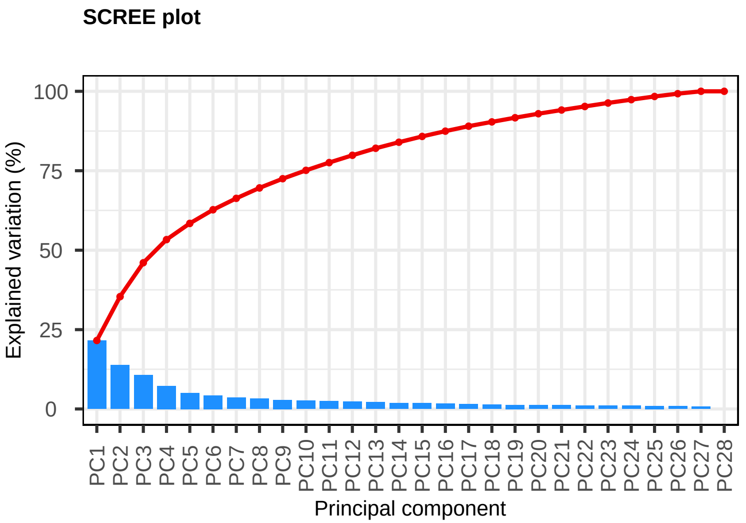


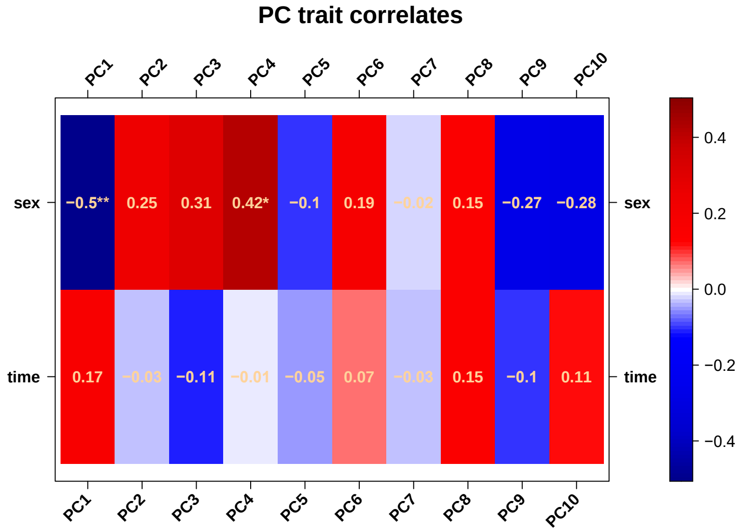

The University of Texas at Austin
Waggoner Center for
Alcohol & Addiction Research

### Gómez-Carballa: voom normalization

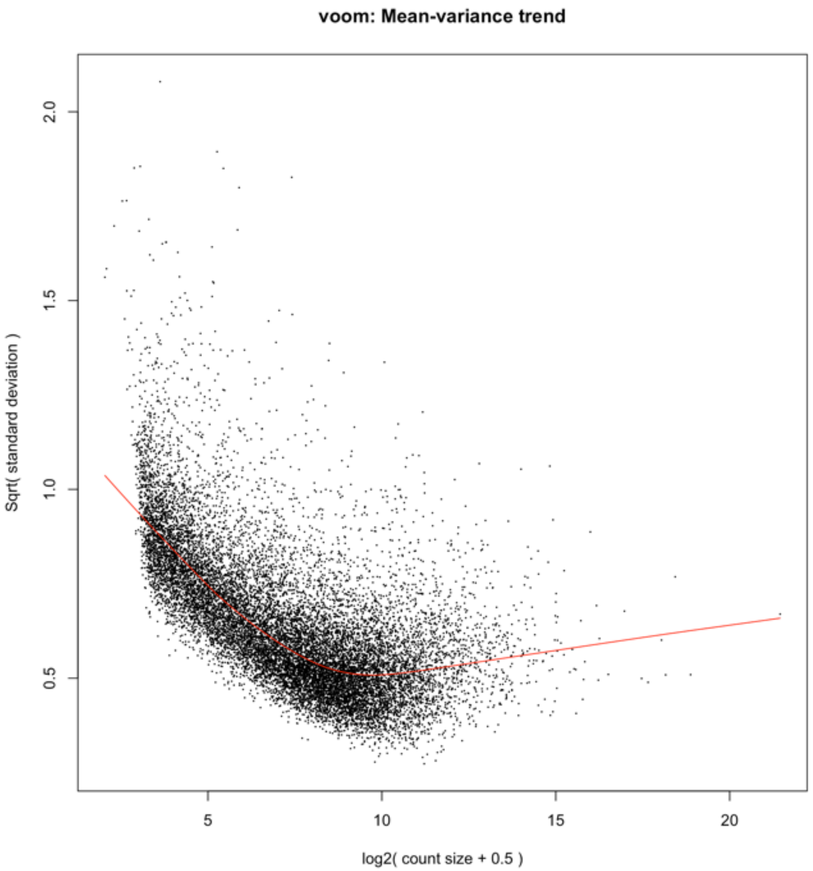


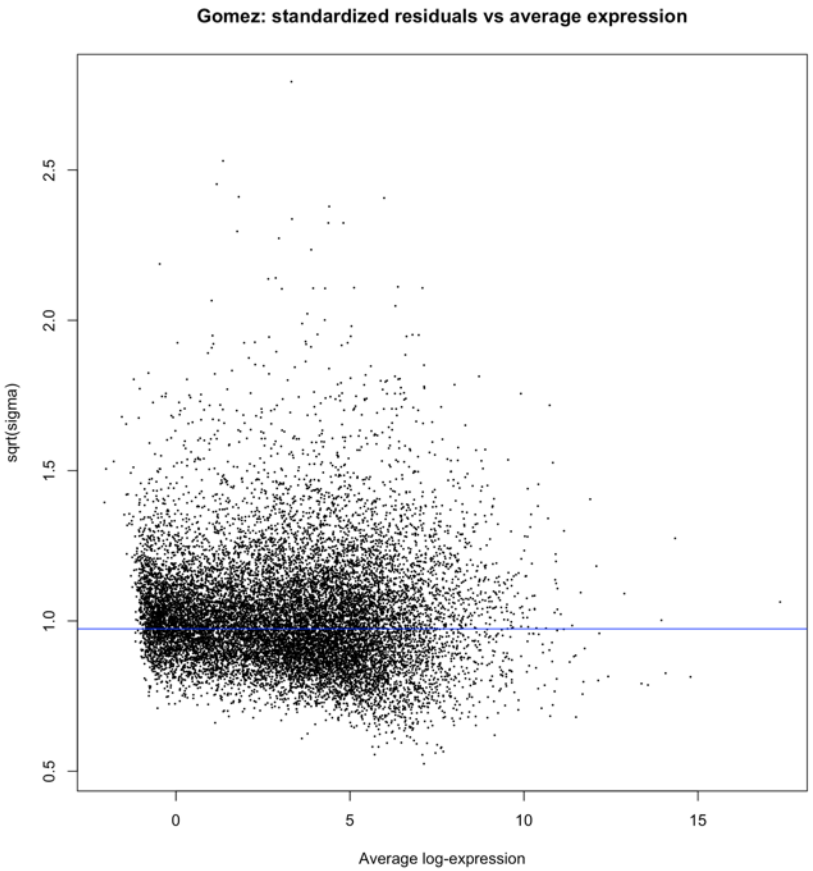

The University of Texas at Austin
Waggoner Center for
Alcohol & Addiction Research

### Gosch et al. (2023)

RNA-seq study, 3-hour time intervals for one day whole day without sleep.

The investigators in this study were interested in discovering blood RNA biomarkers for time-of-incident analysis in forensics. They measured non-fasting capillary blood.

Gosch, A., A. Bhardwaj, and C. Courts. "TrACES of Time: Transcriptomic Analyses for the Contextualization of Evidential Stains – Identification of RNA Markers for Estimating Time-of-Day of Bloodstain Deposition." *Forensic Science International: Genetics* 67 (November 1, 2023): 102915. https://doi.org/10.1016/j.fsigen.2023.102915.

### Gosch et al. (2023) - summary

| Sample | Subject ID | Sex | Time Point | Reason Removed |
|---|---|---|---|---|
| **GSM6965634** | J | Female | 23:00 | Library size outlier |

### Gosch: filterByExpr and library sizes

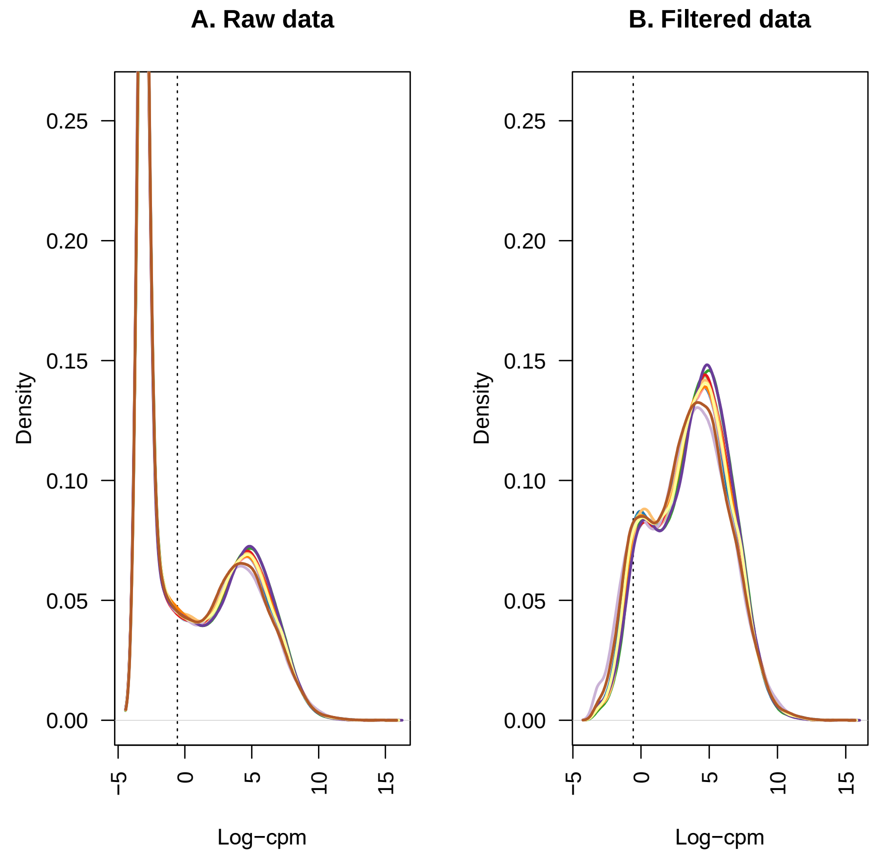


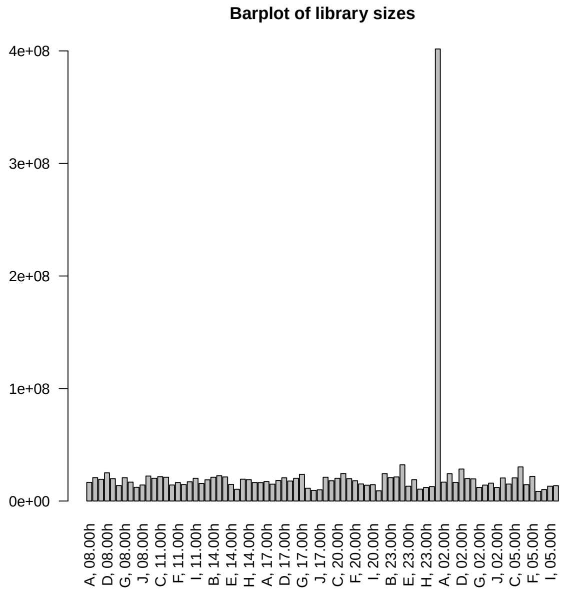


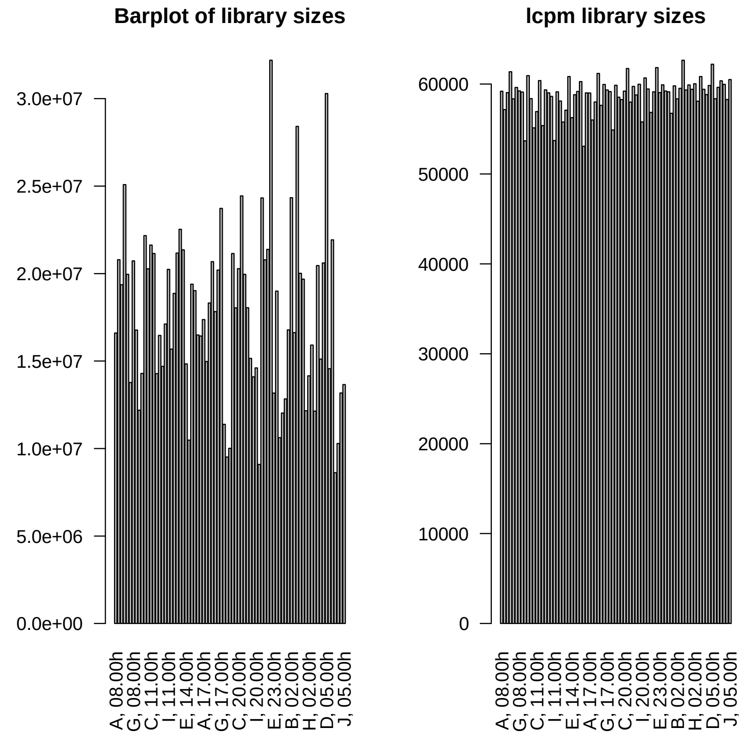


The original data had a sample with unusually large library size (right– sample **GSM6965634**). We removed GSM6965634 from the data prior to filtration and normalization (left two plots).

The University of Texas at Austin
Waggoner Center for
Alcohol & Addiction Research

### Gosch: PCA

**Gomez baseline vs. 50 minutes**

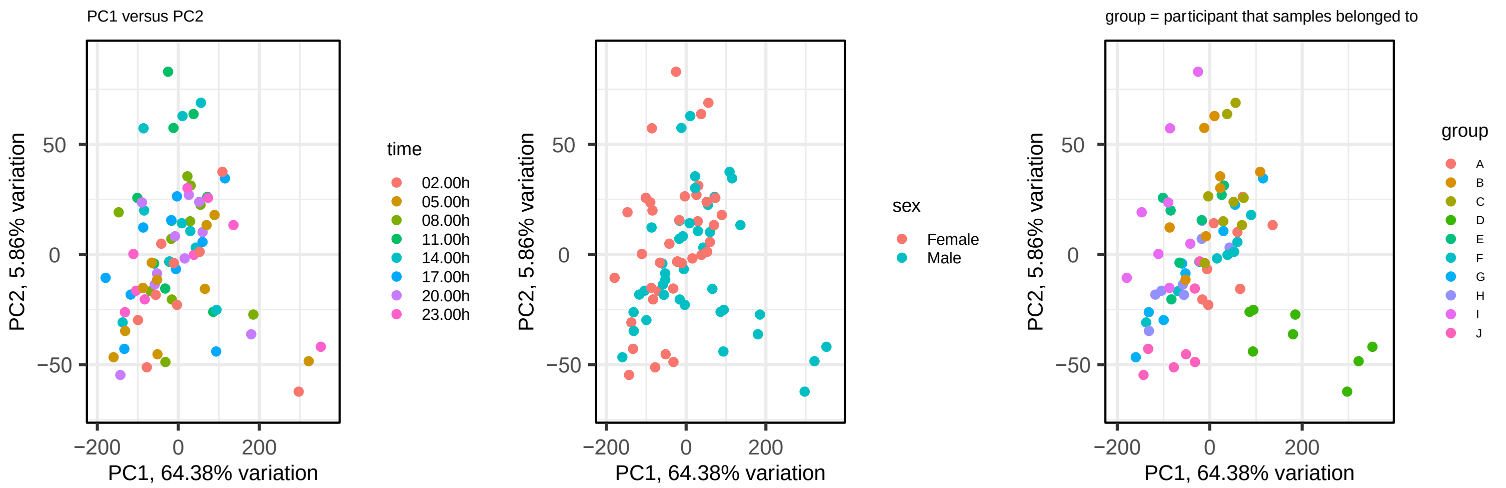


**SCREE plot**

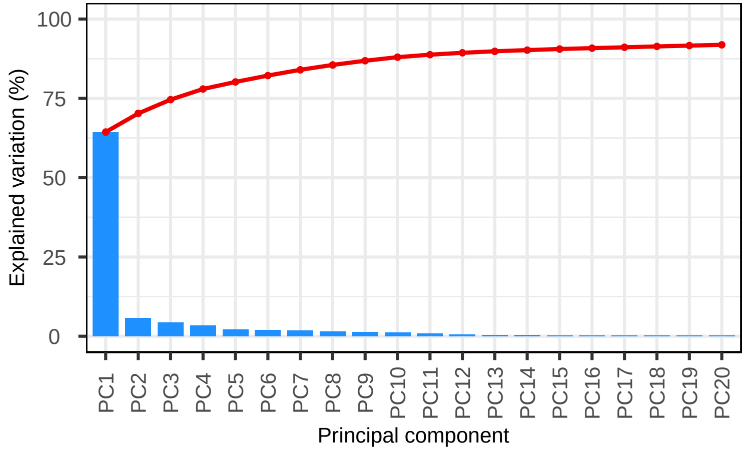


**PC trait correlates**

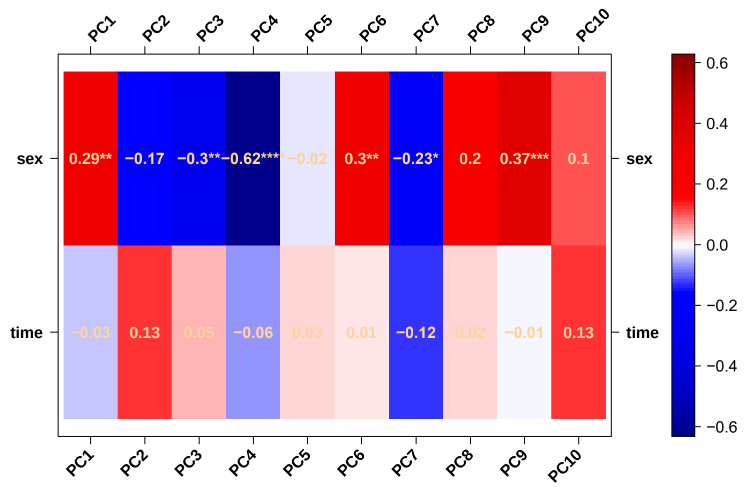

The University of Texas at Austin
Waggoner Center for
Alcohol & Addiction Research

### Gosch: voom normalization

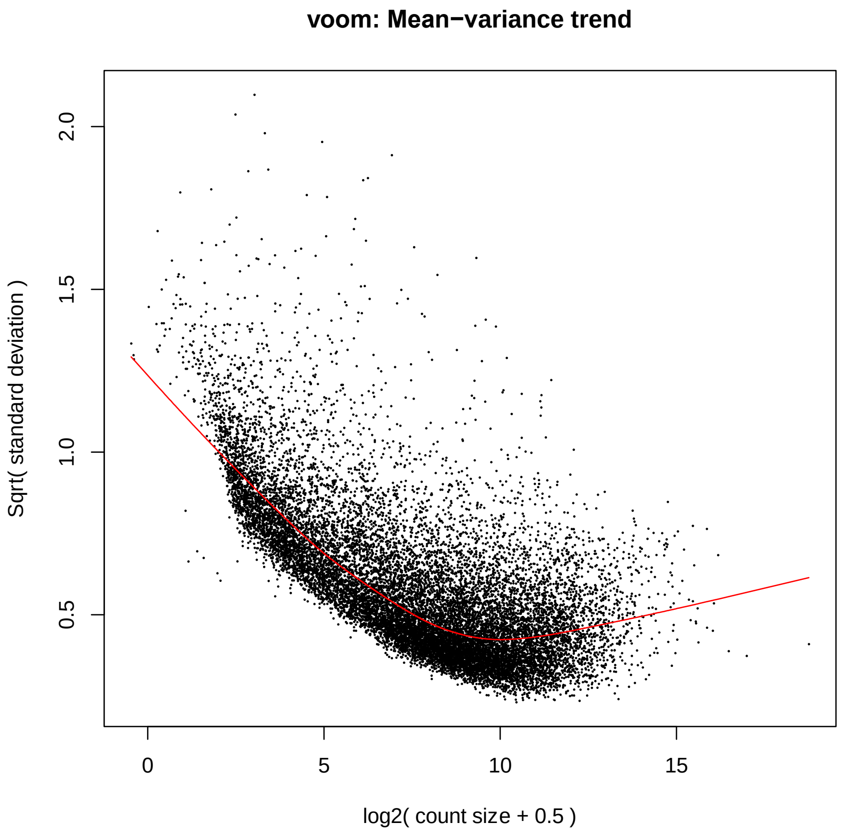


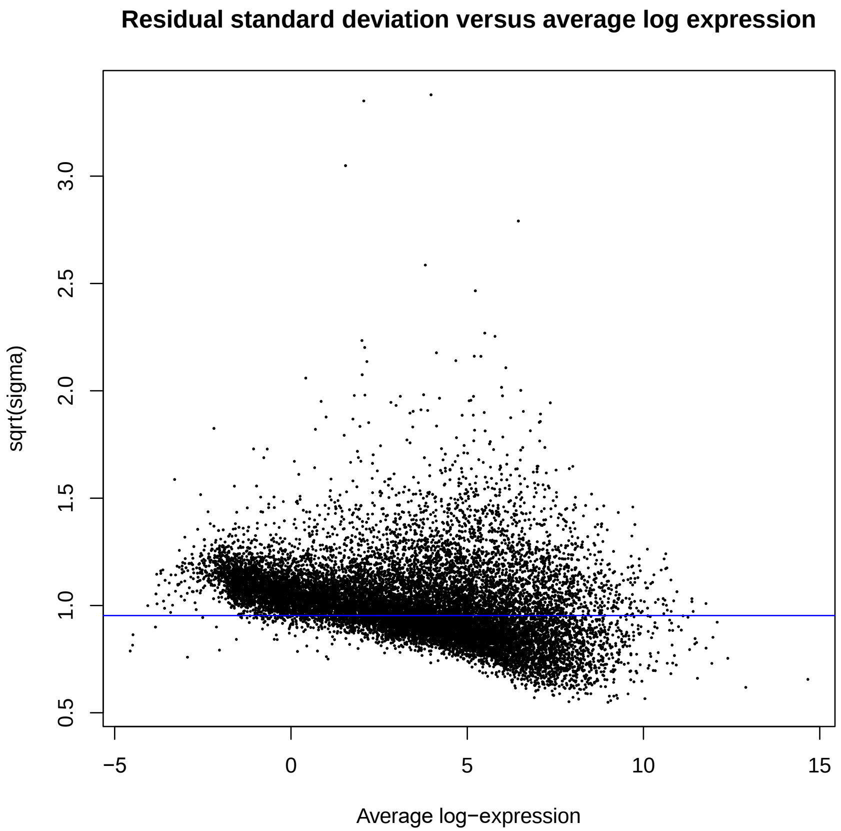

### Illumina HumanHT-12 v3.0 Studies

### Obermoser et al. (2013)

Microarray study using Illumina HT12 v3 BeadChip. During the study, participants had blood drawn before and after receiving an injection of either the 2009–2010 seasonal influenza vaccine (Fluzone®), 23-valent pneumococcal vaccine (Pneumovax23®), or saline injections. All participants were healthy at the time of vaccination.

Blood samples were collected by venipuncture at days −7, 0, 1,3, 7, 10, 14, 21, and 28.

Obermoser, Gerlinde, Scott Presnell, Kelly Domico, Hui Xu, Yuanyuan Wang, Esperanza Anguiano, LuAnn Thompson-Snipes, et al. "Systems Scale Interactive Exploration Reveals Quantitative and Qualitative Differences in Response to Influenza and Pneumococcal Vaccines." *Immunity* 38, no. 4 (April 18, 2013): 831–44. https://doi.org/10.1016/j.immuni.2012.12.008.

### Obermoser et al. (2013) - summary

| Sample | GEO accession | Sex | Time Point | Reason Removed |
|---|---|---|---|---|
| All samples treated “Saline” in the Training_Set_Vein cohort | GSM744889, GSM744890, GSM744891, GSM744892, GSM744893, GSM744894, GSM744895, GSM744896, GSM744897, GSM744898, GSM744899, GSM744900, GSM744901, GSM744902, GSM744903, GSM744904, GSM744905, GSM744906, GSM744907, GSM744908, GSM744909, GSM744910, GSM744911, GSM744912, GSM744913, GSM744914, GSM744915, GSM744916, GSM744917, GSM744918, GSM744919, GSM744920, GSM744921, GSM744922, GSM744923, GSM744924, GSM744925, GSM744926, GSM744927, GSM744928, GSM744929, GSM744930, GSM744931, GSM744932, GSM744933, GSM744934, GSM744935, GSM744936, GSM744937, GSM744938, GSM744939, GSM744940, GSM744941 | 27F, 26M samples | Baseline, treatment (days after baseline), and 1, 3, 7, 10, 14, 21, and 28 days after treatment | Poor inter-individual correlation and anomalous clustering on PCA |
| Validation_Vein cohort | GSM745184, GSM745185, GSM745186, GSM745187, GSM745188, GSM745189, GSM745190, GSM745191, GSM745192, GSM745193, GSM745194, GSM745195, GSM745196, GSM745197, GSM745198, GSM745199, GSM745200, GSM745201, GSM745202, GSM745203, GSM745204, GSM745205, GSM745206, GSM745207, GSM745208, GSM745209, GSM745210, GSM745211, GSM745212, GSM745213 | 15F, 15M samples | Baseline, 1 day, 28 days | No control (saline) samples in this cohort |
| All samples not labeled “Whole Blood” for sample | GSM744141, GSM744142, GSM744143, GSM744144, GSM744145, GSM744146, GSM744147, GSM744148, GSM744149, GSM744150, GSM744151, GSM744152, GSM744153, GSM744154, GSM744155, GSM744156, GSM744157, GSM744158, GSM744159, GSM744160, GSM744161, GSM744162, GSM744163, GSM744164, GSM744165, GSM744166, GSM744167, GSM744168, GSM744169, GSM744170, GSM744171, GSM744172, GSM744173, GSM744174, GSM744175, GSM744176, GSM744177, GSM744178, GSM744179, GSM744180, GSM744181, GSM744182, GSM744183, GSM744184, GSM744185, GSM744186, GSM744187, GSM744188, GSM744189, GSM744190, GSM744191, GSM744192, GSM744193, GSM744194, GSM744195, GSM744196, GSM744197, GSM744198, GSM744199, GSM744200, GSM744201, GSM744202, GSM744203, GSM744204, GSM744205, GSM744206, GSM744207, GSM744208, GSM744209, GSM744210, GSM744211, GSM744212 | 36F, 36M samples | Baseline or follow-up | These samples only measured PBMCs |

The University of Texas at Austin
Waggoner Center for
Alcohol & Addiction Research

### Obermoser: cohort deconvolution

Unsupervised clustering of filtered, normalized data from all cohorts.

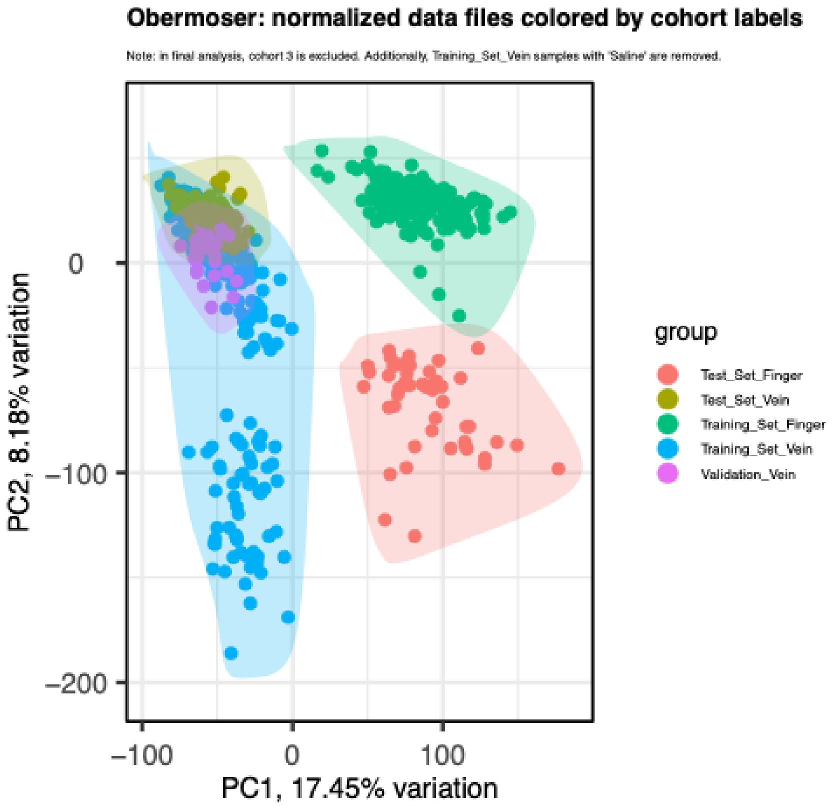


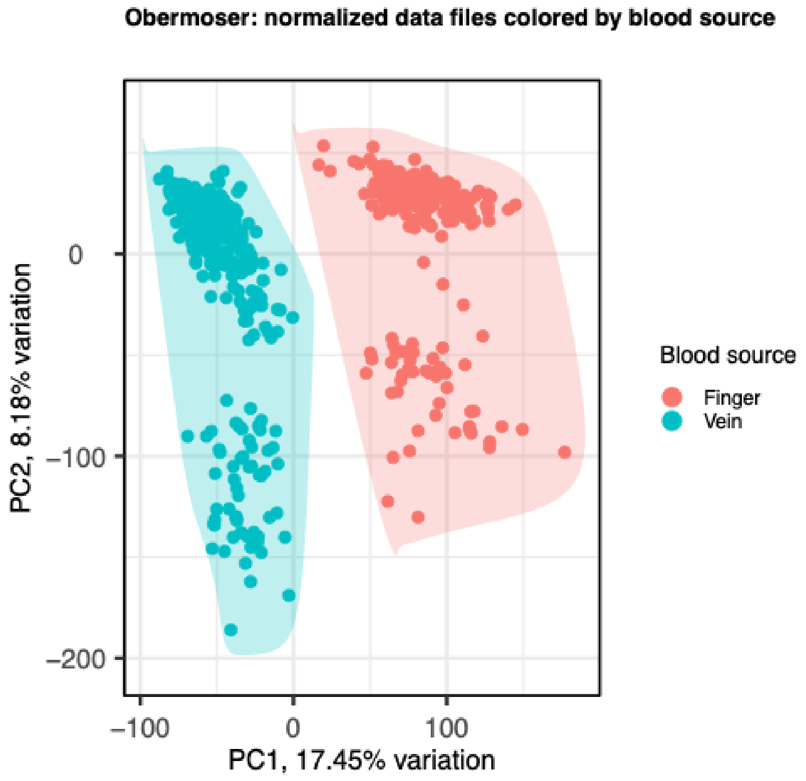


The samples cluster according to the blood used on the x-axis (capillary vs vein, rightmost plot) and sampling cohort (leftmost plot). **Training_Set_Vein** belongs to cohort 1; **Test_Set_Vein, Test_Set_Finger, and Training_Set_Finger** belong to cohort 2. **Validation_Vein** belongs to cohort 3; however, we did not include these samples in our analysis because no control group was included (i.e., nobody received saline).

The University of Texas at Austin
Waggoner Center for
Alcohol & Addiction Research

### Obermoser: cohort deconvolution

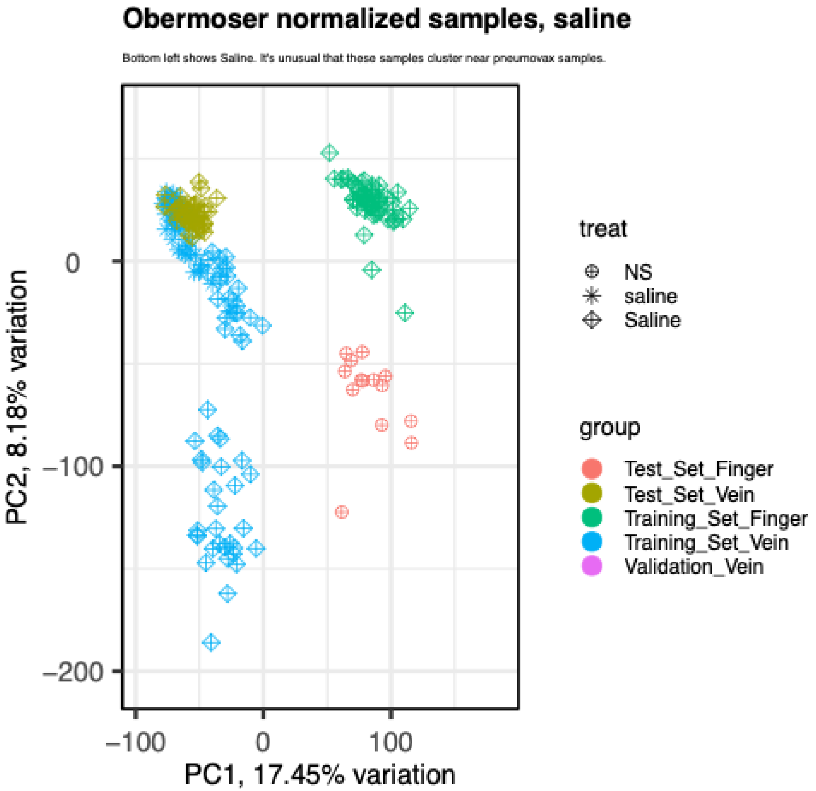


(NS - normal saline)

Saline - Saline

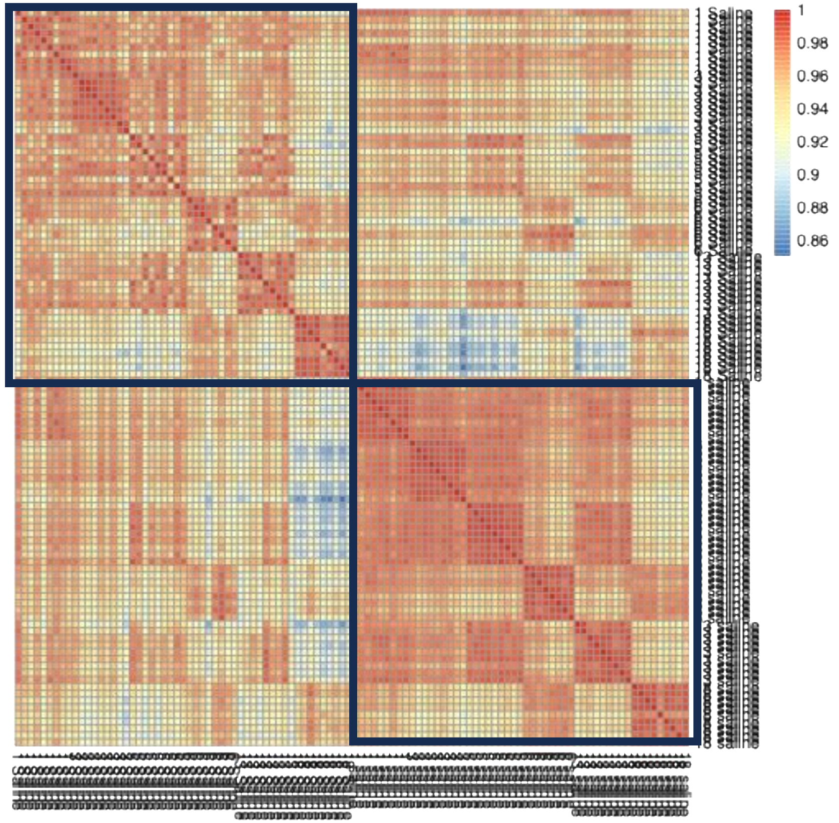


saline - saline

**Training_Set_Vein** had two sets of identically labeled samples, the only difference being the treatment group: 'saline' and 'Saline'. We could not find information about this in the manuscript or on GEO, so we used clustering to guide our decision on which set to use for downstream analyses. We used the 'saline' samples because they show more agreement between samples (measured by correlation coefficient, rightmost plot) and they cluster near other saline samples from other vein cohorts (left).

### Obermoser: PCA (Training_Vein)*

*Does not contain 'Saline' samples; only 'saline' samples. Normalized and filtered.

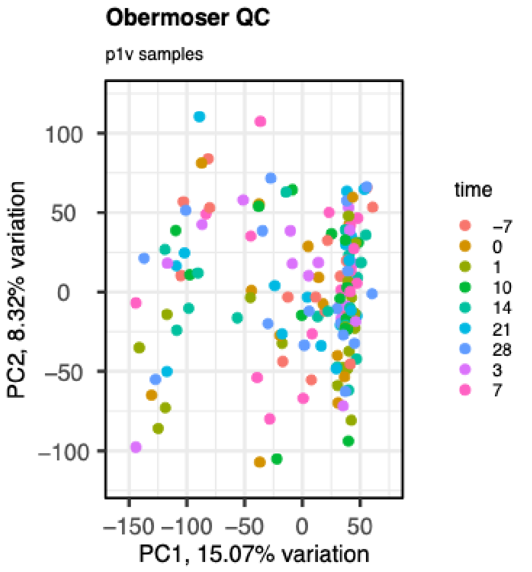


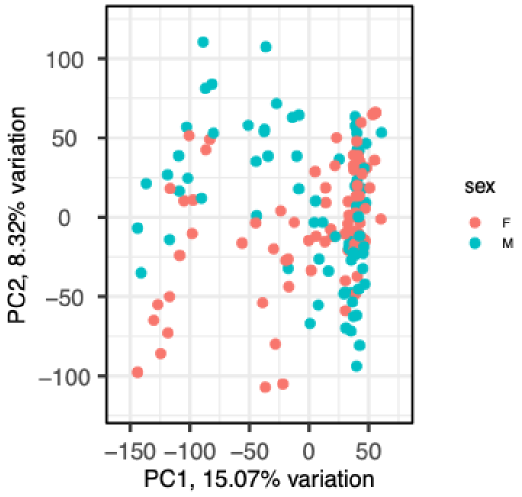


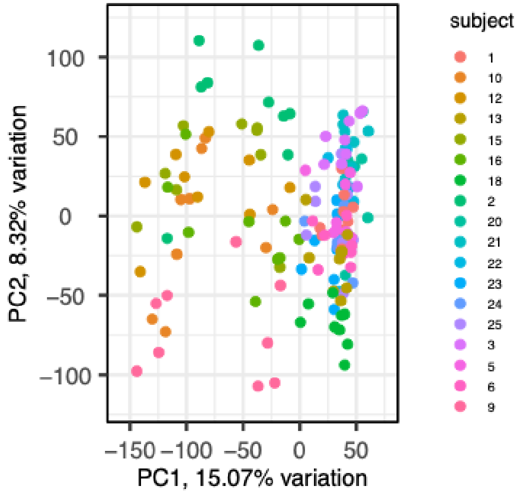


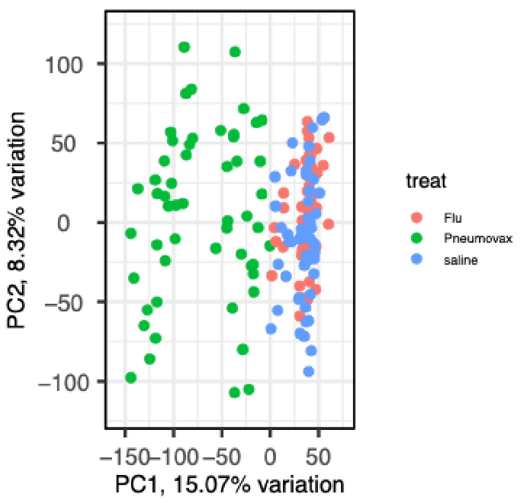


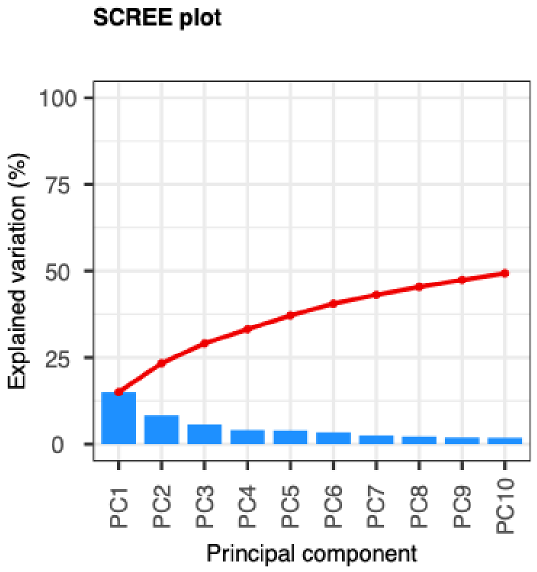


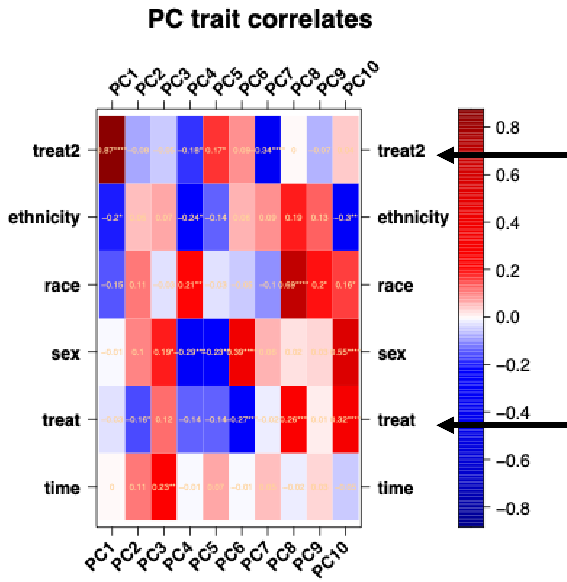


2-factor variable (saline and fluzone collapsed)

3-factor treatment variable (saline, fluzone, and pneumovax)

### Obermoser: PCA (Training_Vein)*

*Does not contain 'Saline' samples; only 'saline' samples. Normalized and filtered.

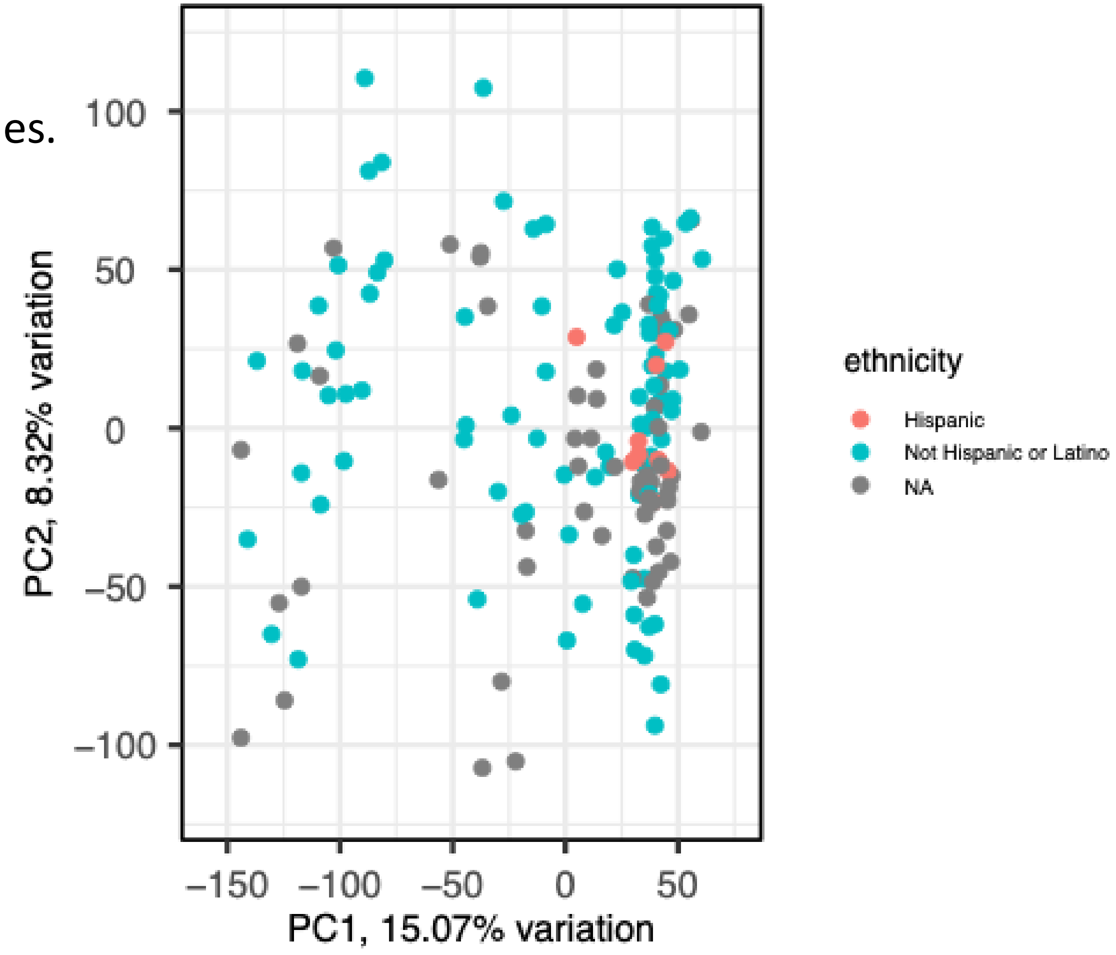


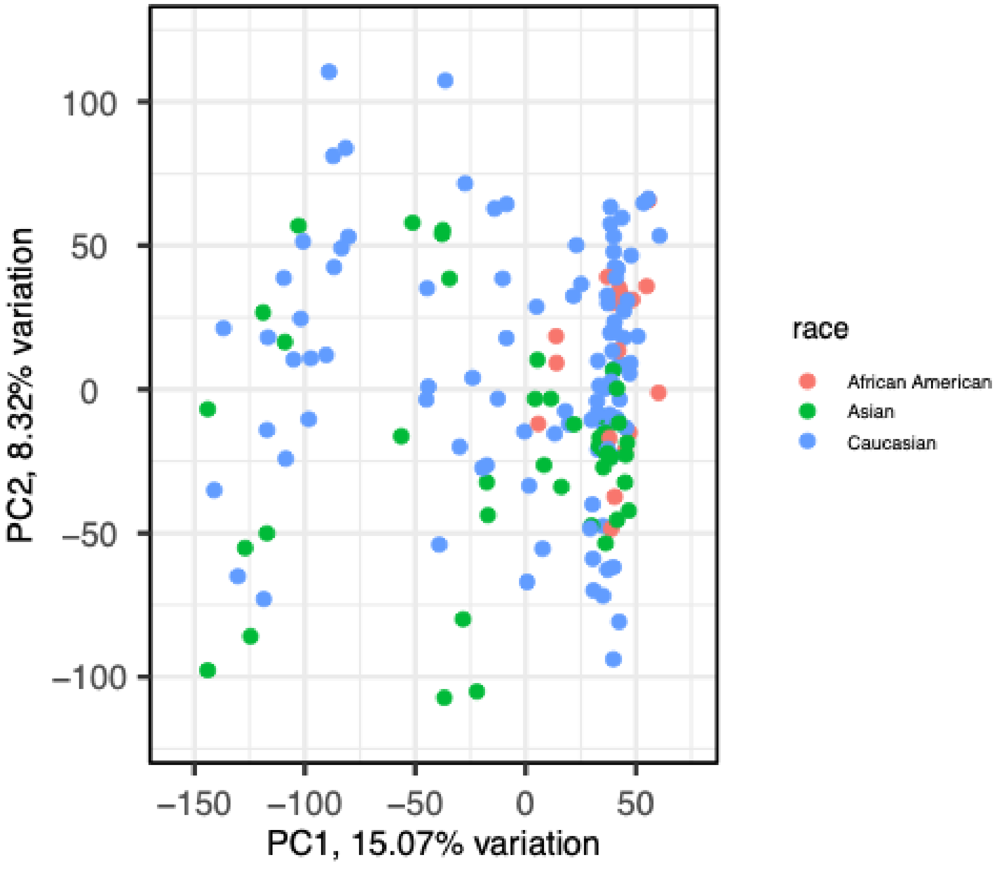

The University of Texas at Austin
Waggoner Center for
Alcohol & Addiction Research

### Obermoser: Sample boxplots (Training_Vein)*

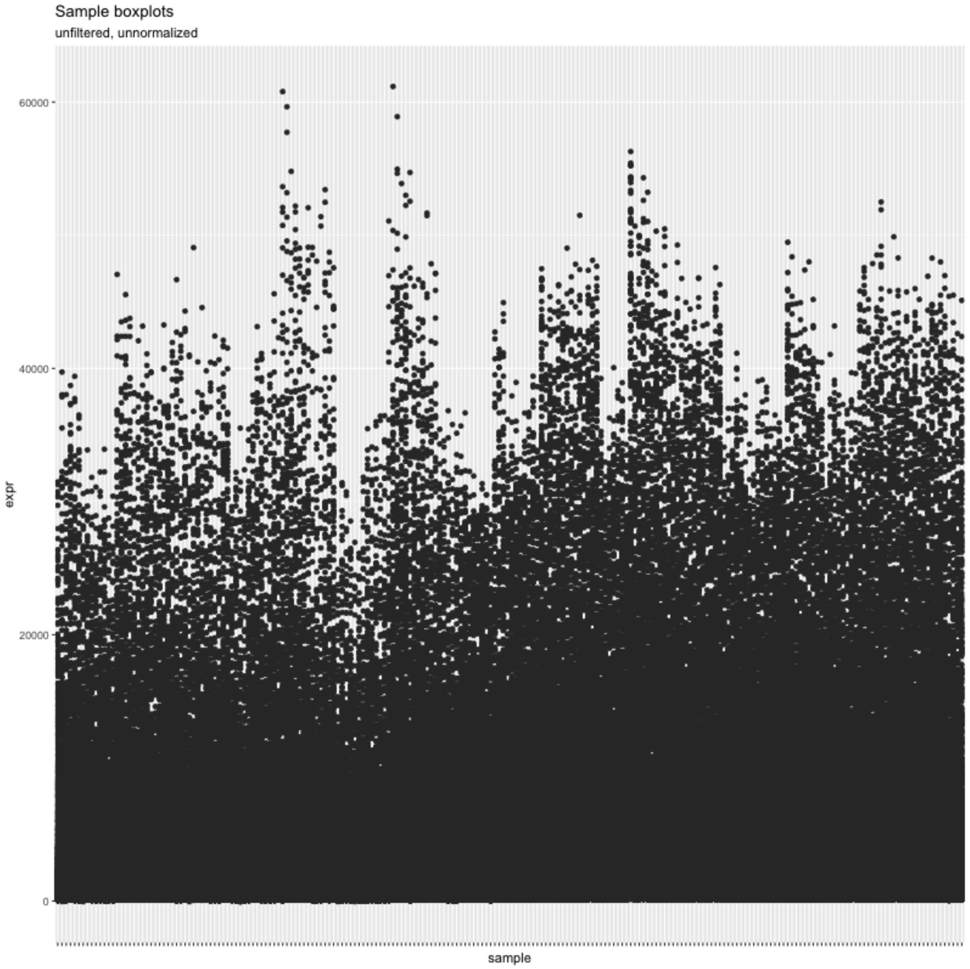


Unnormalized, unfiltered

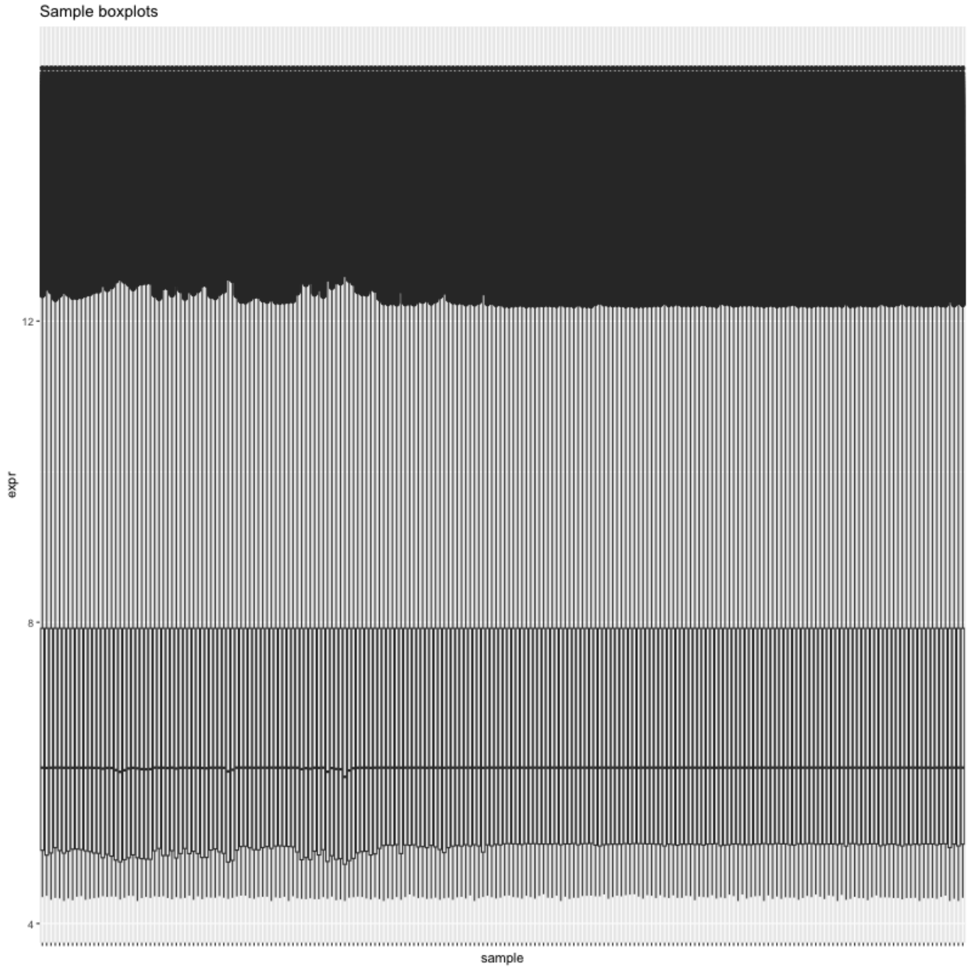


Normalized, filtered

The University of Texas at Austin
Waggoner Center for Alcohol & Addiction Research

### Obermoser: PCA (Test_Vein)

Normalized and filtered.

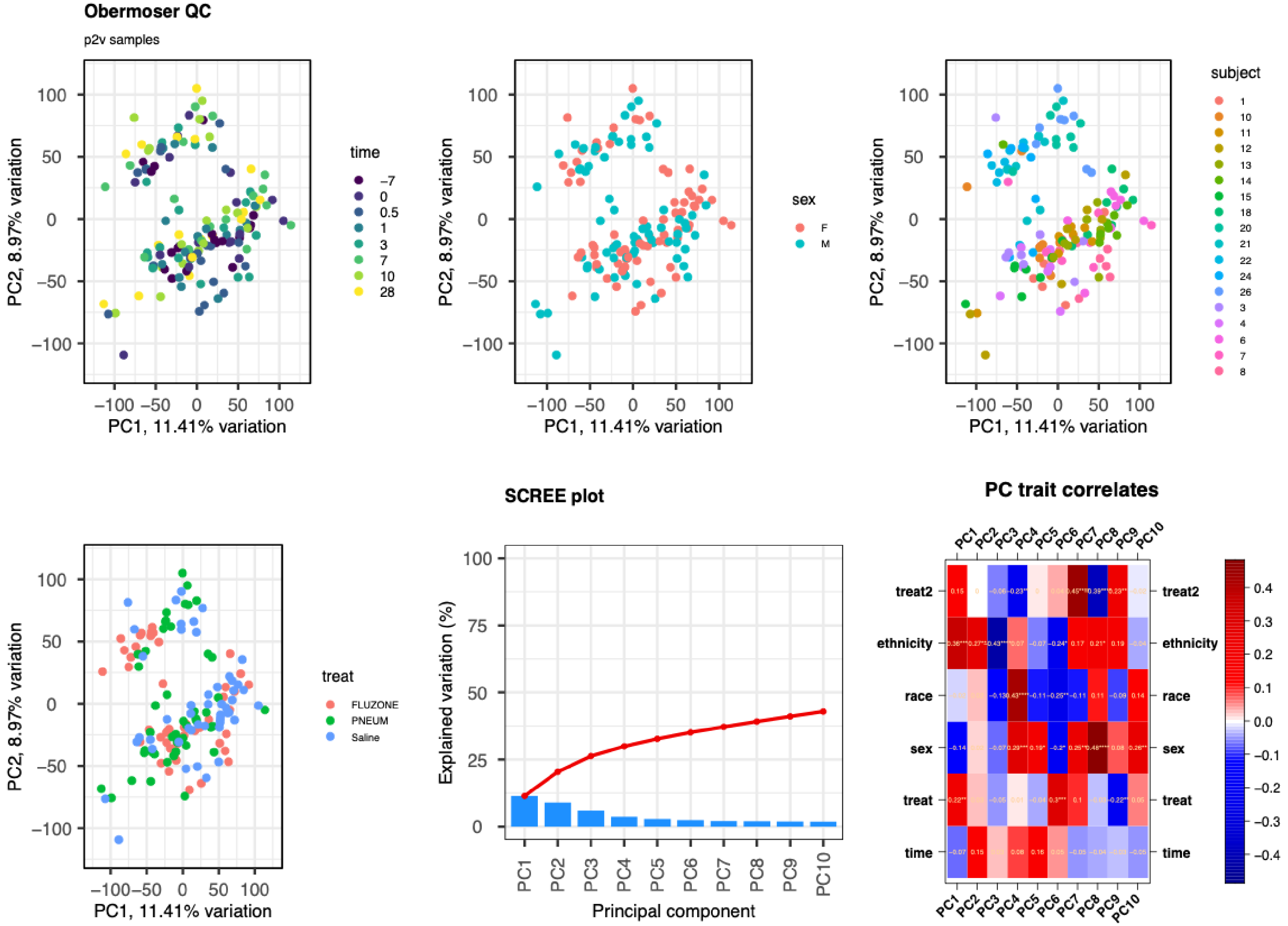

The University of Texas at Austin
Waggoner Center for
Alcohol & Addiction Research

### Obermoser: PCA (Test_Vein)

Normalized and filtered.

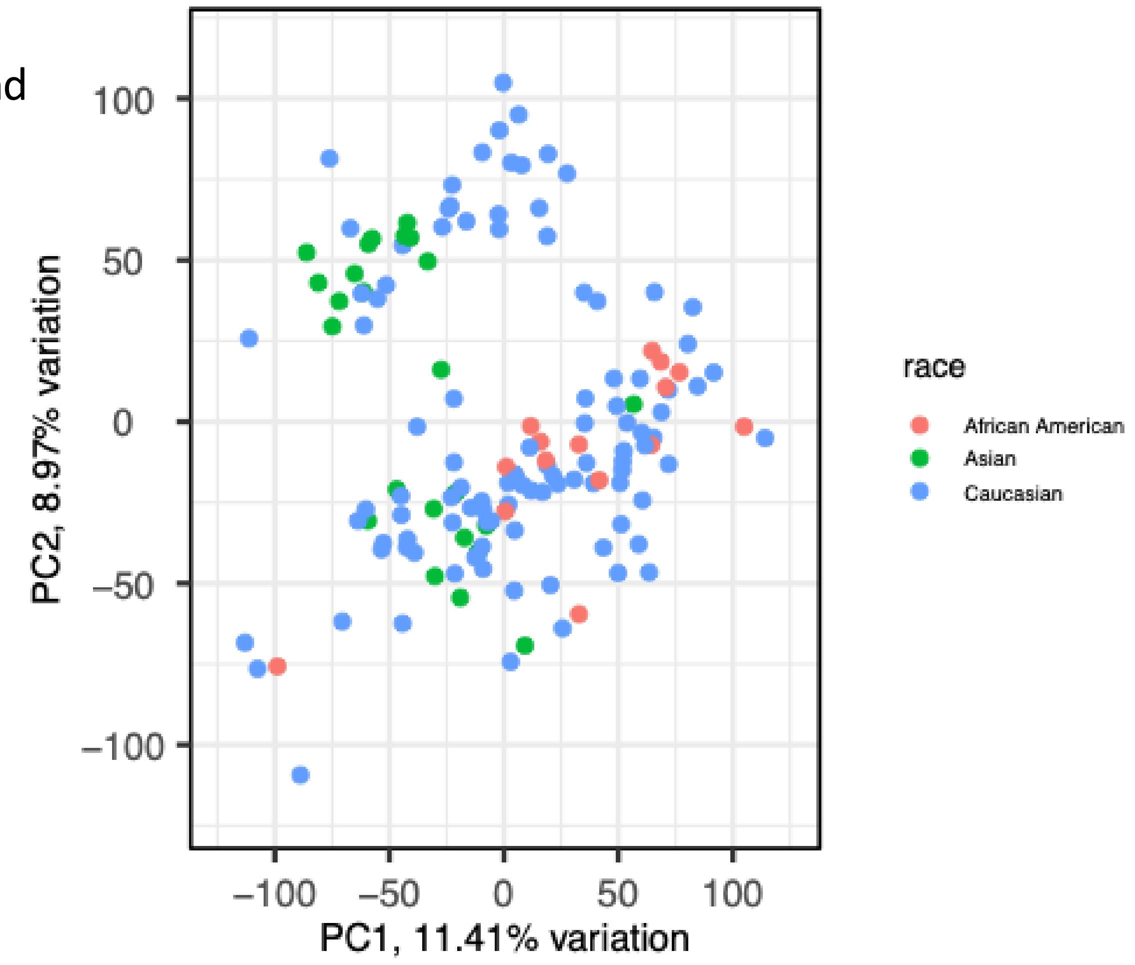


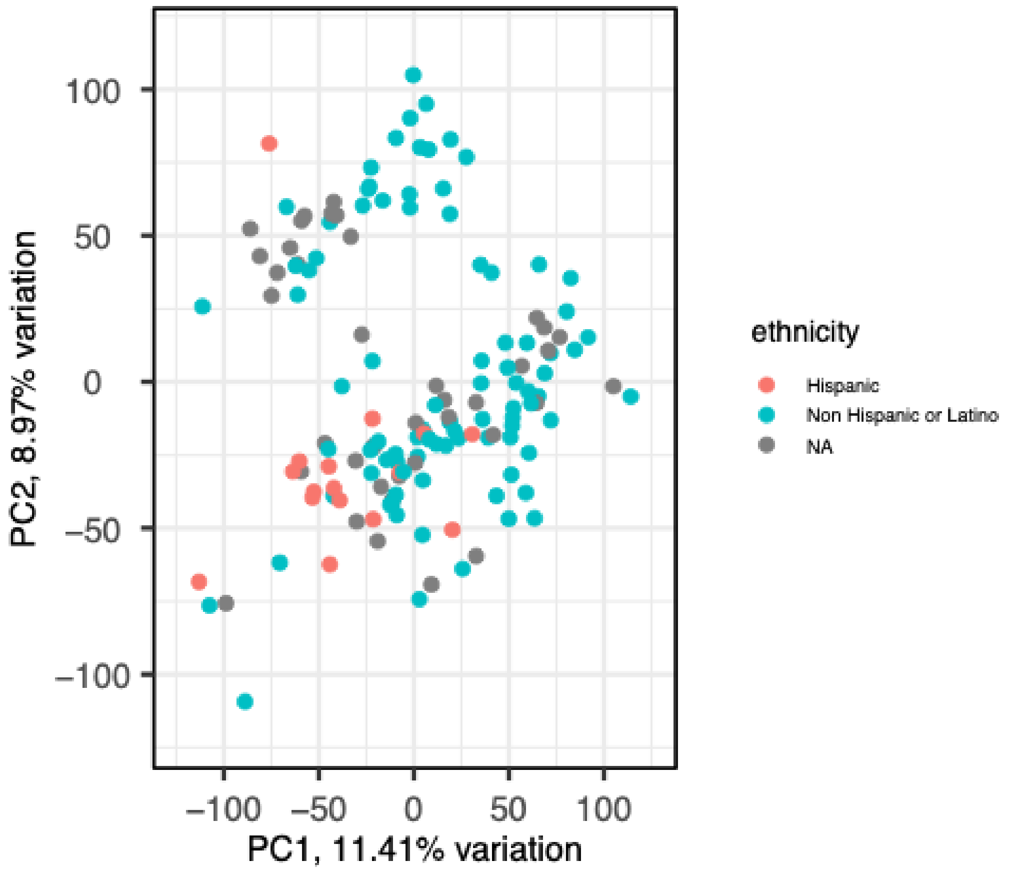

The University of Texas at Austin
Waggoner Center for
Alcohol & Addiction Research

### Obermoser: Sample boxplots (Test_Vein)

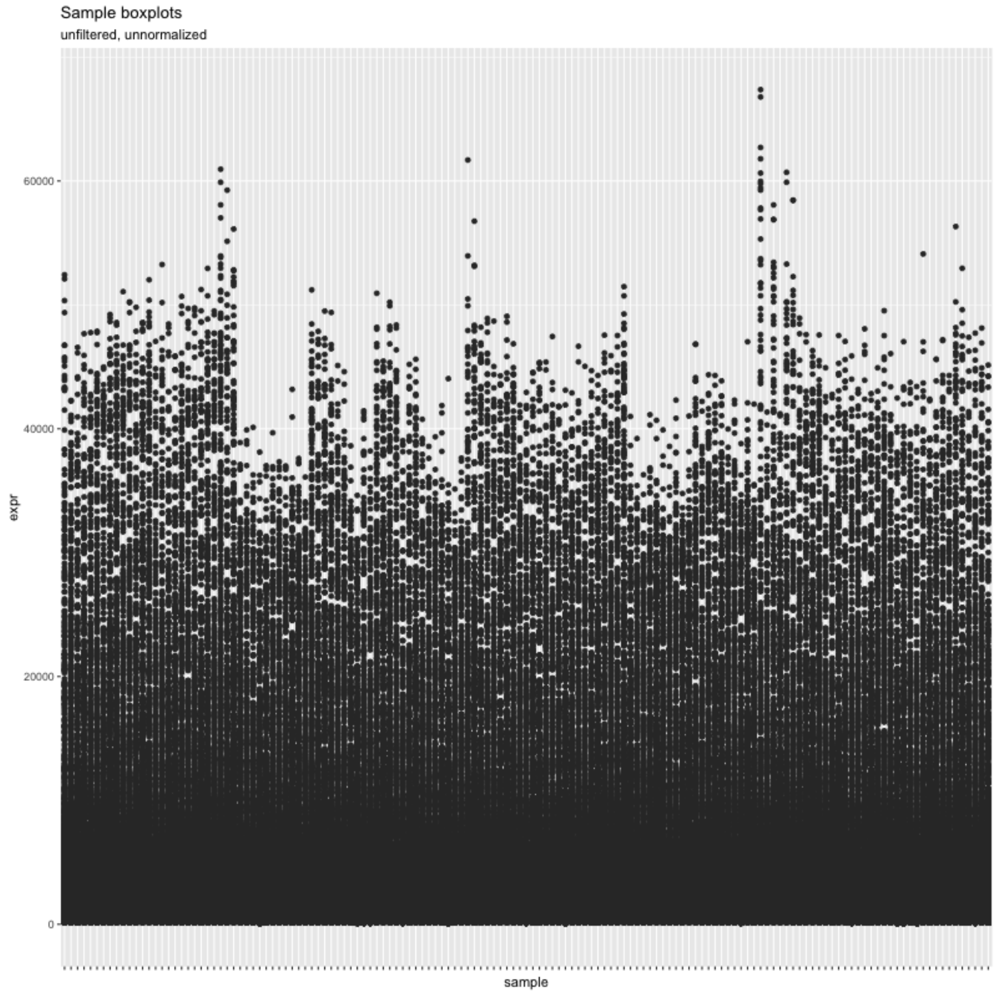


Unnormalized, unfiltered

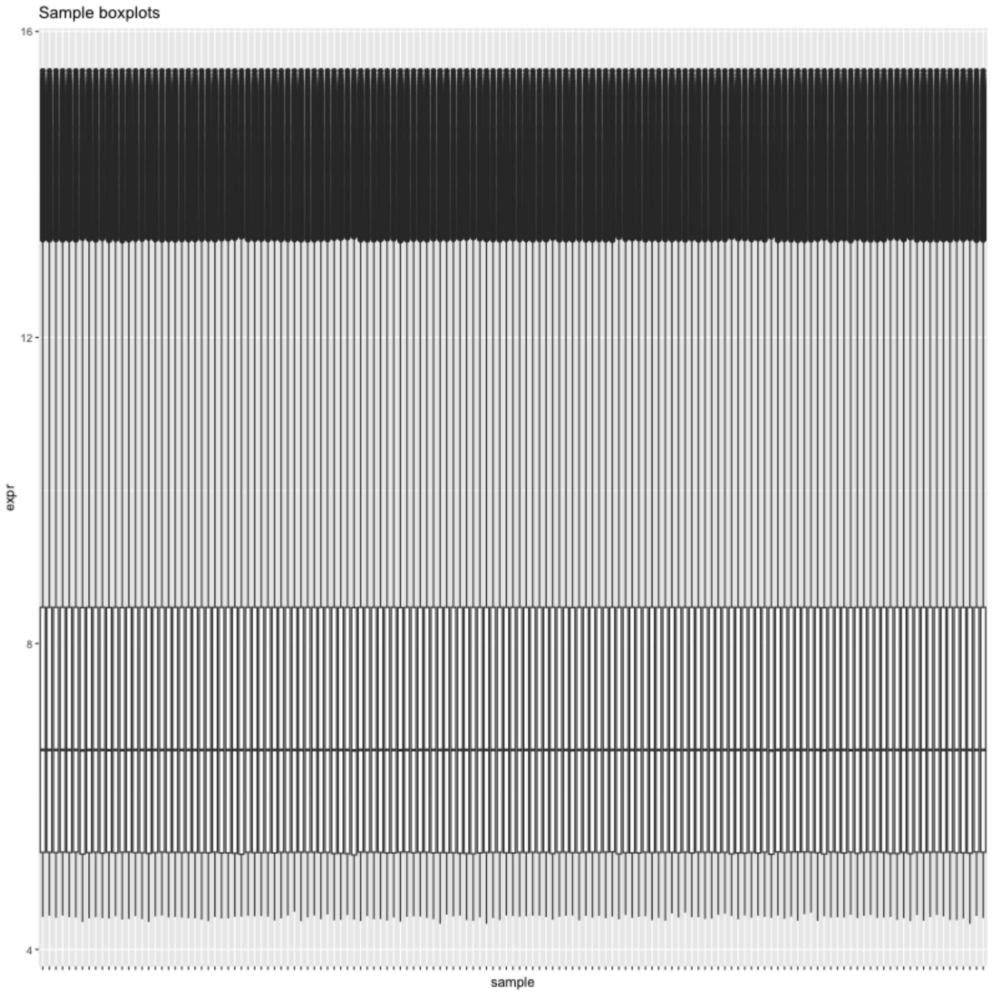


Normalized, filtered

The University of Texas at Austin
Waggoner Center for
Alcohol & Addiction Research

### Obermoser: PCA (Training_Finger)

Normalized and filtered.

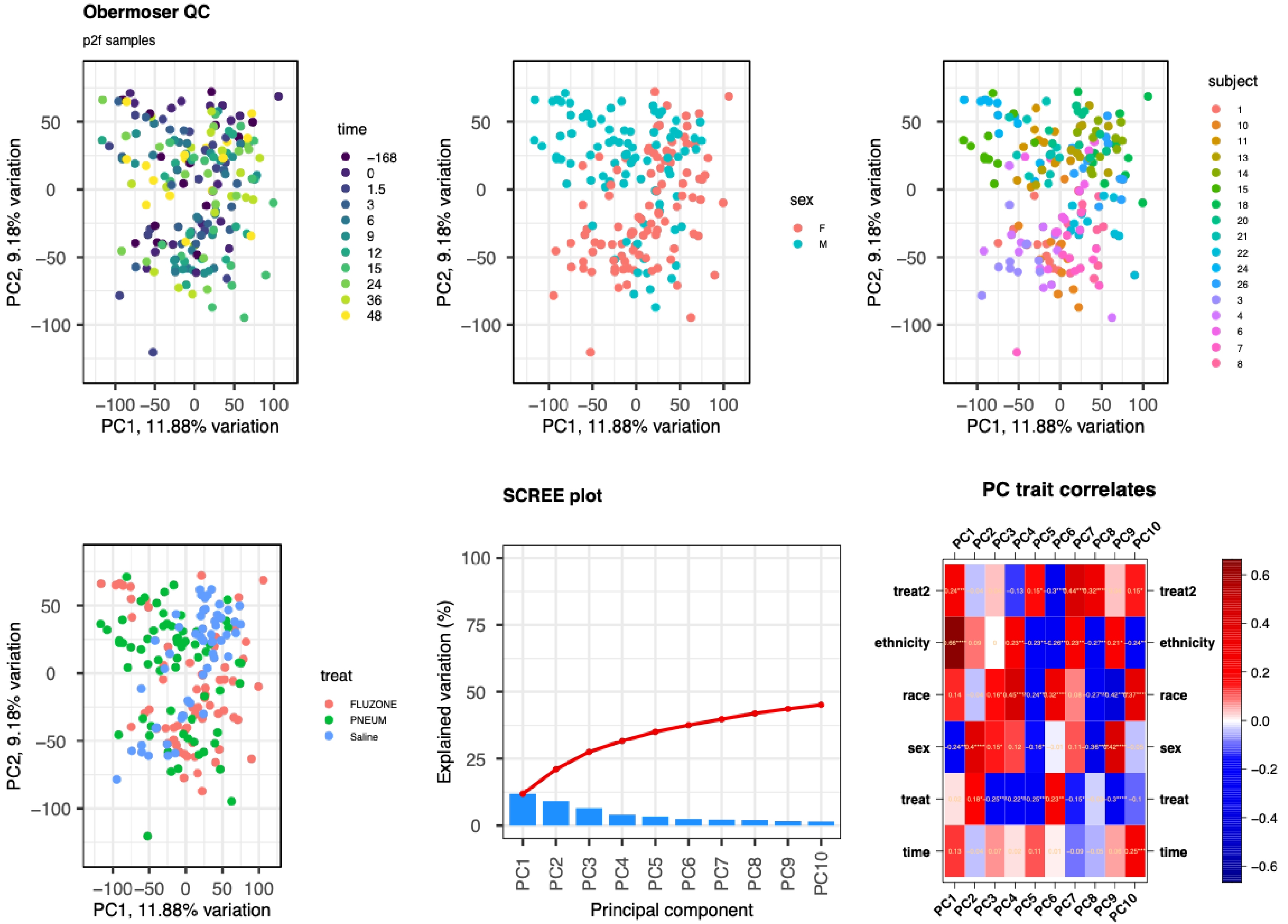

### Obermoser: PCA (Training_Finger)

Normalized and filtered.

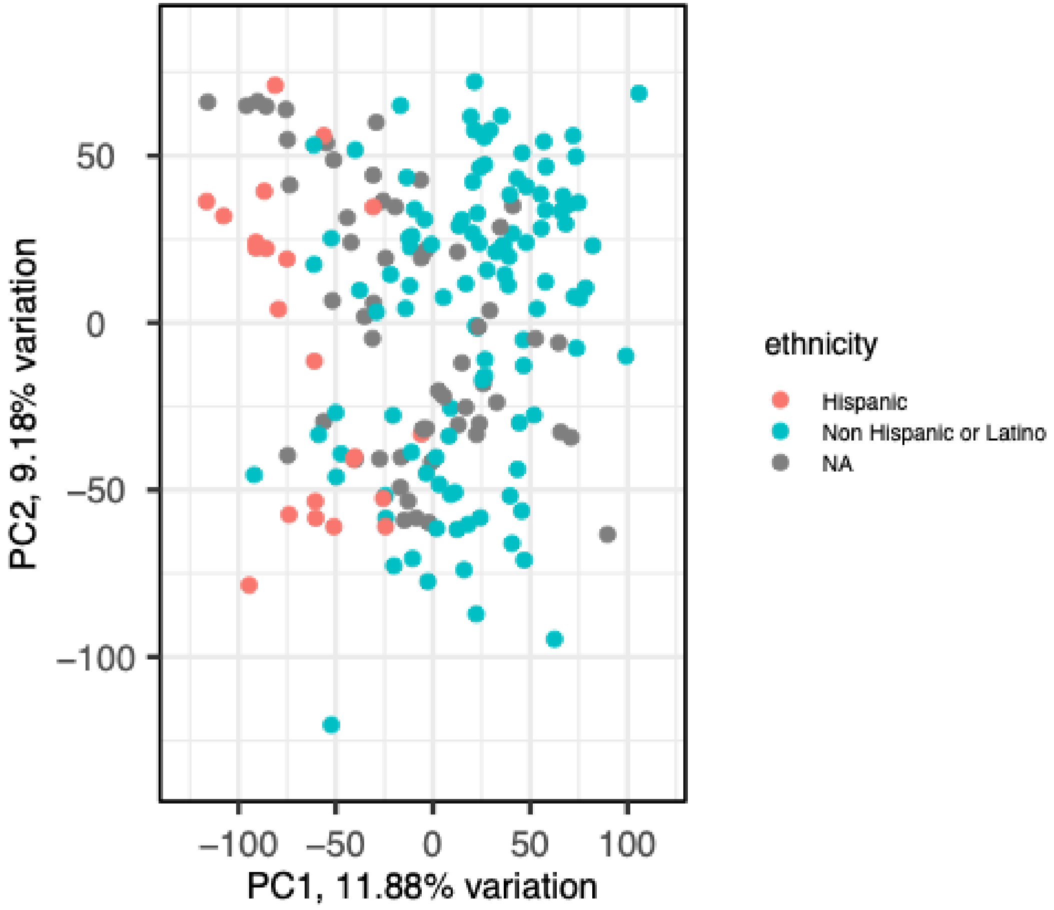


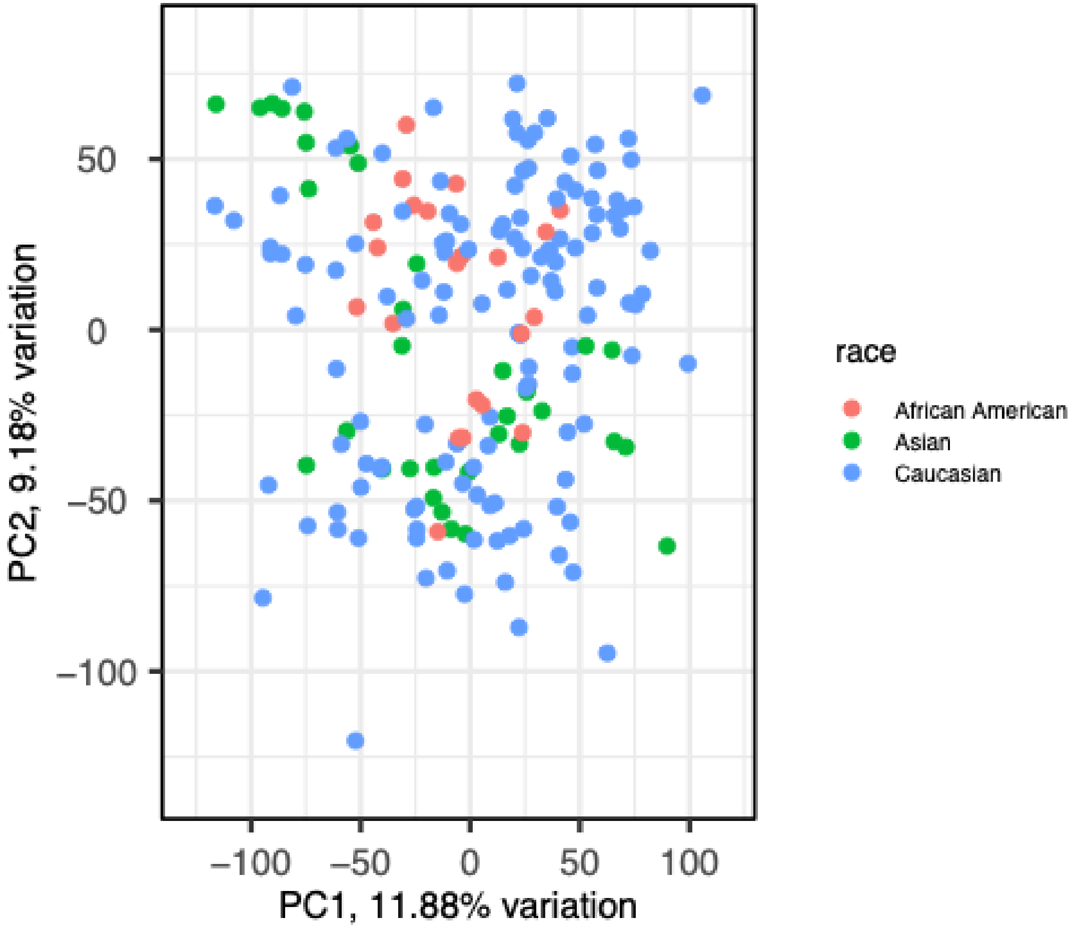

The University of Texas at Austin
Waggoner Center for
Alcohol & Addiction Research

### Obermoser: Sample boxplots (Training_finger)

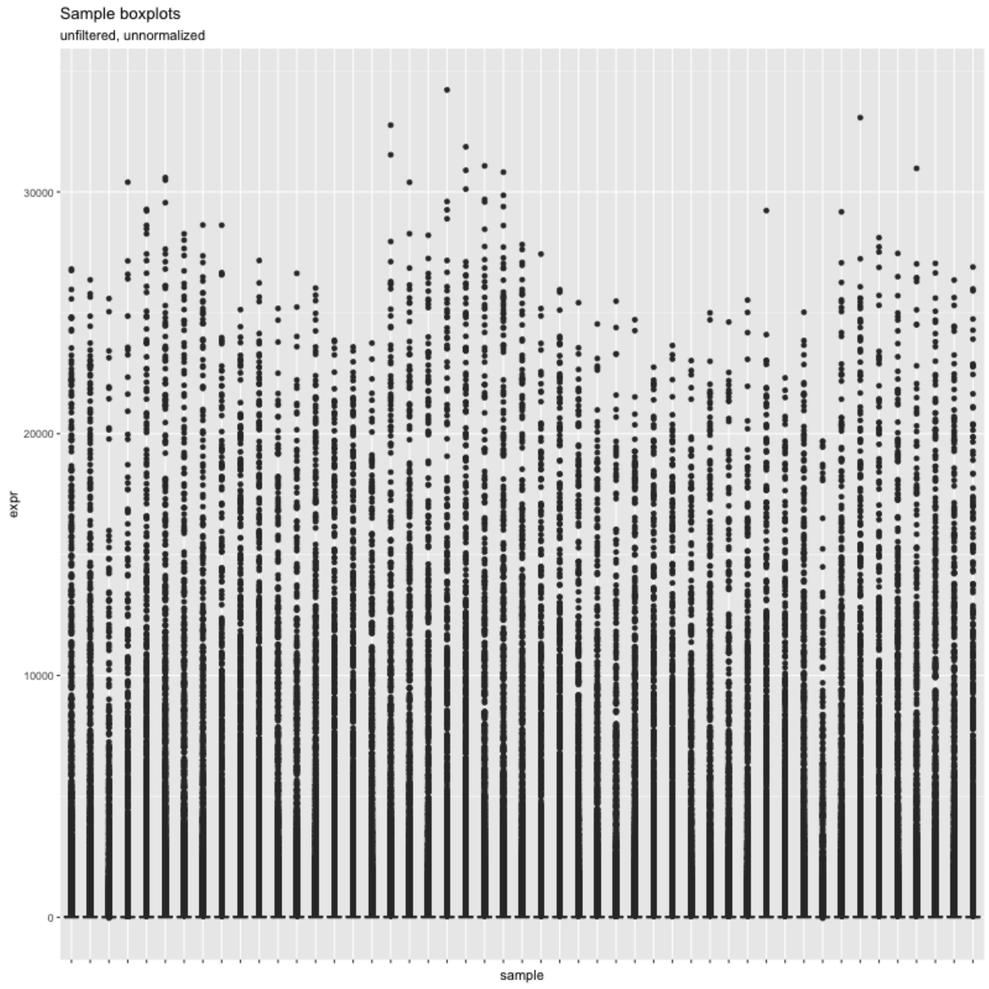


Unnormalized, unfiltered

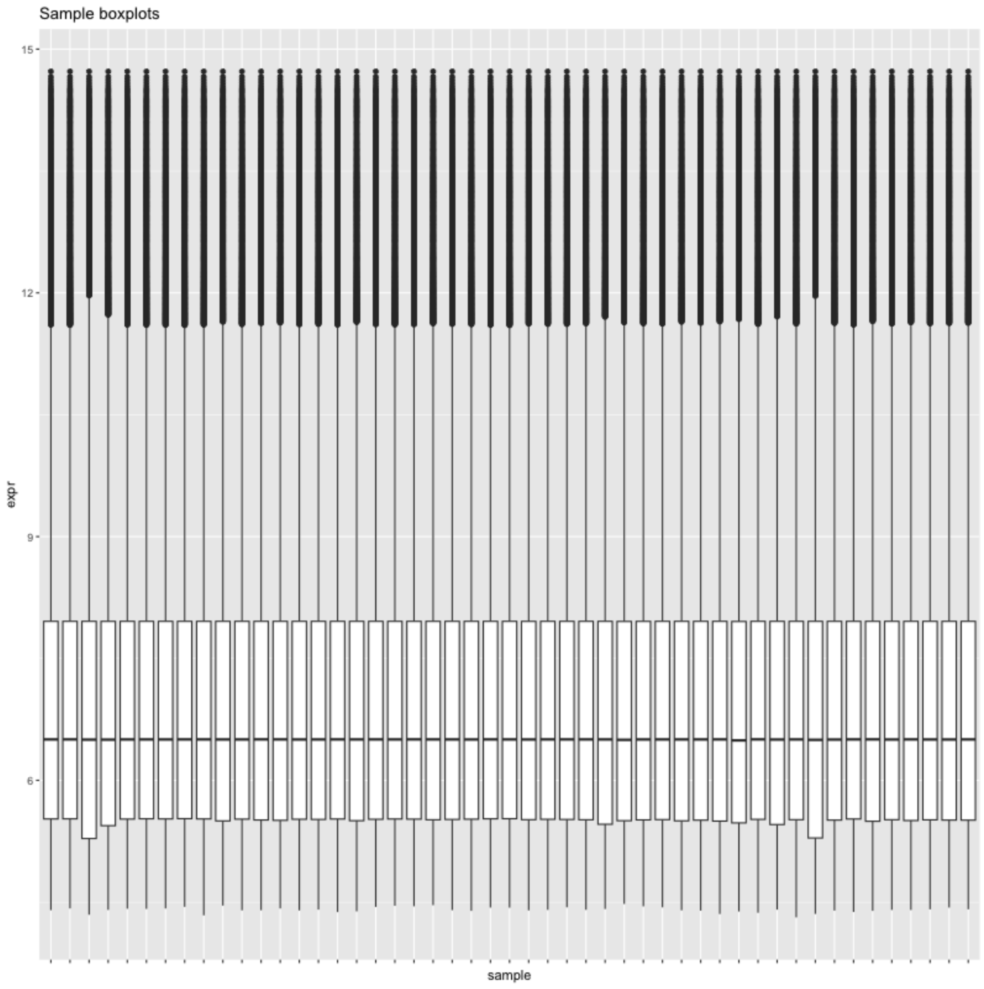


Normalized, filtered

The University of Texas at Austin
Waggoner Center for Alcohol & Addiction Research

### Obermoser: PCA (Test Finger)

Normalized and filtered.

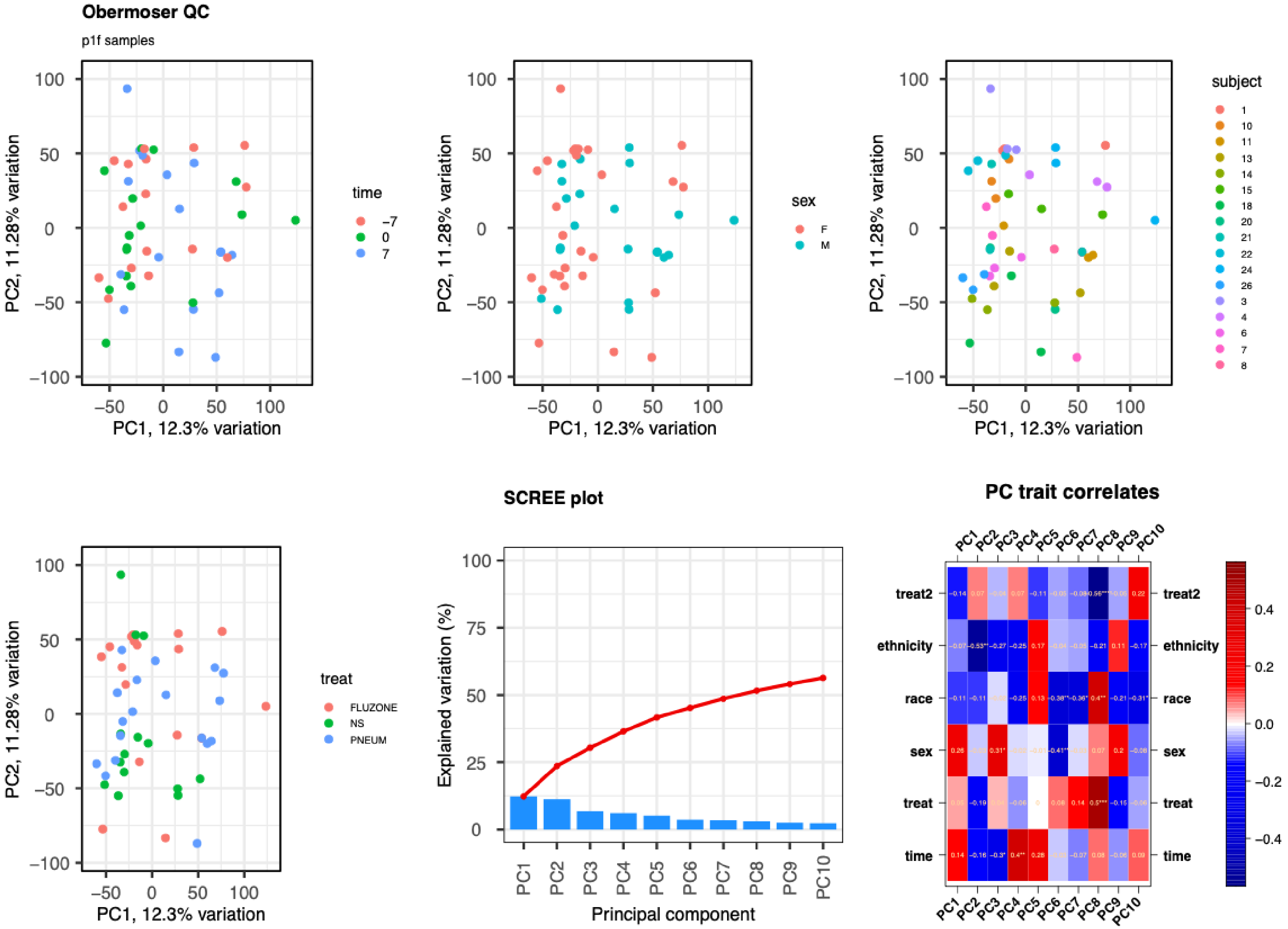

### Obermoser: PCA (Test_Finger)

Normalized and filtered.

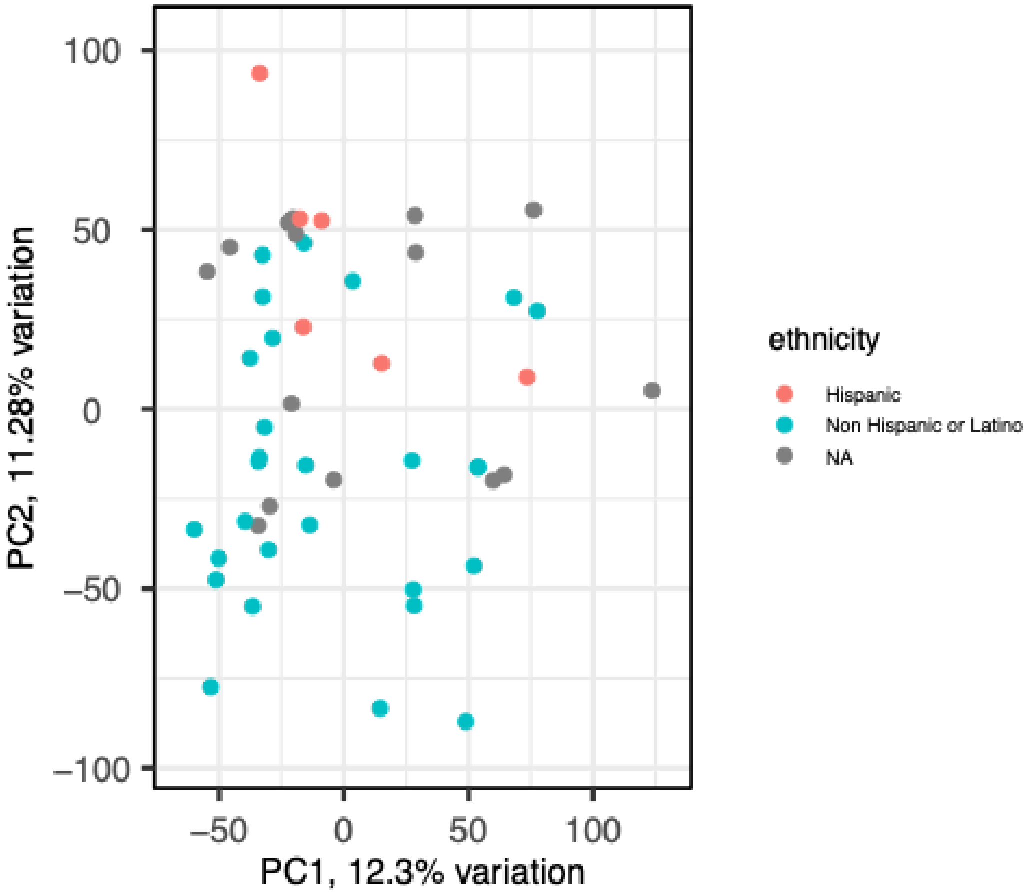


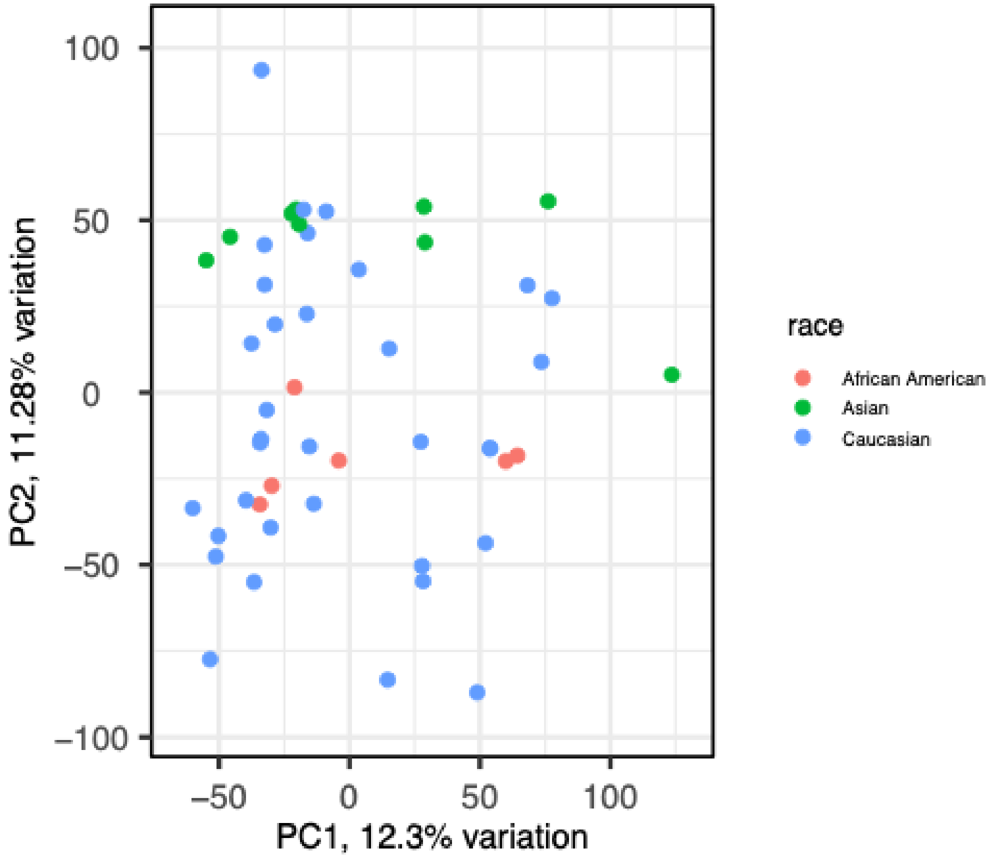

### Obermoser: Sample boxplots (Test_finger)

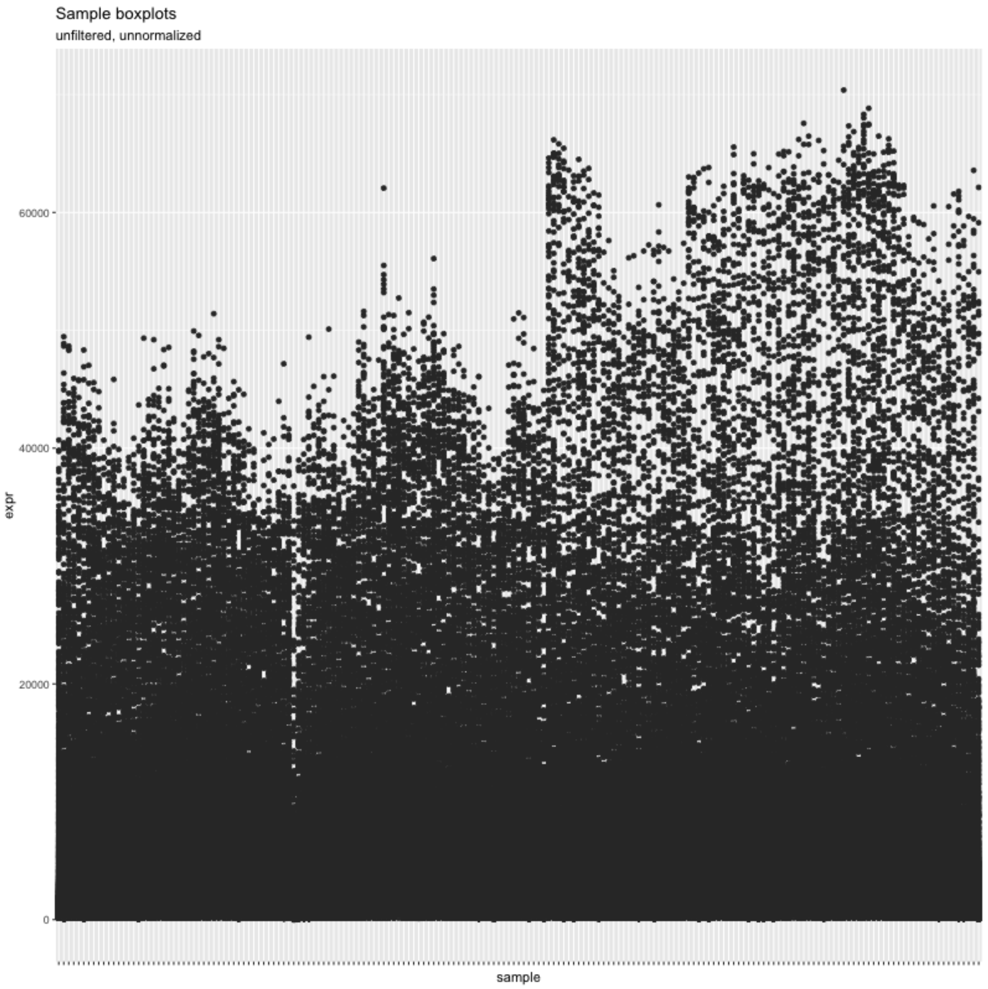


Unnormalized, unfiltered

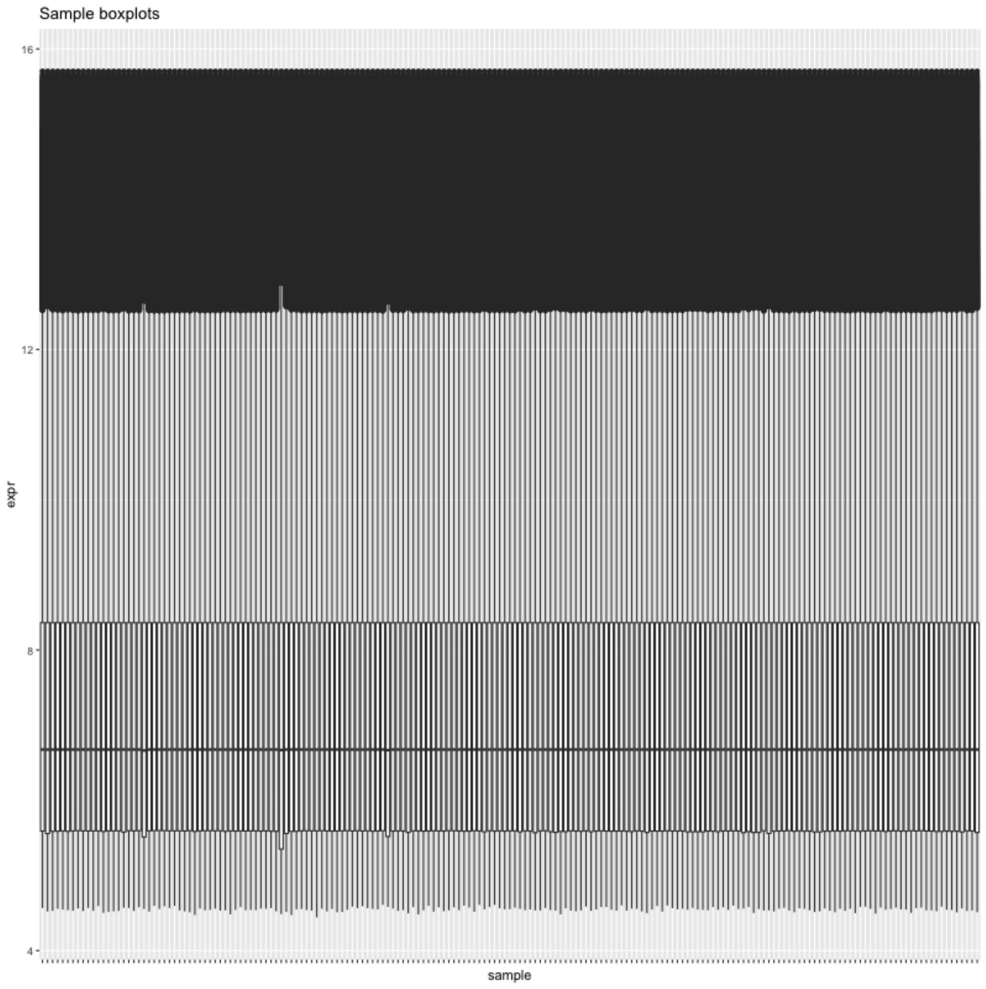


Normalized, filtered

### Affymetrix HGU133 Plus 2.0 Studies

### Dusek et al. (2008)

Microarray study, blood drawn at baseline and after a meditation training period.

Healthy adults with no previous experience with meditation were assigned an 8-week, self-administered meditation regiment designed to induce the "Relaxation Response," or RR. Changes in the transcriptome in response to increased ability to elicit the RR was measured before and after the training period.

Dusek, Jeffery A., Hasan H. Otu, Ann L. Wohlhueter, Manoj Bhasin, Luiz F. Zerbini, Marie G. Joseph, Herbert Benson, and Towia A. Libermann. "Genomic Counter-Stress Changes Induced by the Relaxation Response." *PLoS ONE* 3, no. 7 (July 2, 2008): e2576. https://doi.org/10.1371/journal.pone.0002576.

### Dusek et al. (2008) - summary

| Sample | Subject ID | Sex | Time Point | Reason Removed |
| --- | --- | --- | --- | --- |
| GSM253667 | 30 | Female | Baseline | Sex based on RPS4Y1 (201909_at) expression level was inconsistent with repeated measure. |
| GSM253695 | 30 | Male | Follow-up | Sex based on RPS4Y1 (201909_at) expression level was inconsistent with repeated measure. |
| GSM253709 | 50 | Male | Follow-up | Array quality concerns because of abnormal NUSE & RLE values, and pseudoimage. GSM253709 also clustered separately from other samples before filtering lowly expressed probes. |

The University of Texas at Austin
Waggoner Center for
Alcohol & Addiction Research

### Dusek: determination of sexes

- First, patient 30’s samples were removed because of mislabeling of the sample. (GSM253667 and GSM253695)
    - Because sex was not included in the study metadata, we imputed sex using a probe for RPS4Y1 (201909_at). These two samples did not agree in the sex of the patient.

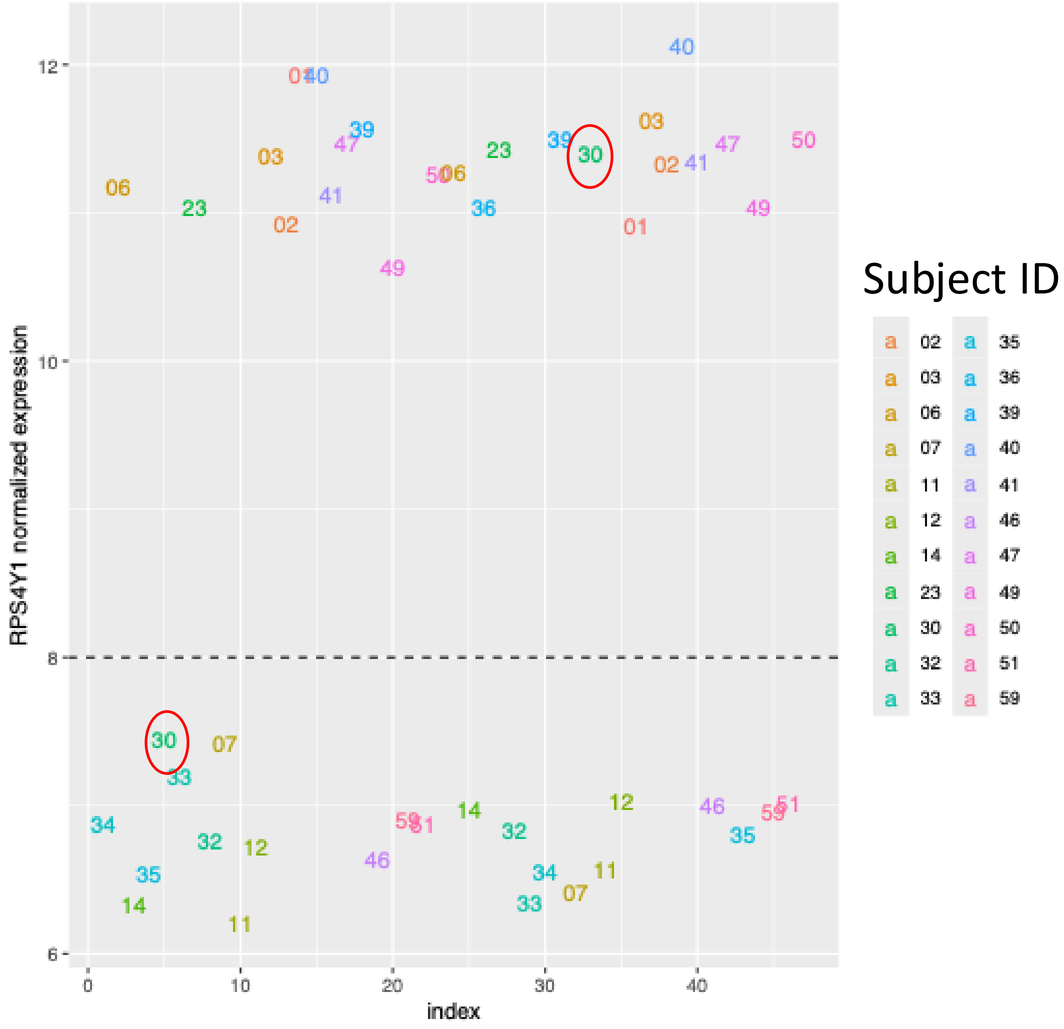

### Dusek: RLE, NUSE, and pseudoimages

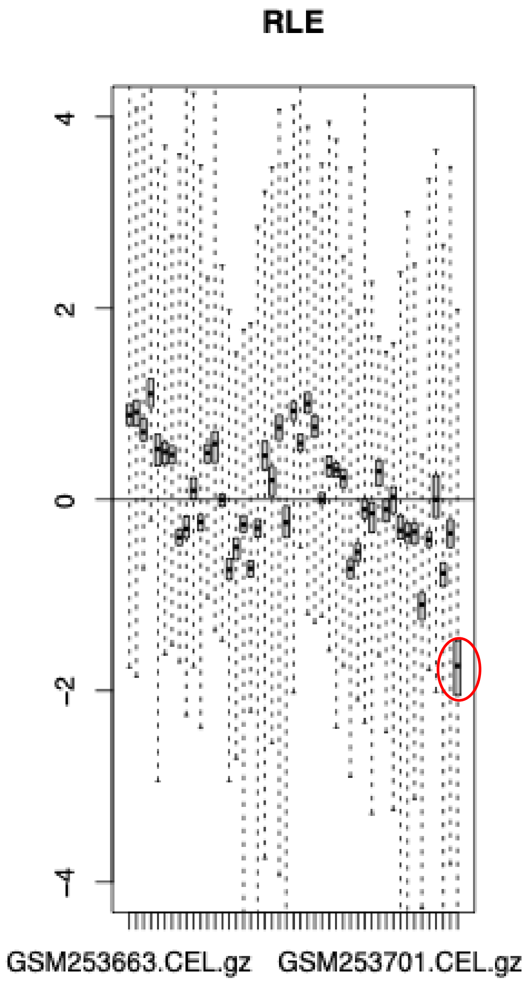


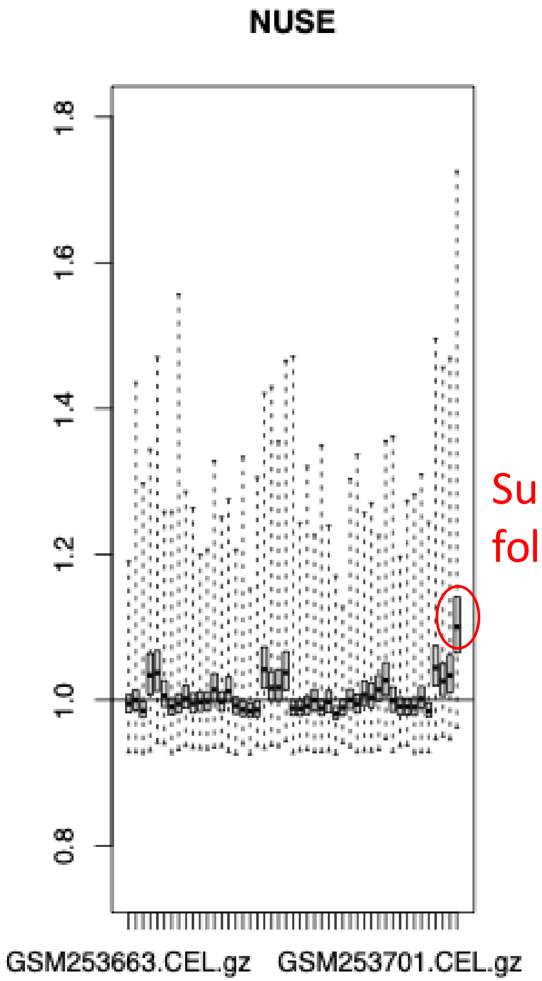


Subject 50,
follow-up

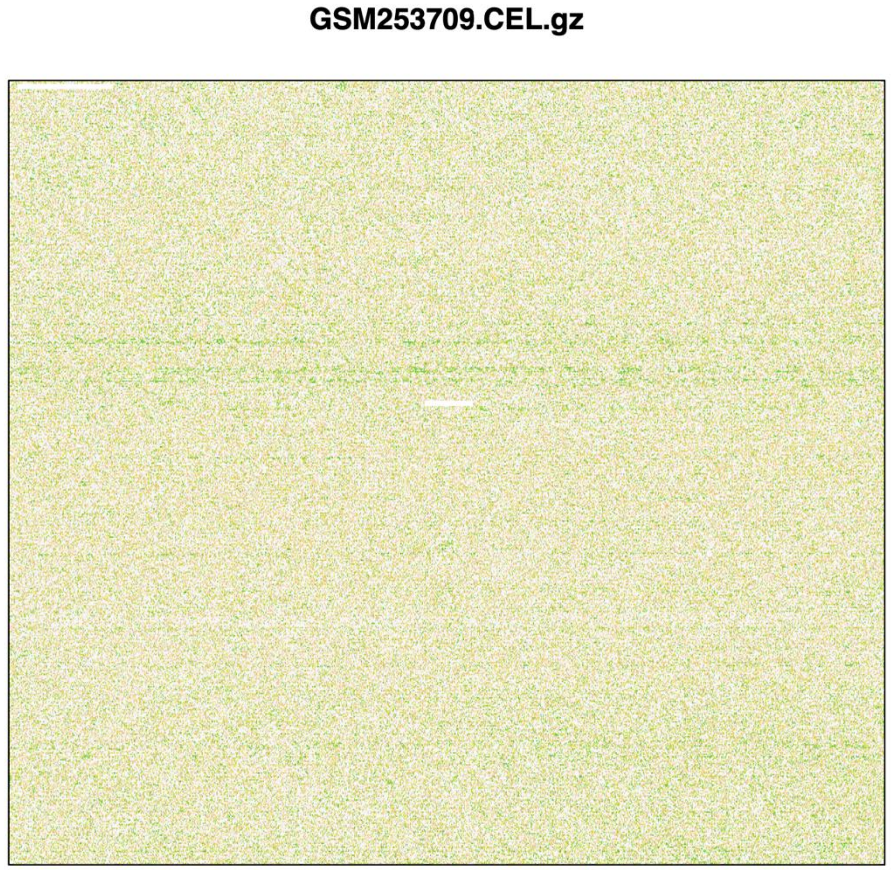


Subject 50 follow-up

The University of Texas at Austin
Waggoner Center for
Alcohol & Addiction Research

### PCA

- A PCA of the RMA background corrected and normalized data, but NOT filtered, shows that subject 50 at the follow-up time has abnormal probe values that cause it to vary from other samples.
    - This is not reflected in filtered data, so the biologically relevant genes may not be contributing to this variation. However, the technical variation due to noise after correction drove us to **remove this sample (GSM253709)** from the downstream analysis.

unfiltered

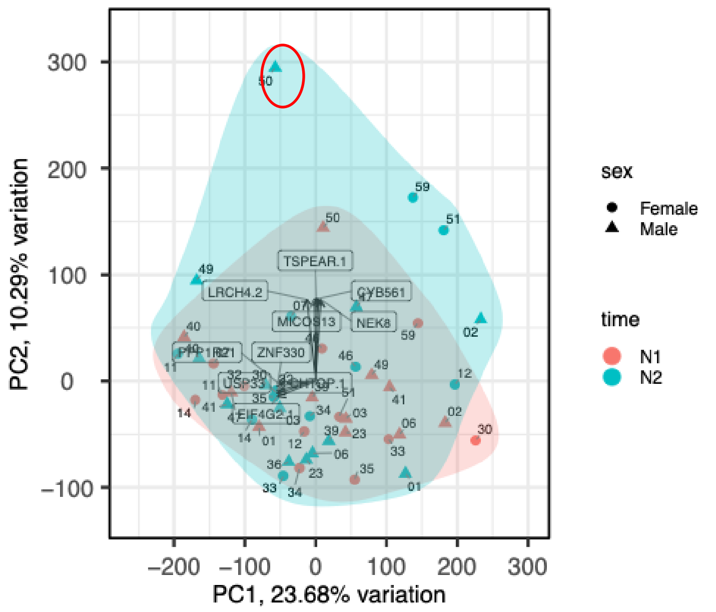


filtered

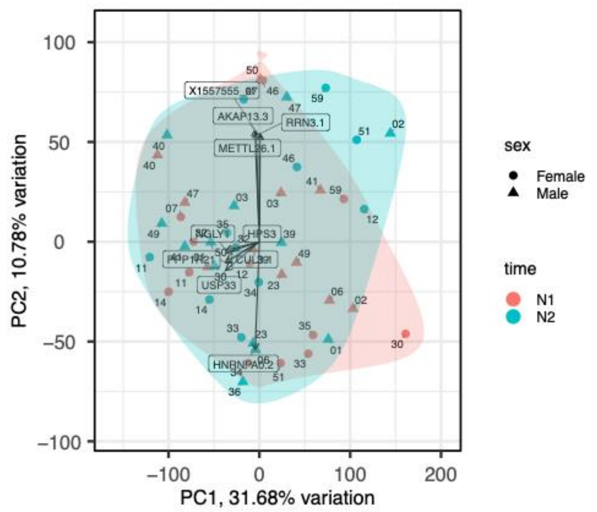

The University of Texas at Austin
Waggoner Center for
Alcohol & Addiction Research

### Dusek: Plots of features (L) and samples (R) after removal of GSM253667, GSM253695, and GSM253709

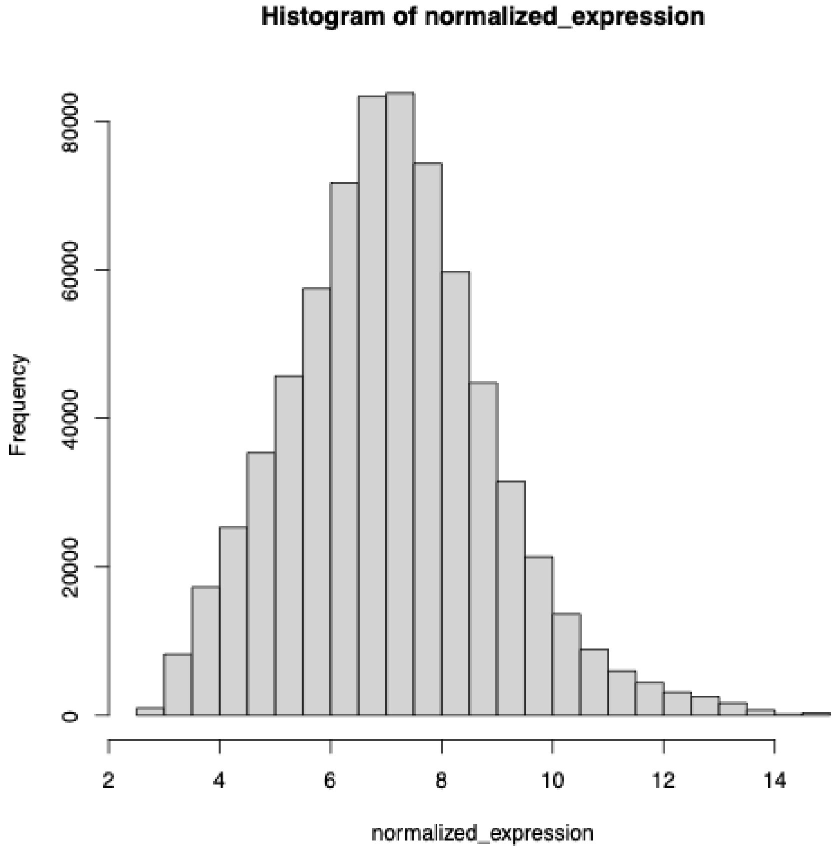


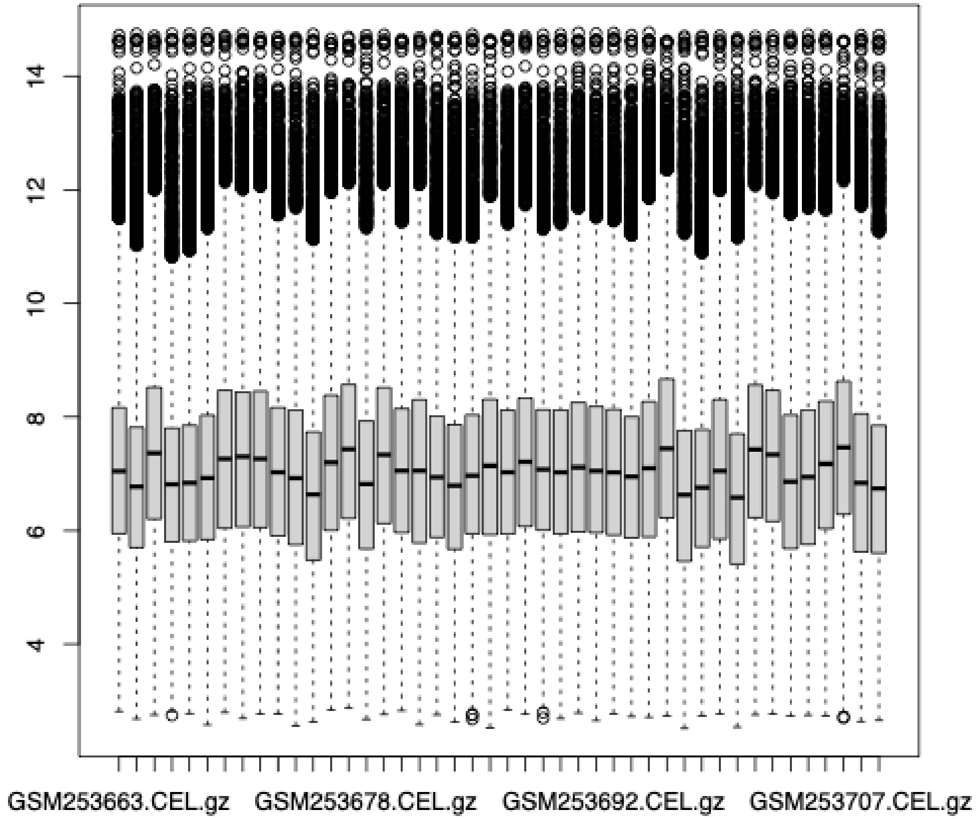

### Dusek: PCAtools analysis after removal of after removal of GSM253667, GSM253695, and **GSM253709**

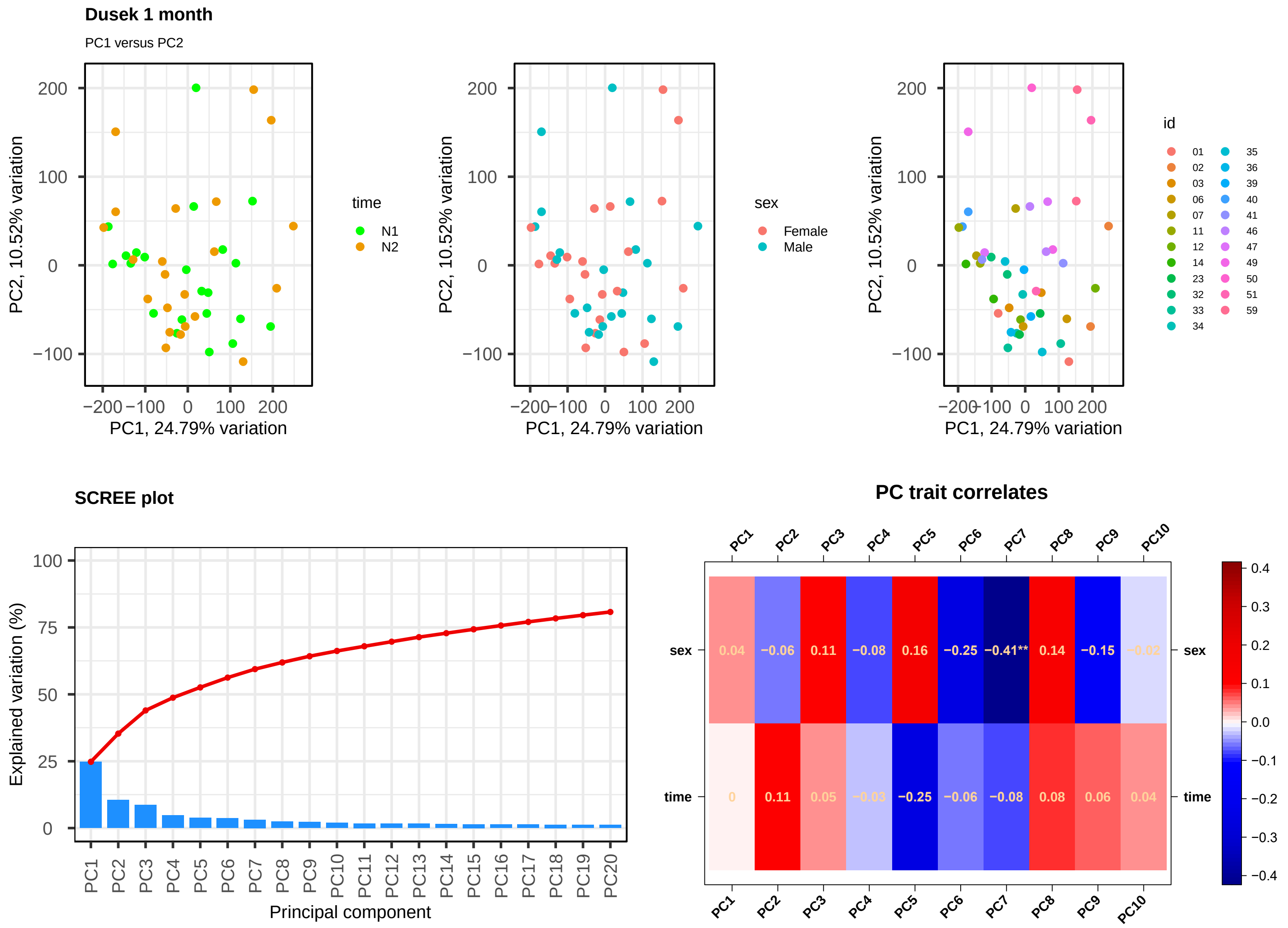

### Rusch et al. (2019)

Affymetrix HGU 133 Plus 2.0 microarray study, blood drawn at baseline and three months. The study aimed to compare military men who visited a clinic with complaints of sleep disturbance and stratified the men according to PTSD status. We filtered the data for men who did not have PTSD and for whom two samples were available (one baseline, one follow-up). Some men were clinically diagnosed with obstructive sleep apnea, while others did not receive a clinical diagnosis.

Rusch, Heather L., Jeffrey Robinson, Sijung Yun, Nicole D. Osier, Christiana Martin, Chris R. Brewin, and Jessica M. Gill. "Gene Expression Differences in PTSD Are Uniquely Related to the Intrusion Symptom Cluster: A Transcriptome-Wide Analysis in Military Service Members." *Brain, Behavior, and Immunity* 80 (August 1, 2019): 904–8. https://doi.org/10.1016/j.bbi.2019.04.039.

### Rusch et al. (2019) - summary

| Sample | Subject ID | Sex | Time Point | Reason Removed |
|---|---|---|---|---|
| GSM217526 | 108 | Male | Follow-up | This sample had abnormal NUSE and RLE statistics, artifacts on pseudoimage, and did not cluster with other samples in the experiment on PCA with all probes included. |
| GSM2175214 | 76 | Male | Baseline | This sample artifacts on pseudoimage and did not cluster with other samples in the experiment on PCA before or after filtering lowly-expressed genes. |

The University of Texas at Austin
Waggoner Center for
Alcohol & Addiction Research

### Rusch: RLE, NUSE, and pseudoimages

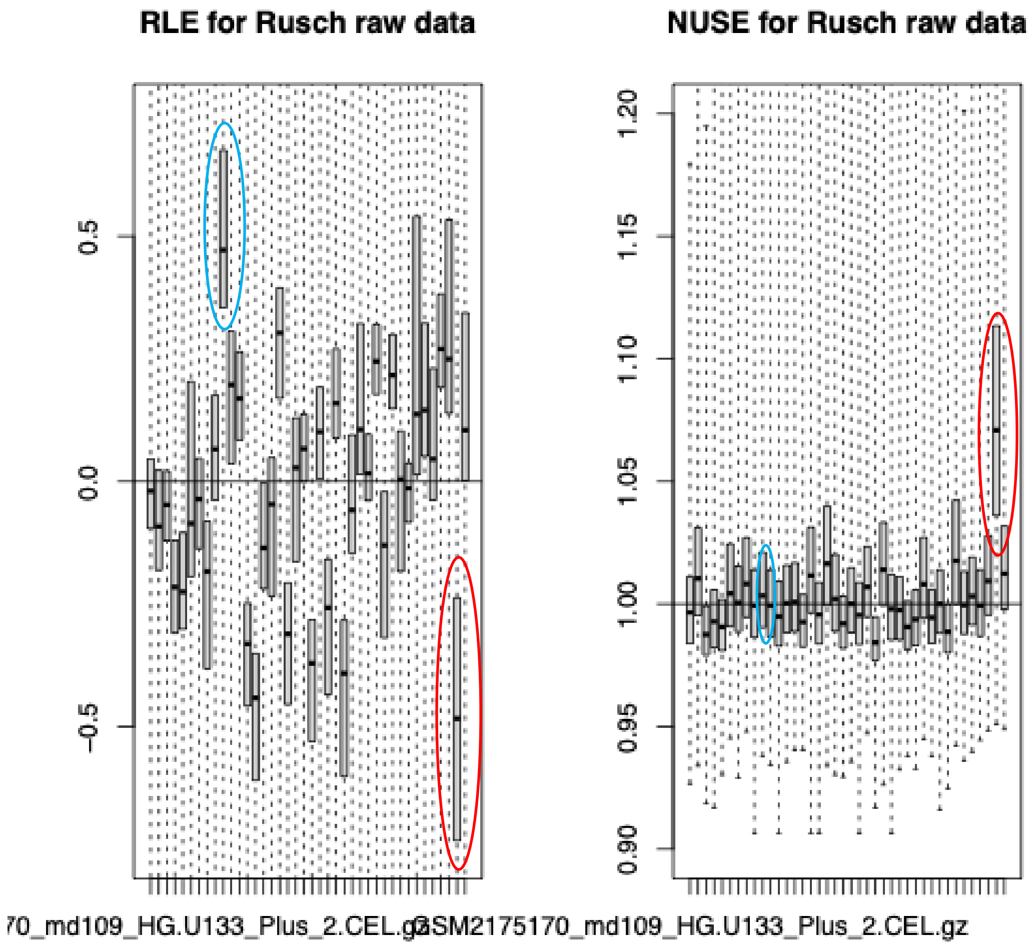


GSM2175202

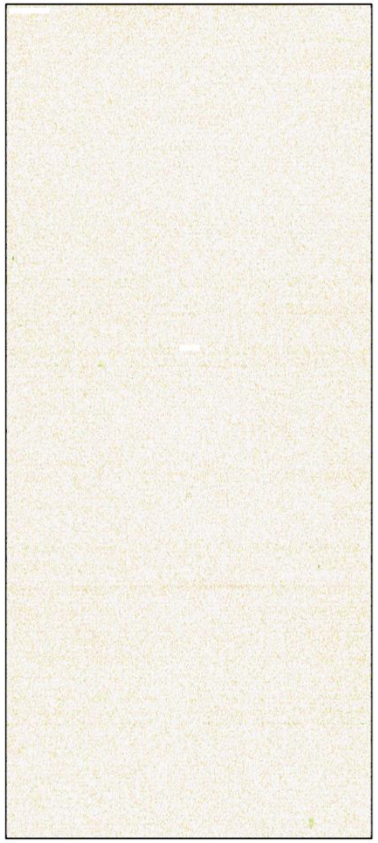

Subject 38, timepoint 2

GSM2175261

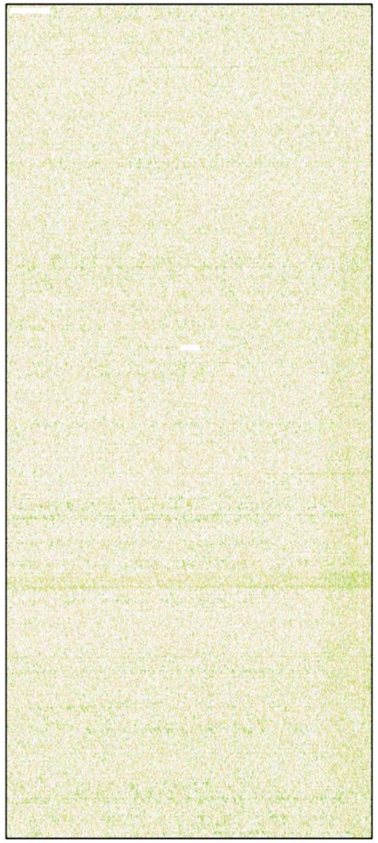

Subject 108, timepoint 2

The University of Texas at Austin
Waggoner Center for
Alcohol & Addiction Research

### Rusch: Initial PCA

Although GSM2175202 had relatively high RLE values, its NUSE values and pseudoimage were not concerning, and this array clustered with other samples in a PCA plot. So, we did not remove it from the analysis.

**GSM2175261** (patient 108, follow-up sample) was removed because of abnormal NUSE statistics and its separation from the experiment's cluster. We removed this sample and renormalized the data for further analysis.

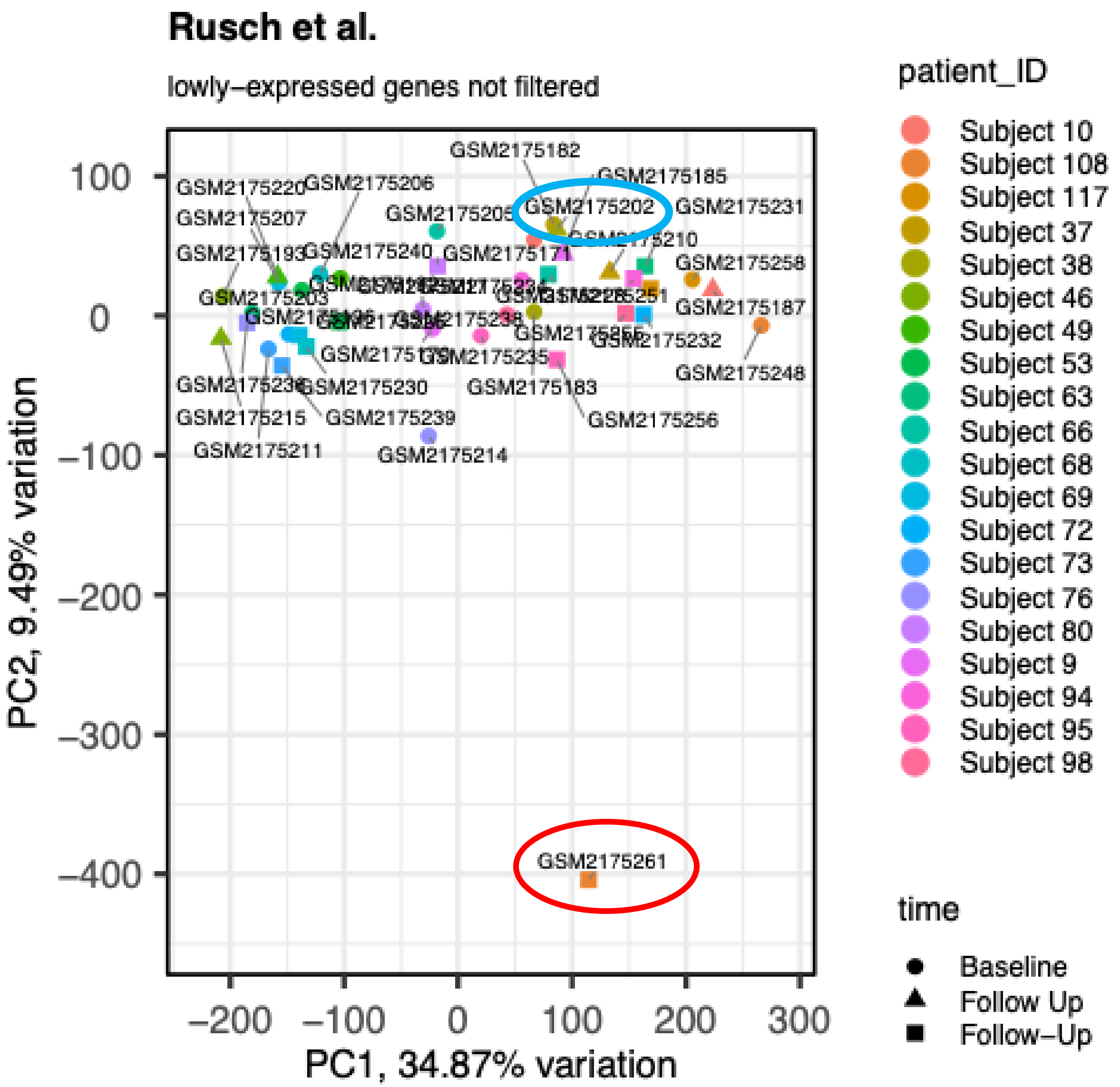

The University of Texas at Austin
Waggoner Center for
Alcohol & Addiction Research

### Rusch: Unfiltered vs. filtered PCA

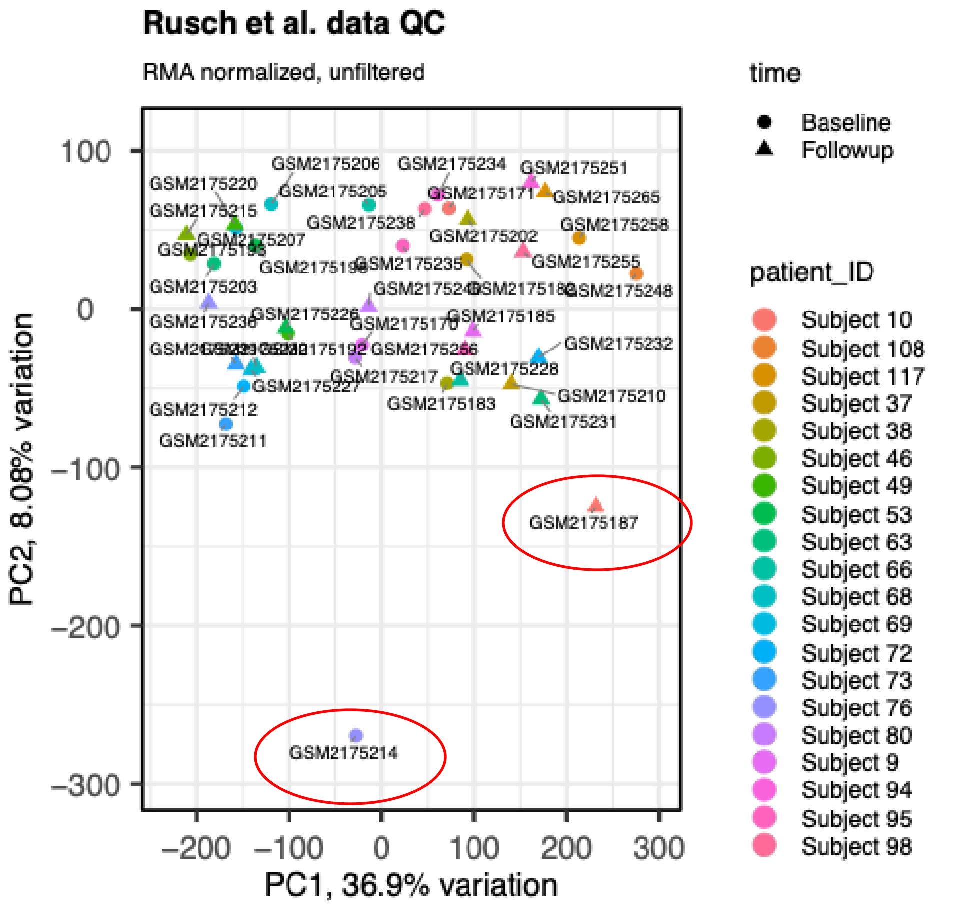


*Filter lowly-expressed genes*

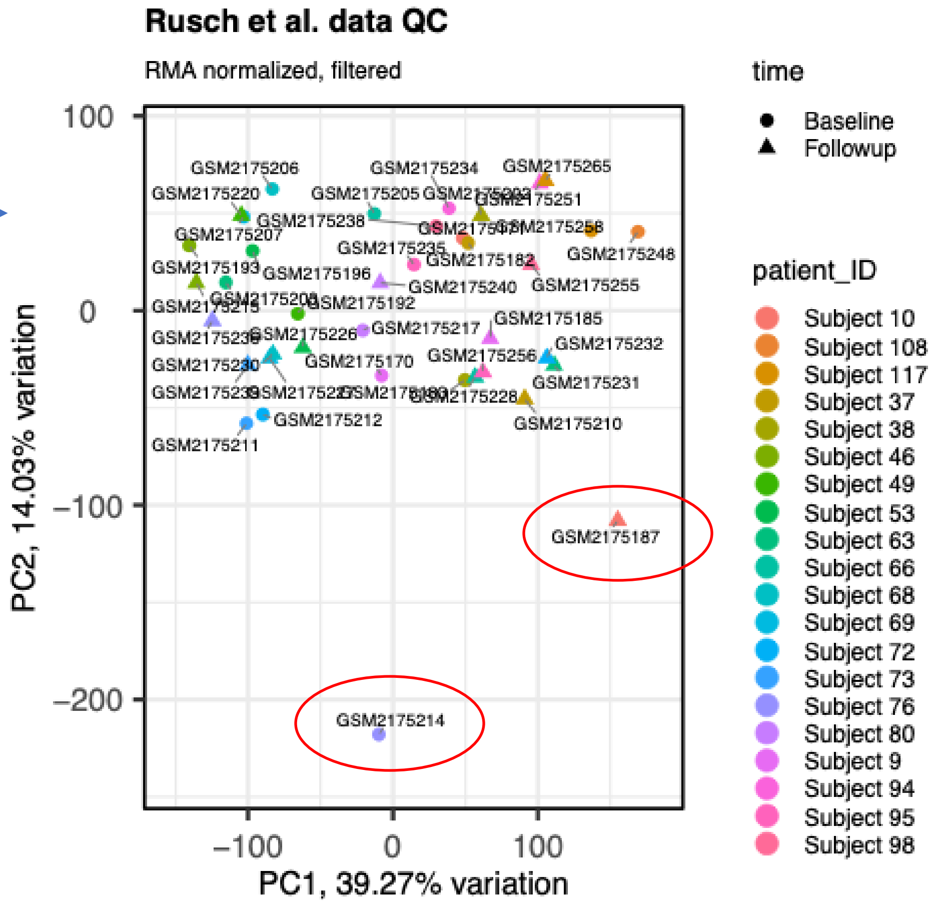


*Note: GSM217526 was already removed.*

The University of Texas at Austin
Waggoner Center for
Alcohol & Addiction Research

### Rusch: pseudoimages

GSM2175187

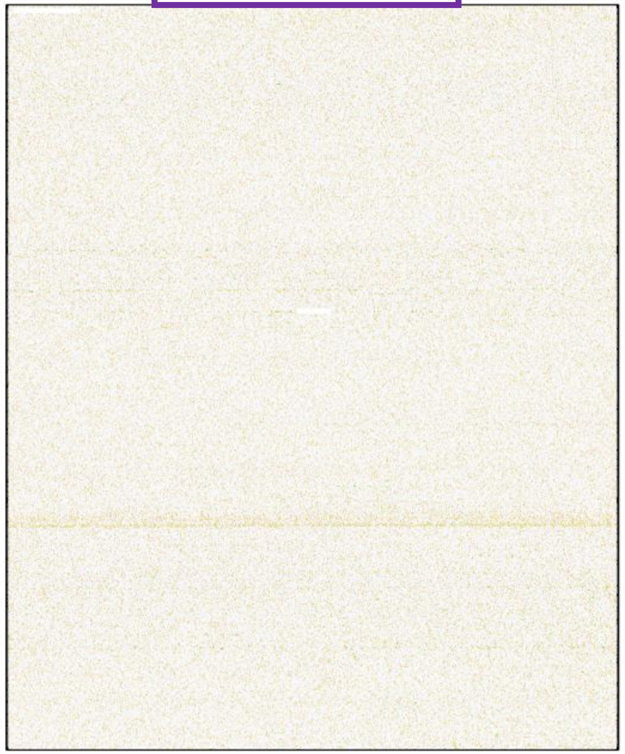

Subject 10, timepoint 2.

GSM2175214

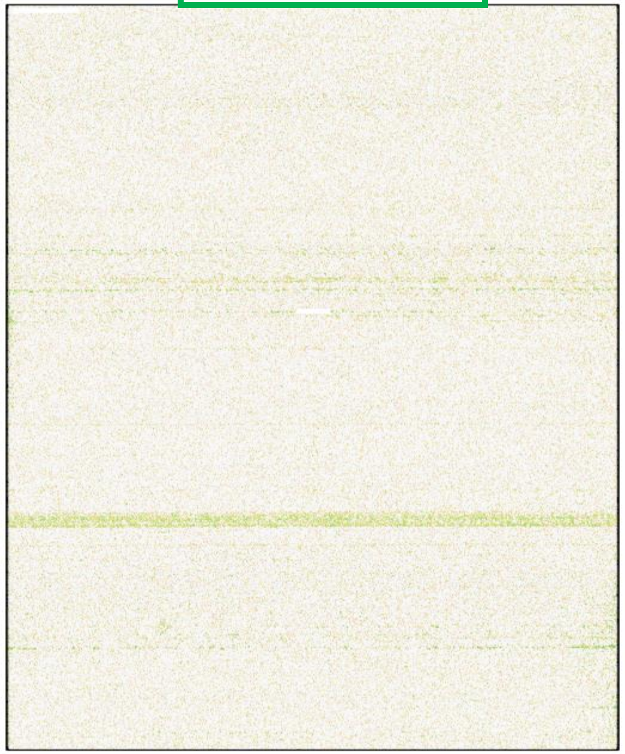

Subject 76, timepoint 1.
**Sample removed.**

We removed sample GSM2175214 because it is a clear outlier sample on PCA and has a visible artifact on pseudoimage (right).

While GSM2175187 (left) has similar features as those stated for GSM2175214, these deviations are not as extreme. For this reason, it is harder to judge whether these differences in the GSM2175187 sample are cause for concern. We chose to keep GSM2175187 because its RLE and NUSE values were normal and because the PCA plots do not clearly indicate that this sample is an outlier.

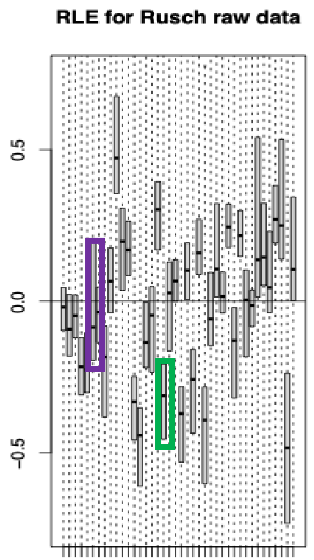


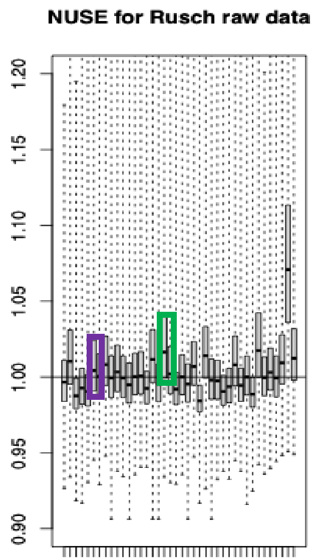

The University of Texas at Austin
Waggoner Center for
Alcohol & Addiction Research

### Rusch: After removing GSM2175214

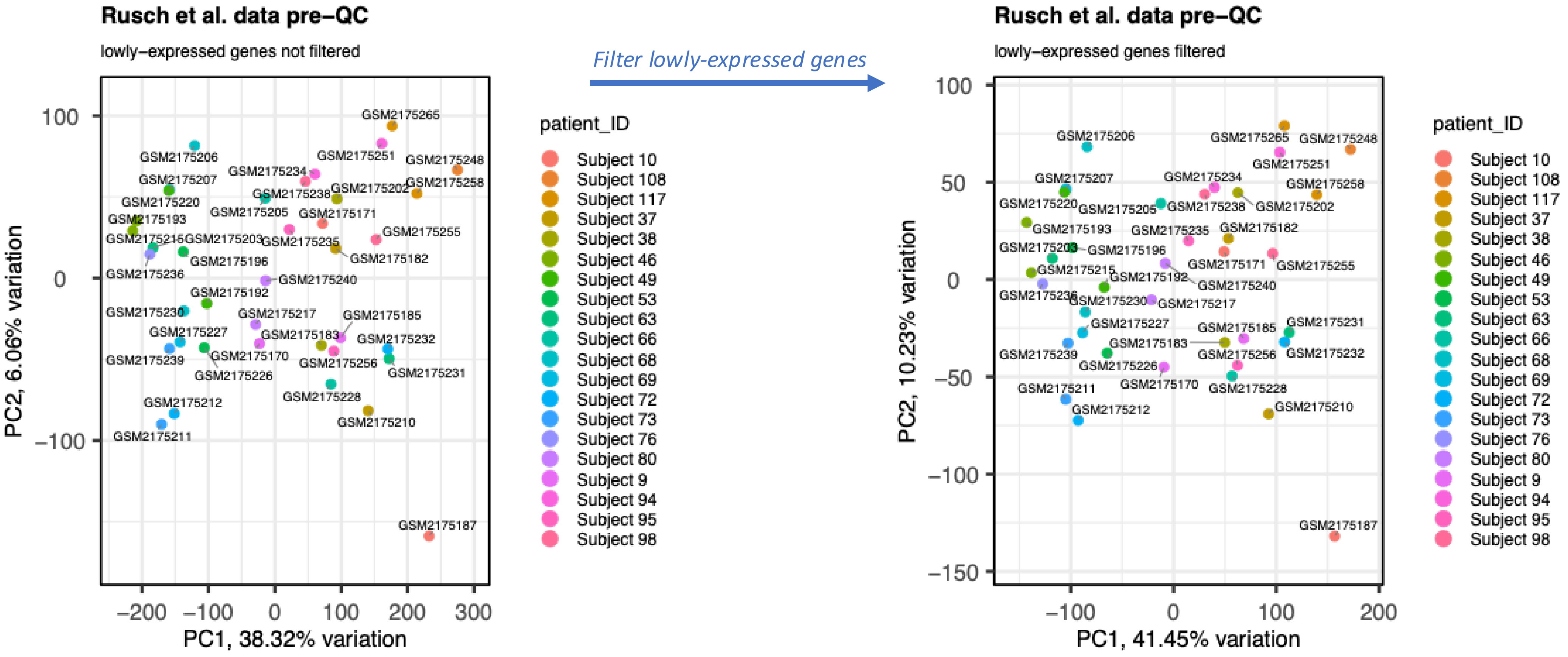

The University of Texas at Austin
Waggoner Center for
Alcohol & Addiction Research

### Rusch: Plots of features (L) and samples (R) after removing GSM2175261 and GSM2175214

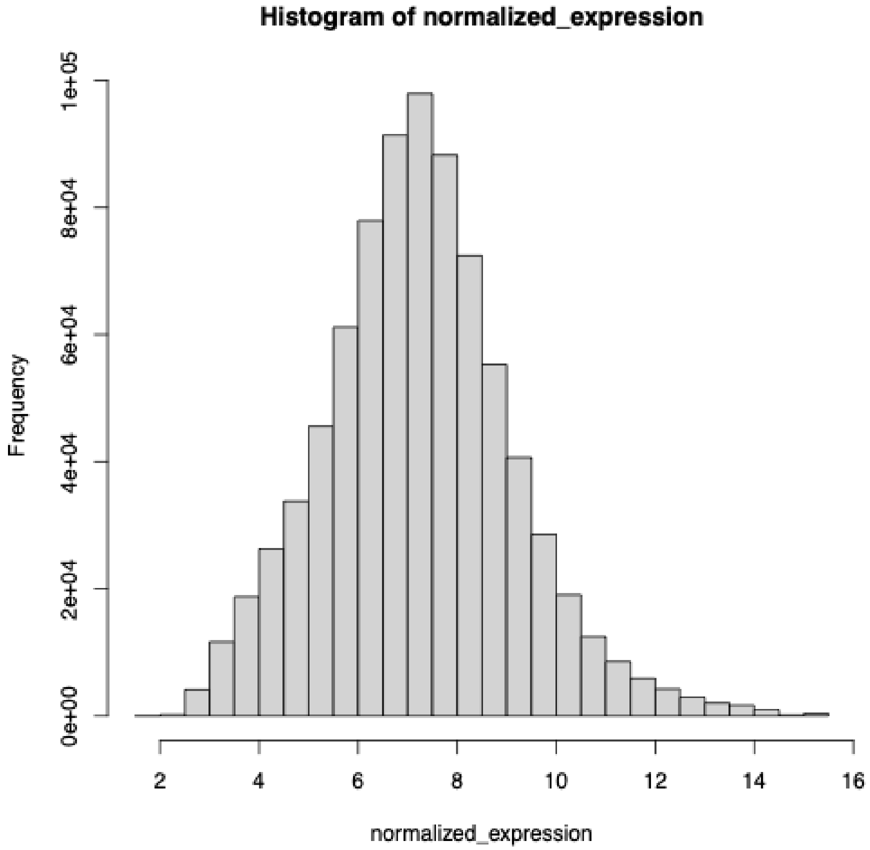


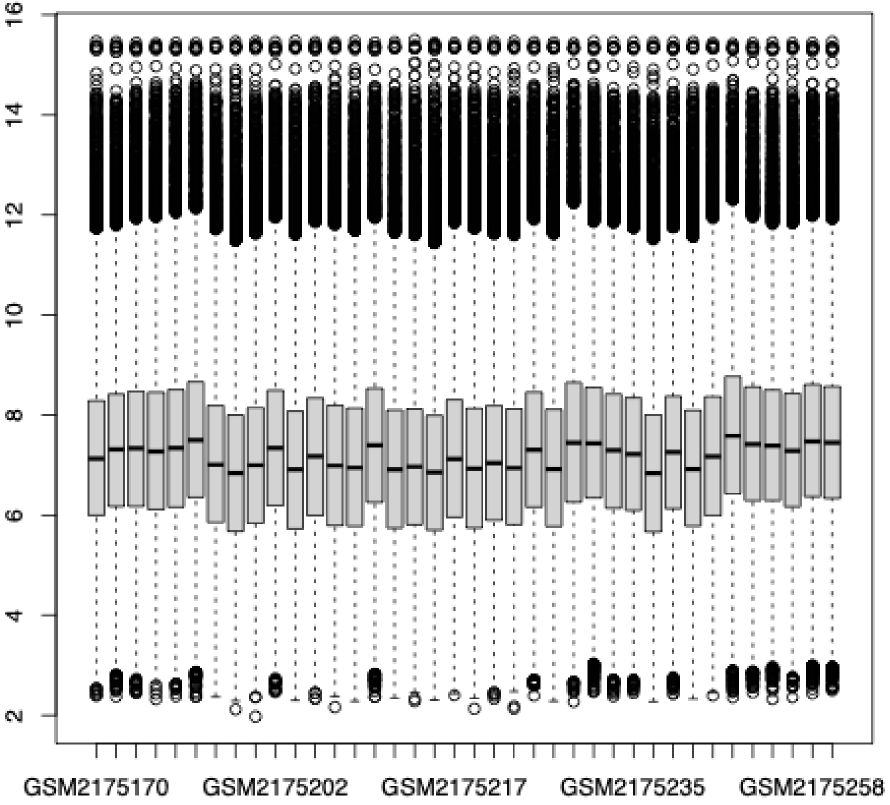

The University of Texas at Austin
Waggoner Center for
Alcohol & Addiction Research

### Rusch: PCA After removing GSM2175621 and GSM2175214

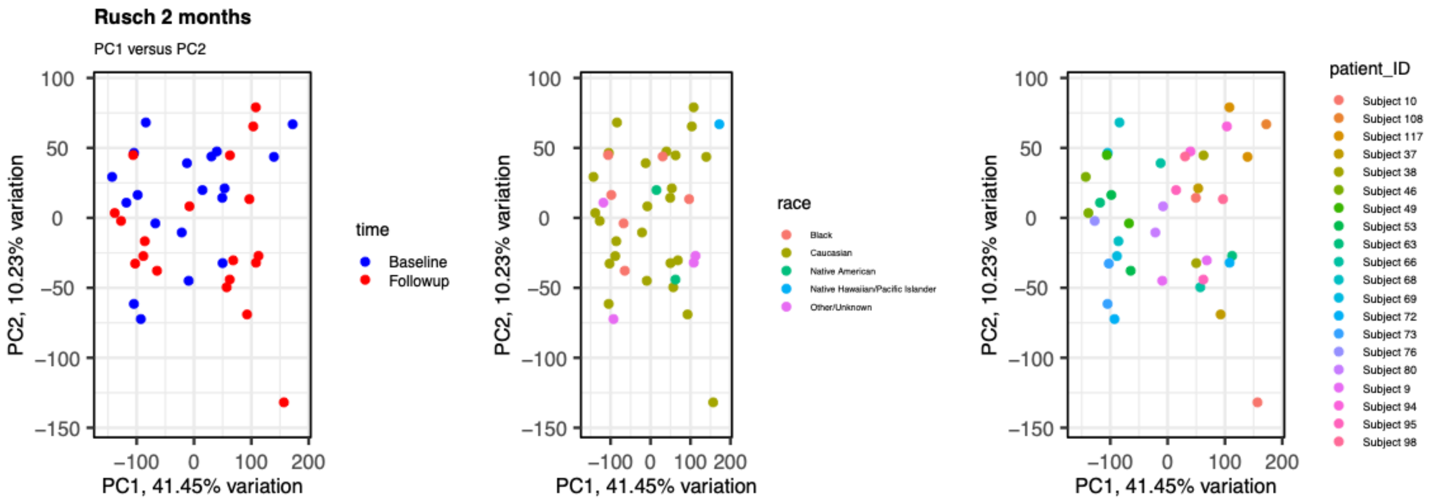


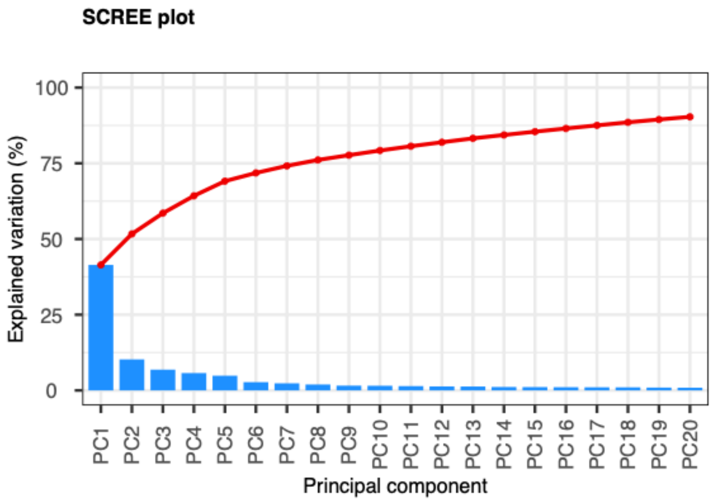


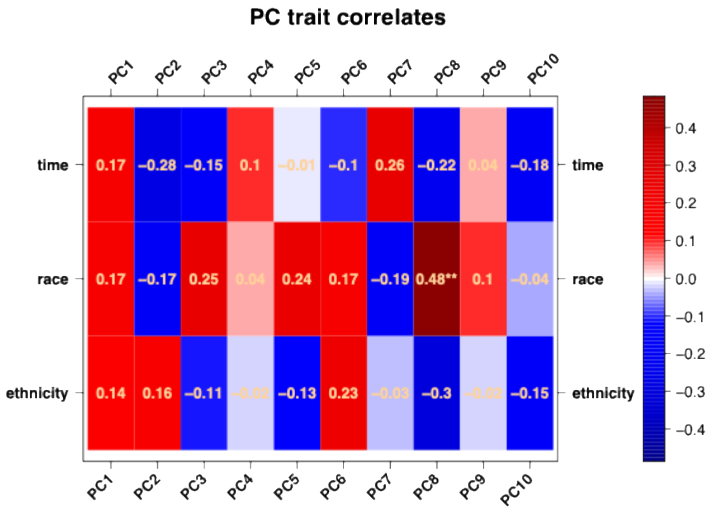

The University of Texas at Austin
Waggoner Center for
Alcohol & Addiction Research

### Meaburn et al. (2009)

Microarray study, blood was drawn at baseline and 4 hours, repeated ten months later. All samples come from sets of twins, who were twelve years of age at the time of initial sample collection.

Meaburn, Emma L., Cathy Fernandes, Ian W. Craig, Robert Plomin, and Leonard C. Schalkwyk. "Assessing Individual Differences in Genome-Wide Gene Expression in Human Whole Blood: Reliability Over Four Hours and Stability Over 10 Months." *Twin Research and Human Genetics : The Official Journal of the International Society for Twin Studies* 12, no. 4 (August 2009): 10.1375/twin.12.4.372. https://doi.org/10.1375/twin.12.4.372.

### Meaburn et al. (2009) - summary

- No samples were removed

### Meaburn: control probes

Although the publication did not indicate whether the samples were processed and hybridized to arrays in separate batches, the differences in control probes plotted here suggest that the samples collected on the first day were hybridized separately from the samples collected 10 months later. For this reason, we performed all analyses separately for the first day (Day 1) and second day (Day 2).

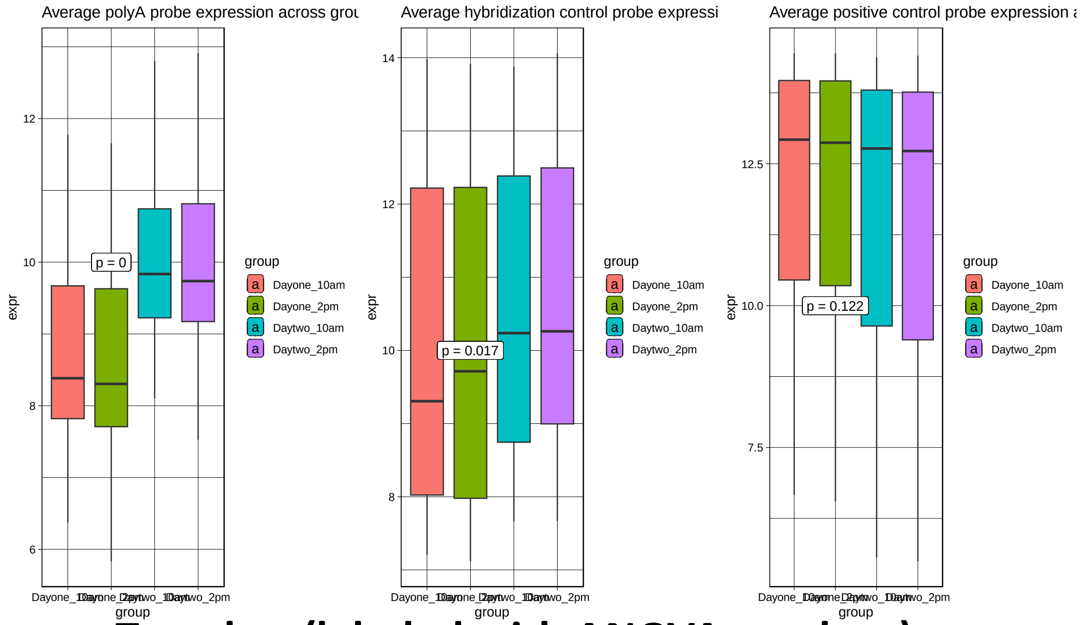


Together (labeled with ANOVA p-values)

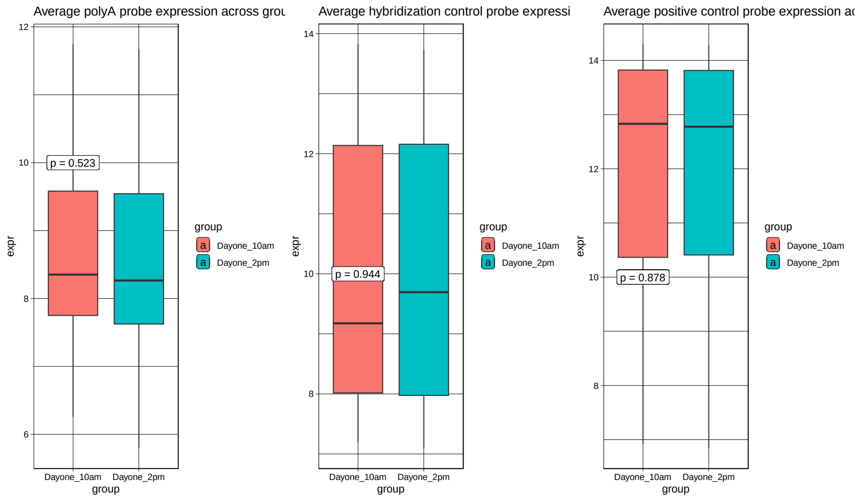


Day 1

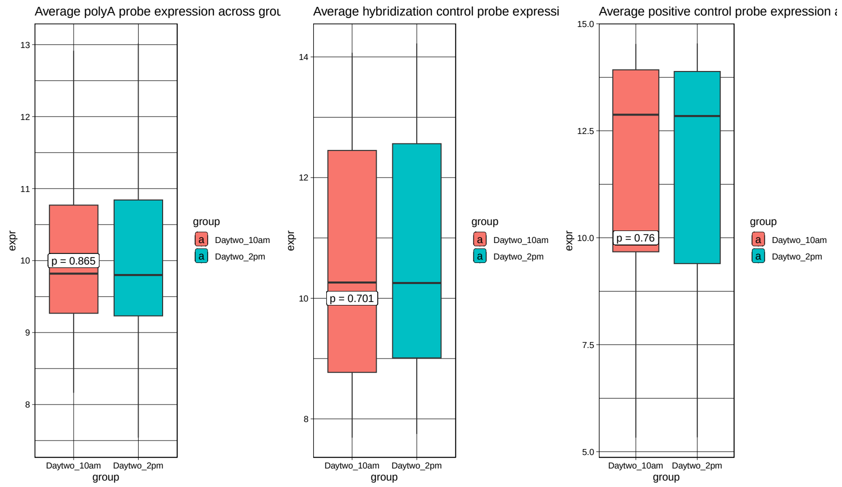


Day 2

(t-test)

The University of Texas at Austin
Waggoner Center for
Alcohol & Addiction Research

### Meaburn: RLE and NUSE for days 1 and 2

None of the arrays pre-normalization have major deviations in RLE, and NUSE statistics look good.

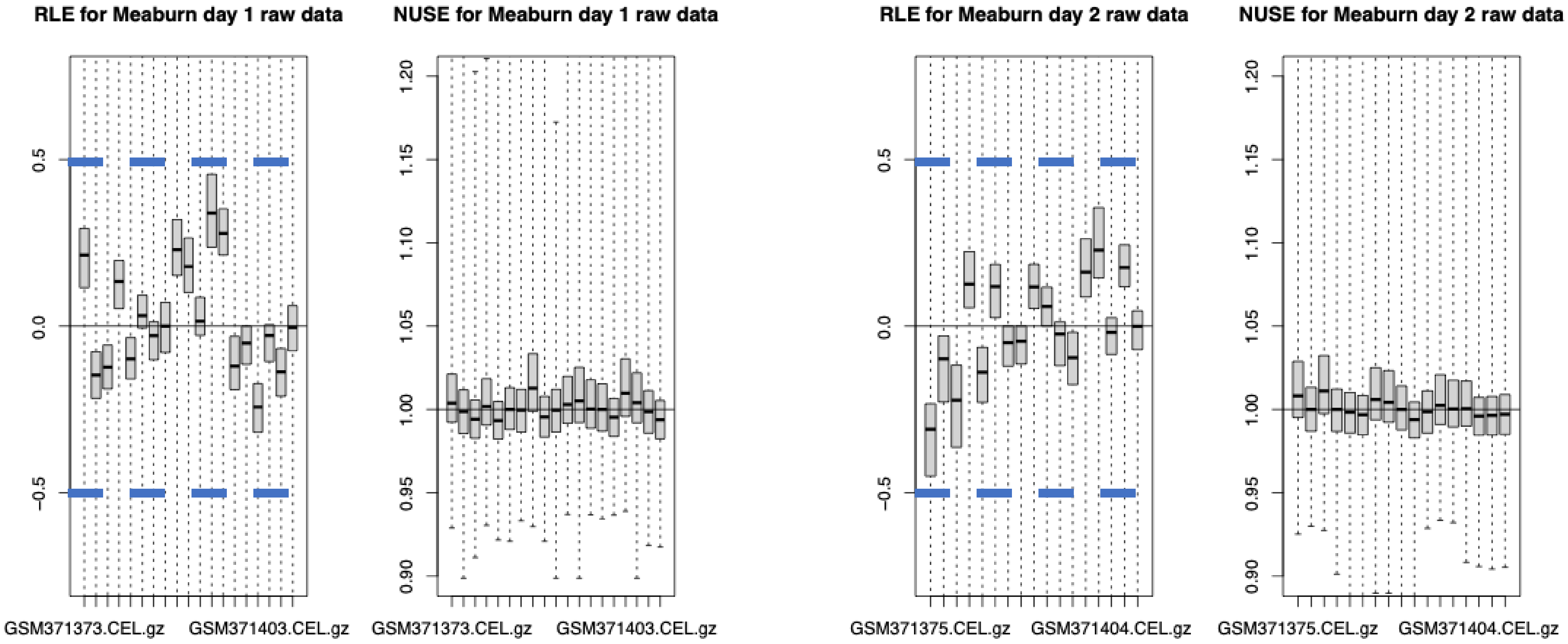


**DAY 1** **DAY 2**

The University of Texas at Austin
Waggoner Center for
Alcohol & Addiction Research

### Meaburn: Plots of features and samples

**Day 1**

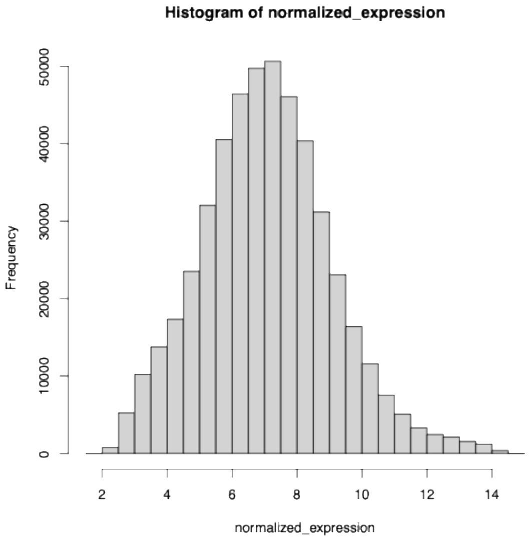


**Features**

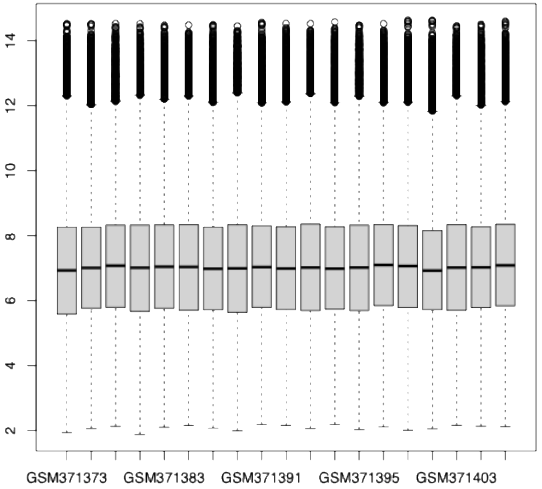


**Samples**

**Day 2**

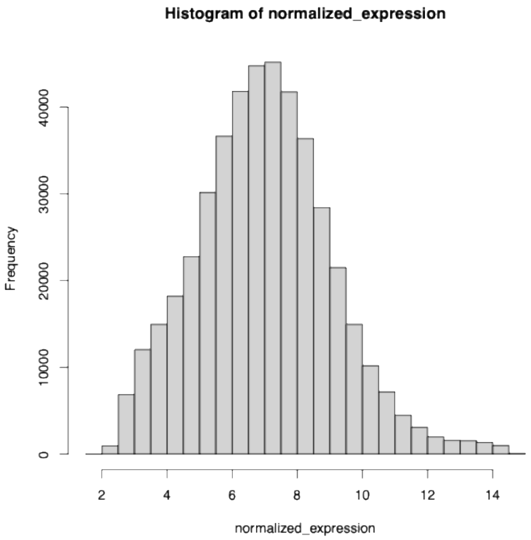


**Features**

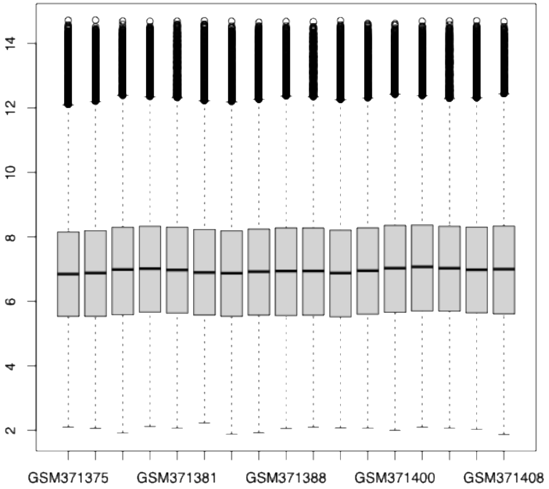


**Samples**

The University of Texas at Austin
Waggoner Center for
Alcohol & Addiction Research

### Meaburn: day 1 PCA before filtering genes

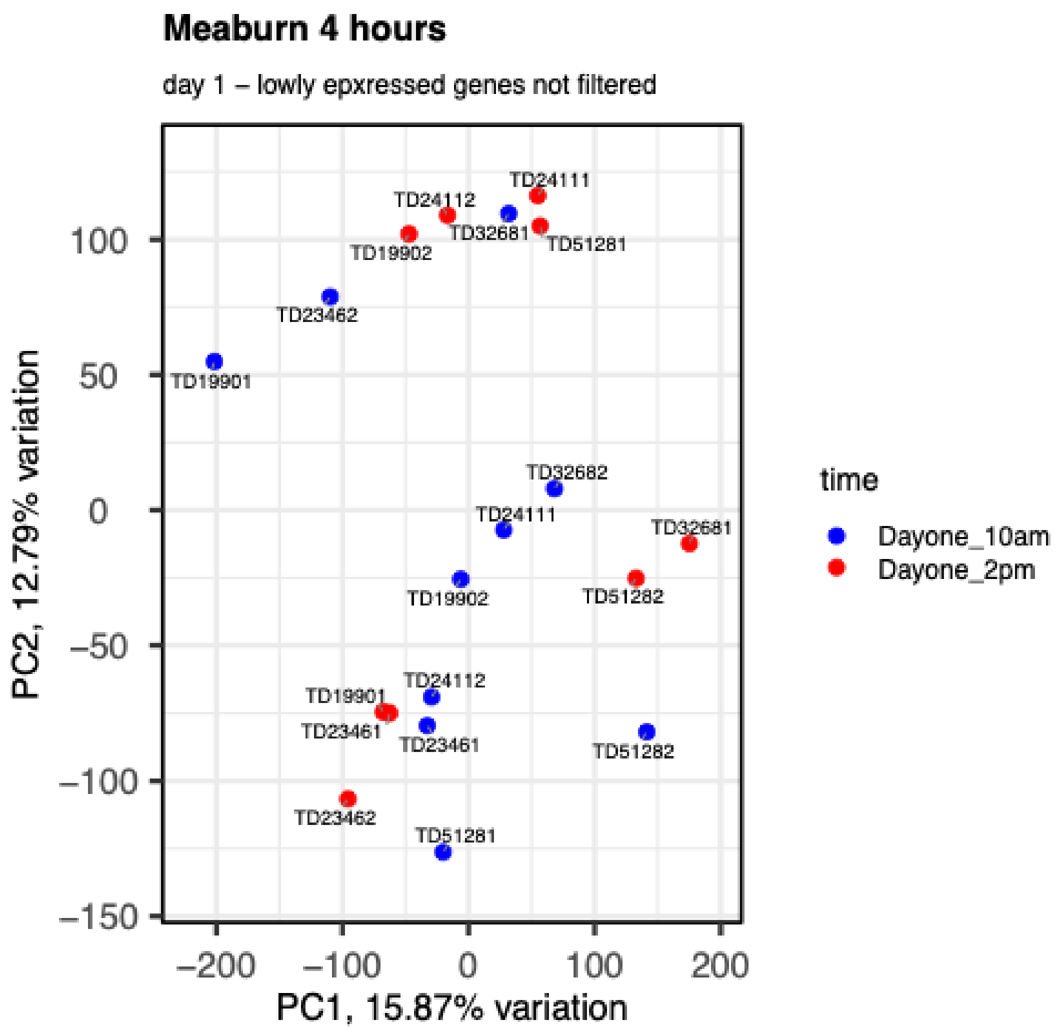

### Meaburn: day 1 PCA after filtering genes

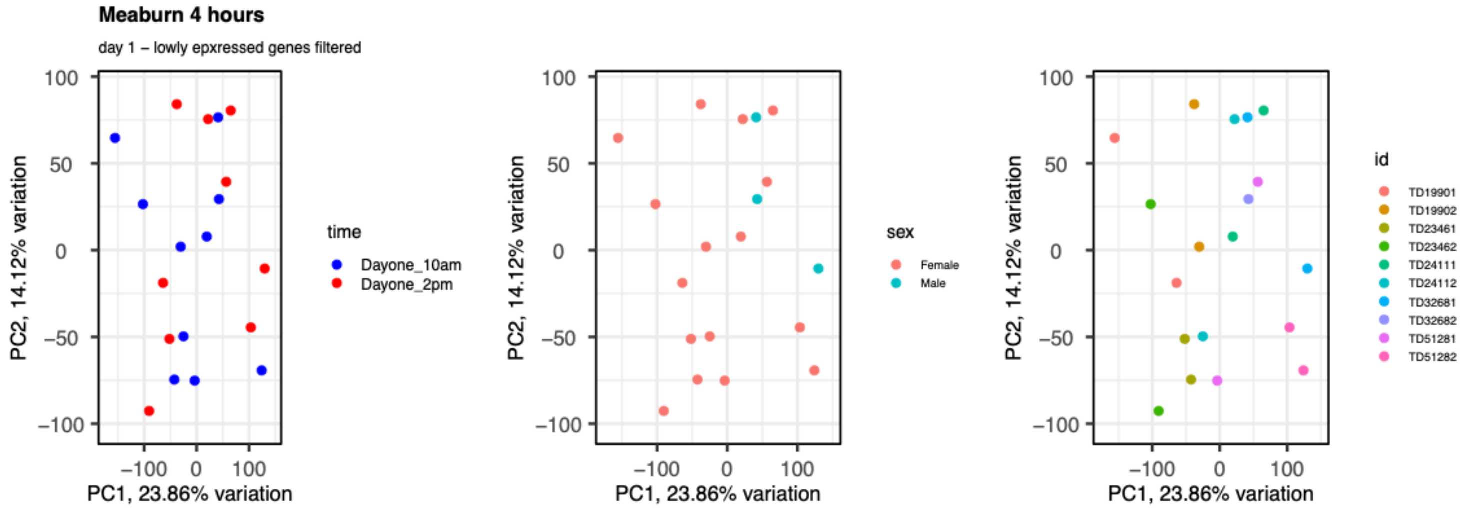


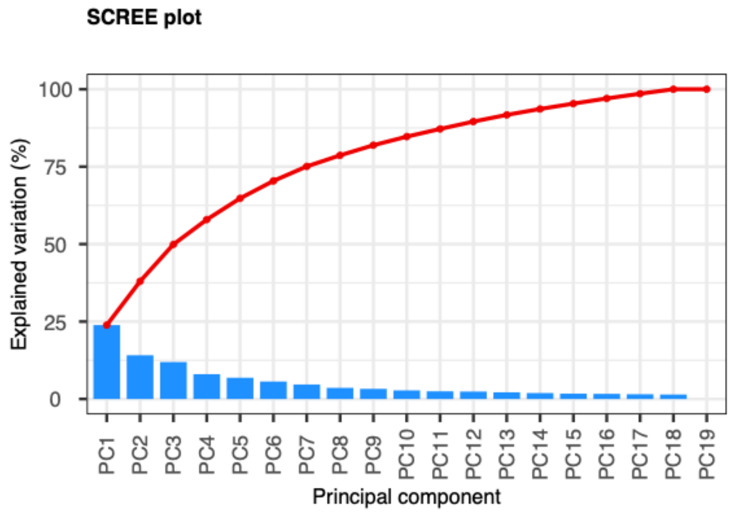


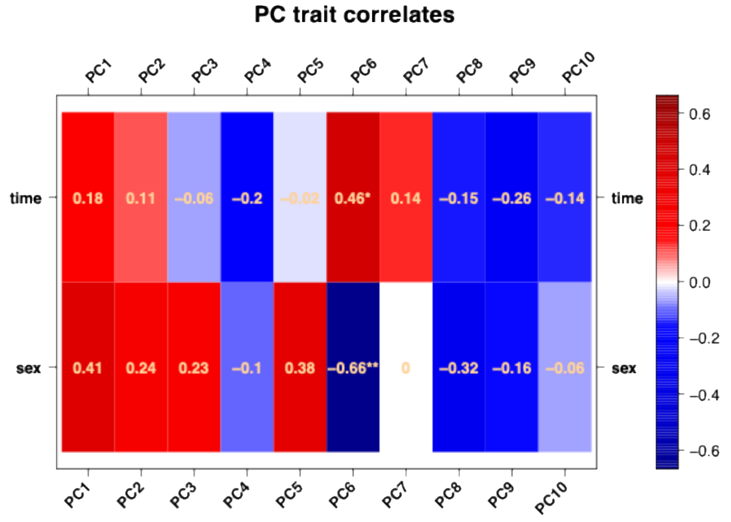

### Meaburn: day 2 PCA before filtering genes

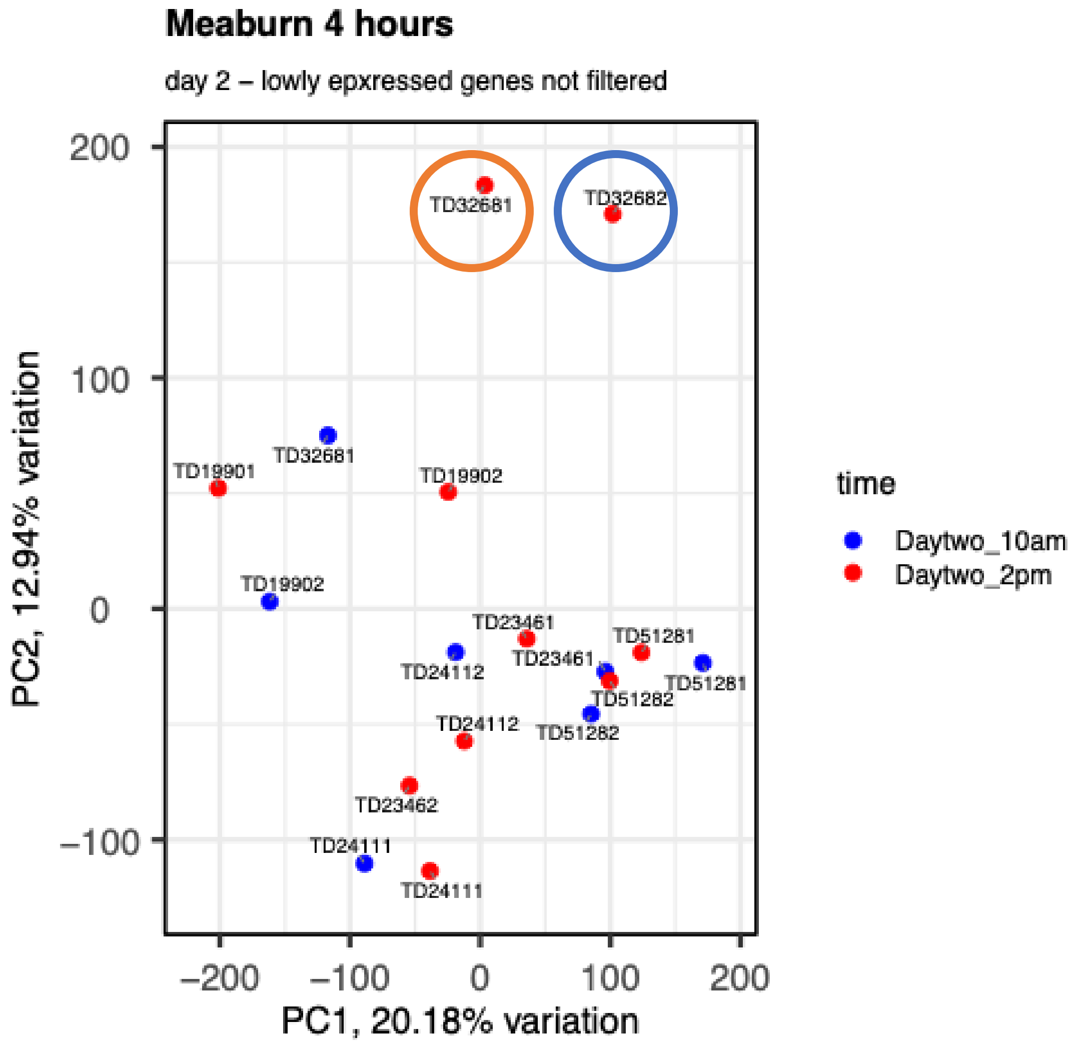


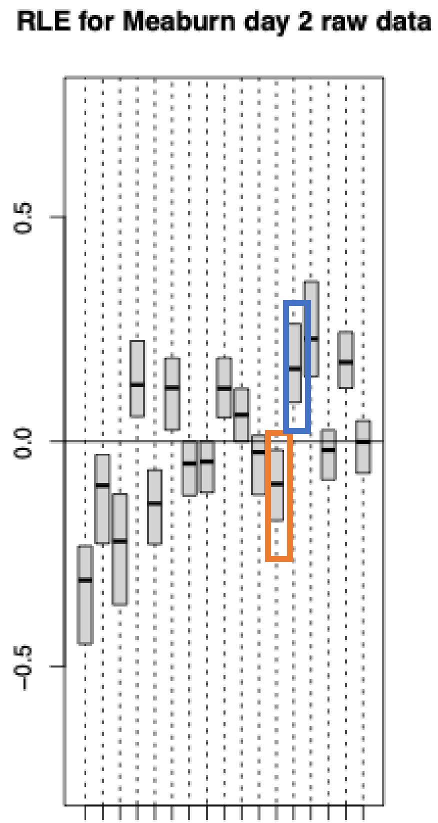


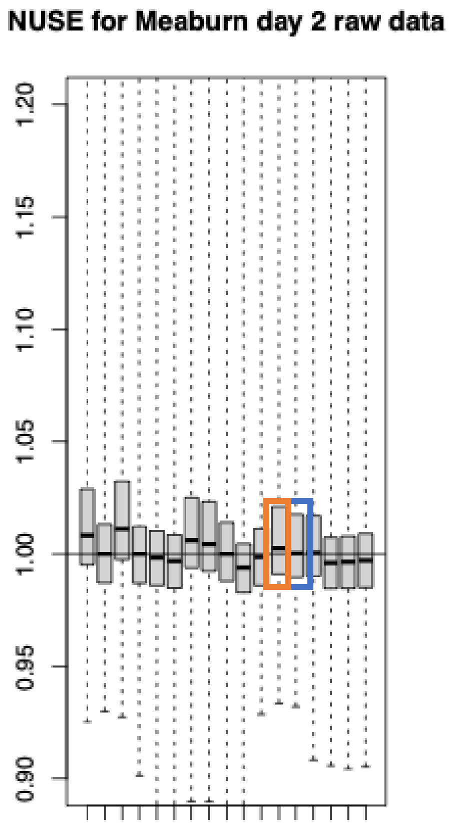

The University of Texas at Austin
Waggoner Center for
Alcohol & Addiction Research

### Meaburn: day 2 PCA before filtering genes

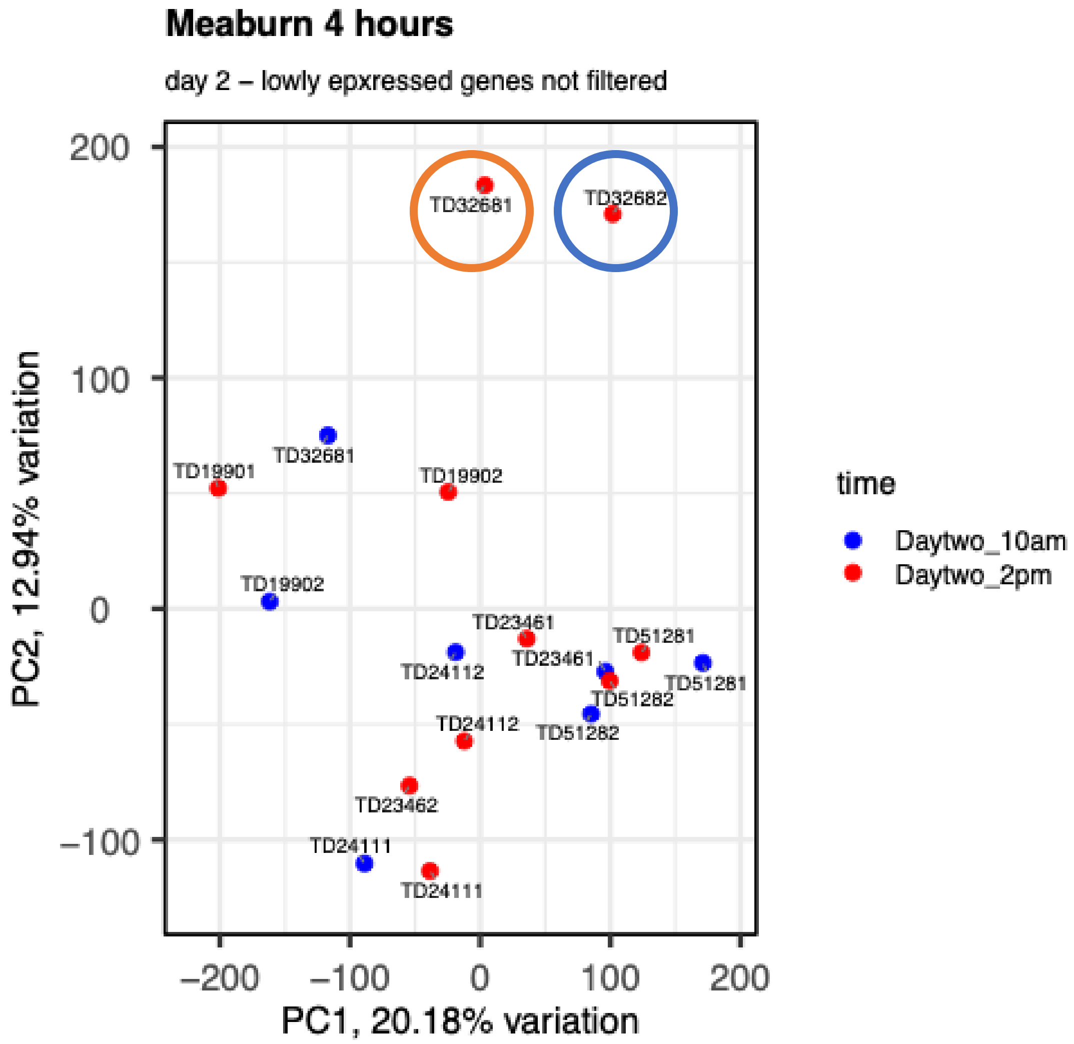


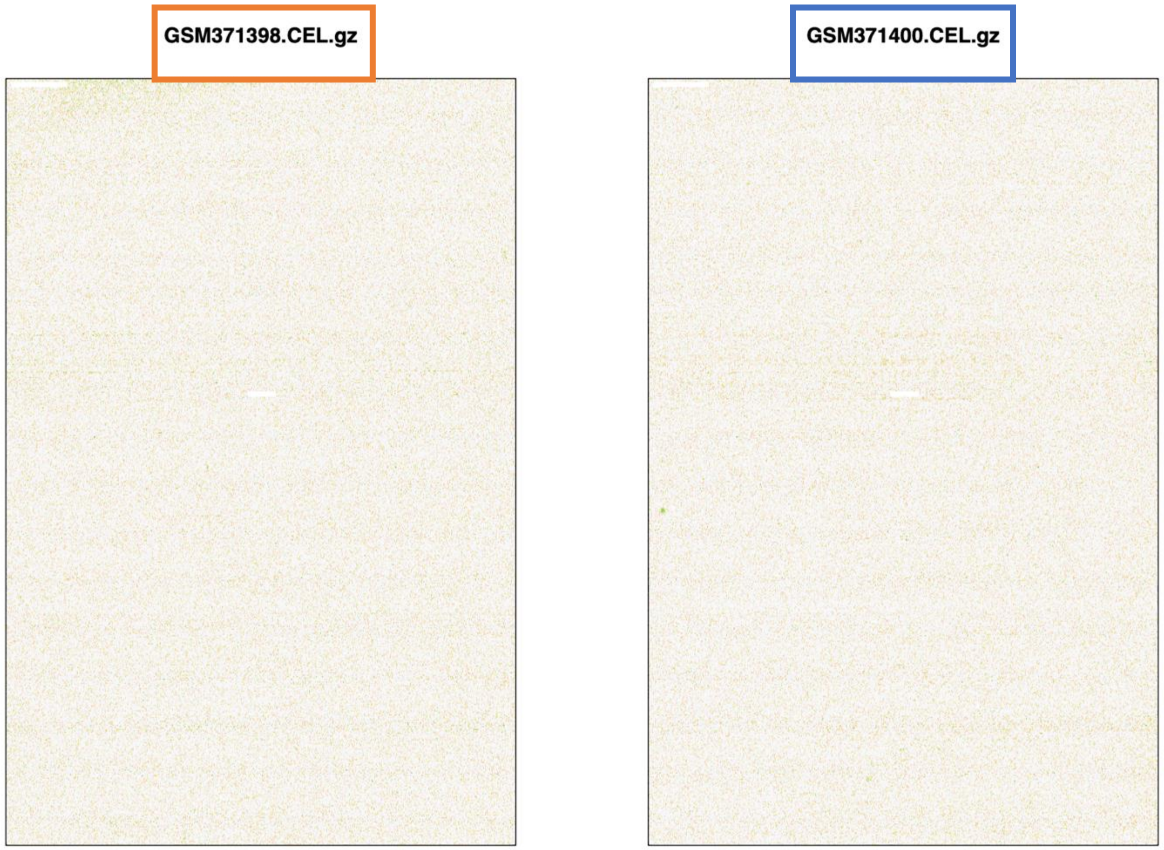


Because these two samples don't have anomalous quality statistics or pseudoimage abnormalities, we kept both of these samples in the analysis.

The University of Texas at Austin
Waggoner Center for
Alcohol & Addiction Research

### Meaburn: day 2 PCA after filtering genes

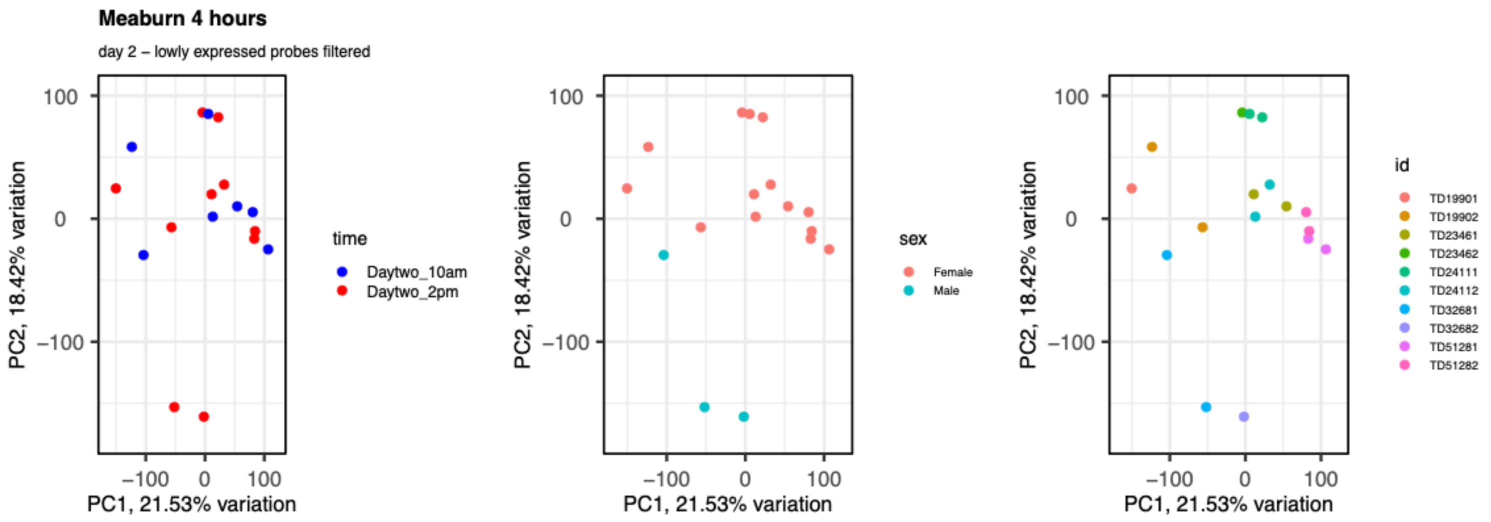


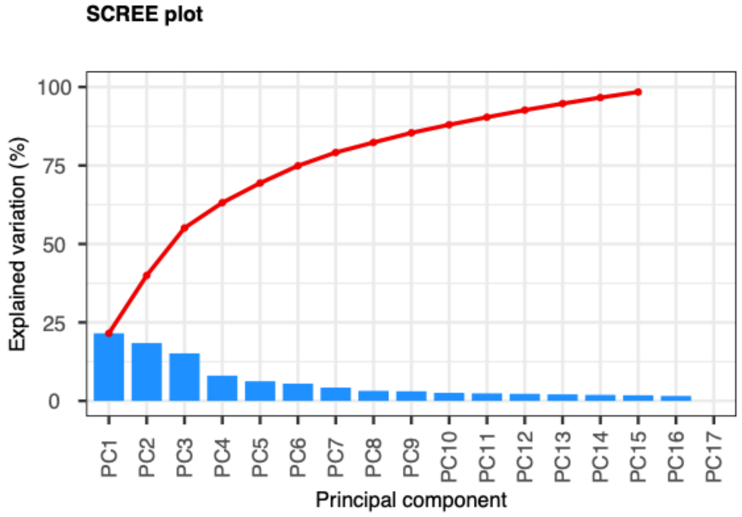


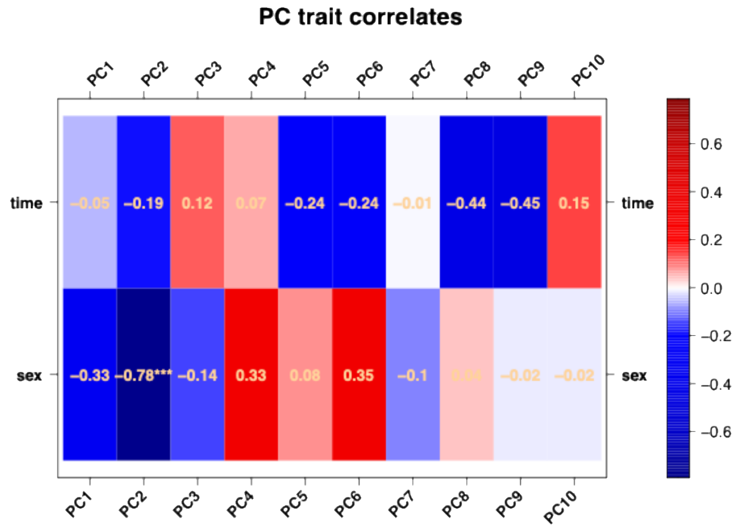

### Karlovich et al. (2009)

Microarray study, blood was drawn from healthy volunteers at baseline, 14 days, 28 days, 90 days, and 180 days. The researchers state that hybridization was performed in two batches. Batch 1 contained the baseline, day 14, and day 28 samples. Batch 2 contained the day 90 and day 180 samples. An initial analysis confirmed these batches using Principal Component Analysis (not shown). We analyzed batch 1 and batch 2 separately. The researchers also state that RNA isolation occurred as the samples arrived, adding another potential batch effect to the data. However, the samples did not cluster by day in PCA, so we assume that this effect was negligible across samples.

### Karlovich et al. (2009) - summary

| Sample | Subject ID | Sex | Time Point | Reason Removed |
|---|---|---|---|---|
| GSM401087 | 121 | Male | Day 90 | Elevated NUSE statistic, and an outlier on PCA plots pre- and post-gene filtering |
| GSM401148 | 172 | Female | Day 180 | Elevated NUSE statistic, pseudoimage abnormalities, and an outlier on PCA plots pre- and post-gene filtering |
| GSM401157 | 174 | Female | Day 90 | Authors state that this participant was diagnosed with lung cancer on day 70 of the study. This array also has abnormal quality control metrics. |

### Karlovich: Batch 1 RLE, NUSE, and pseudoimages

Minor aberrations– the effect on variability can be assessed with a PCA

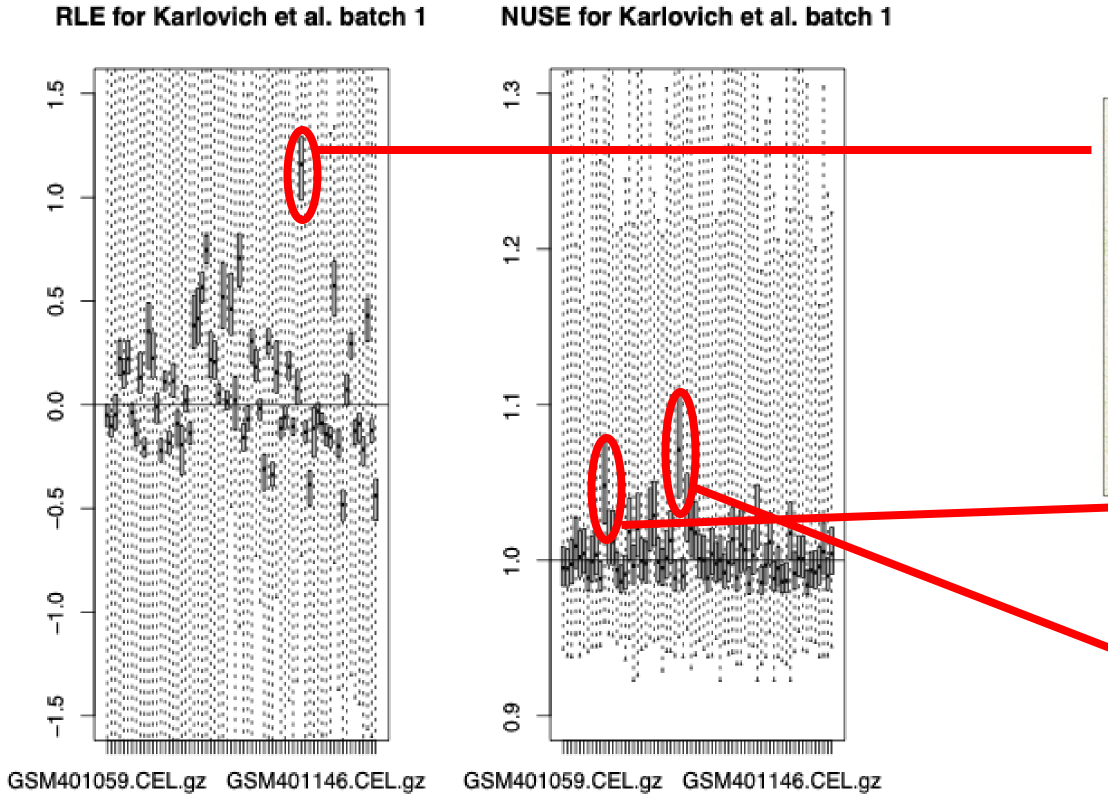


**Batch 1**

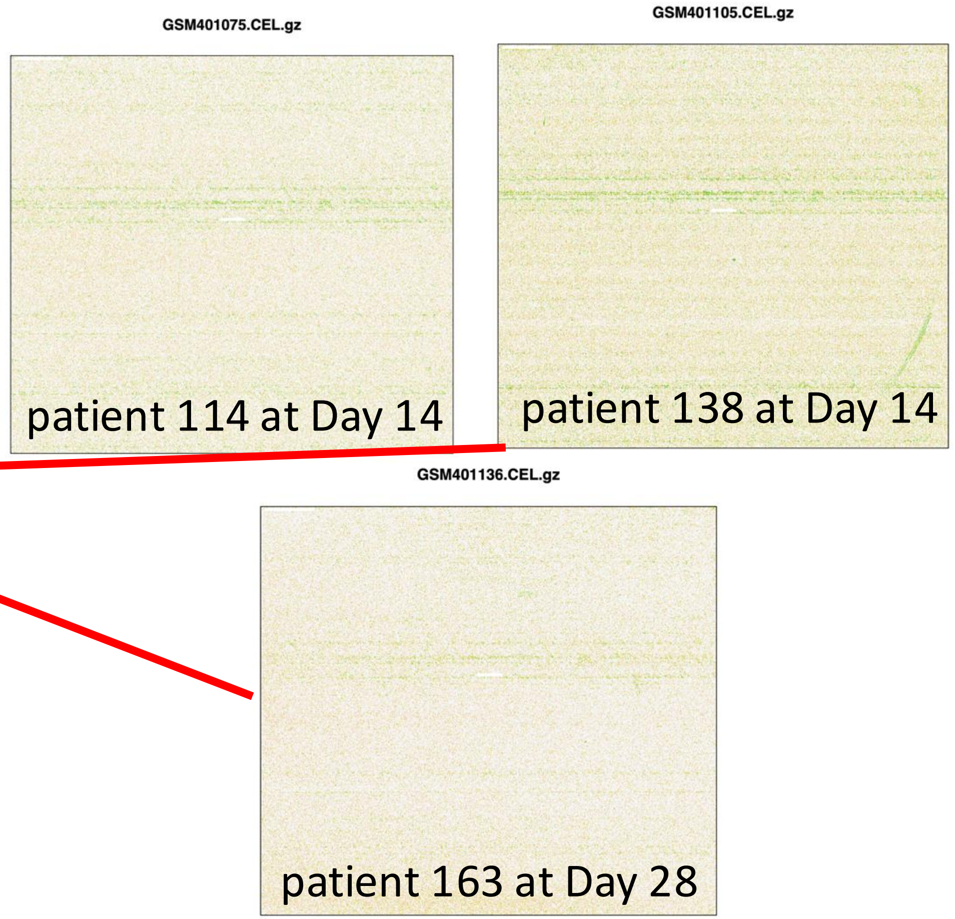

The University of Texas at Austin
Waggoner Center for
Alcohol & Addiction Research

### Karlovich: Batch 1 unfiltered vs. filtered PCA

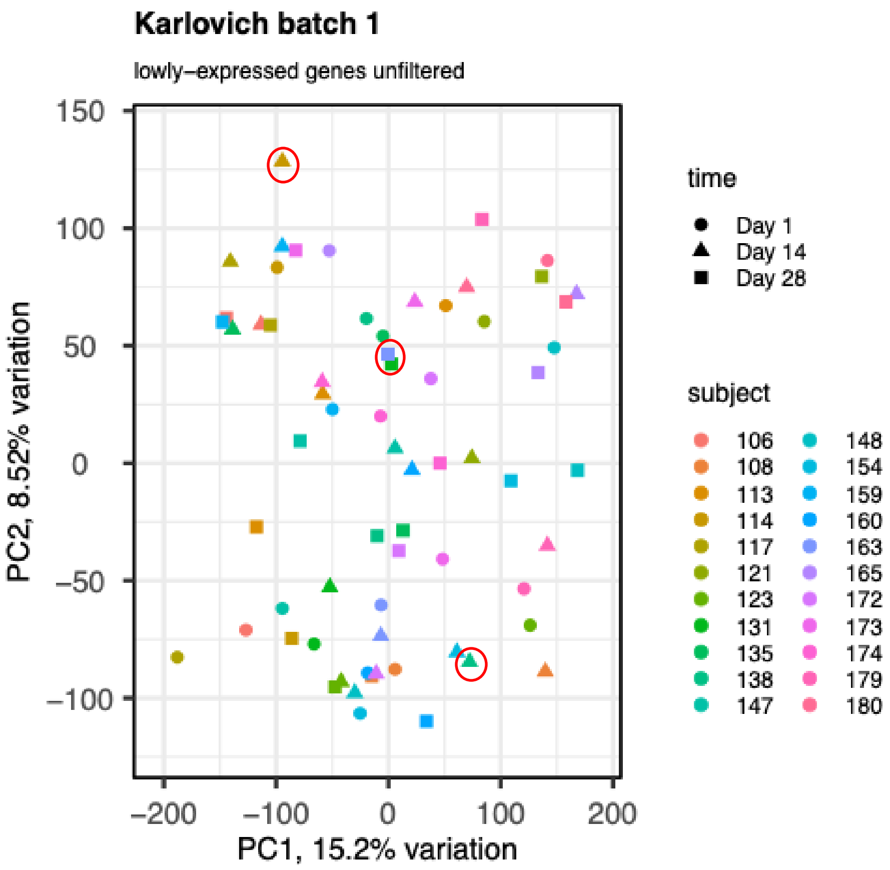


*Filter lowly-expressed genes*

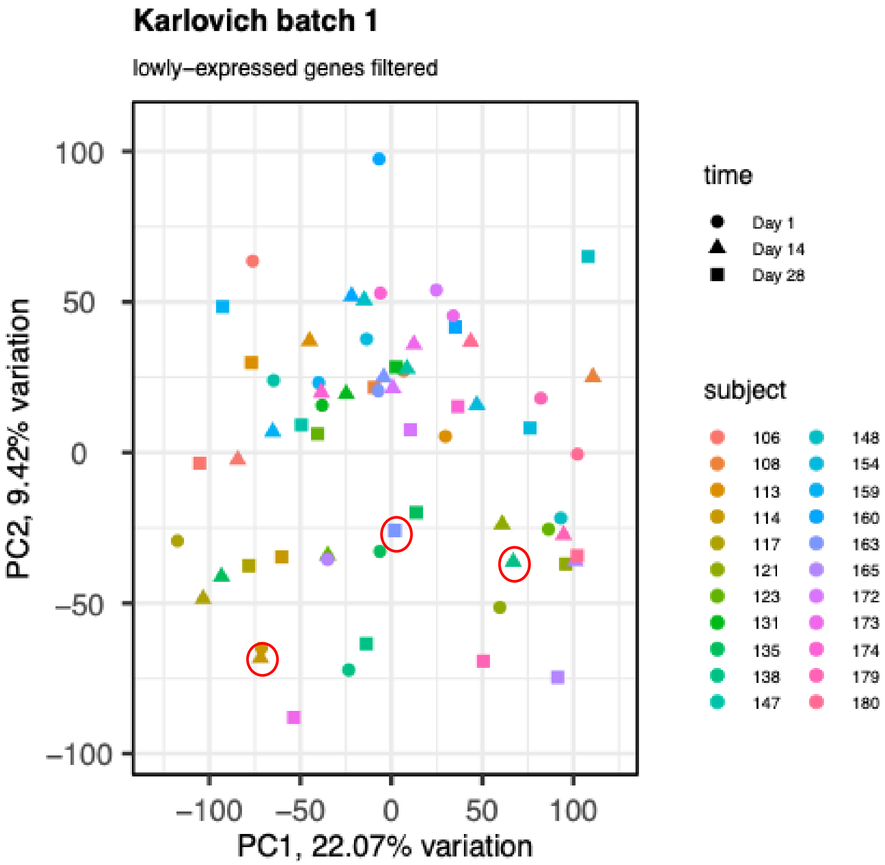


There appears to be little to no effect, so we decided to keep these samples for analysis.

### Karlovich: batch 1 PCA after sample removal and filtering

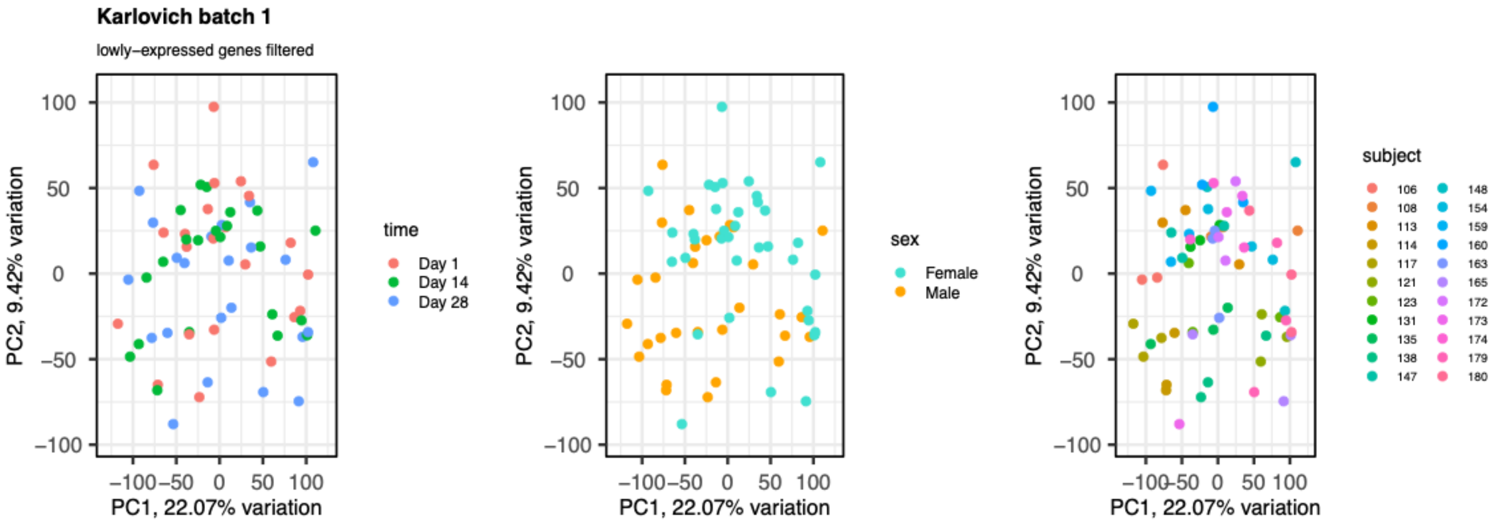


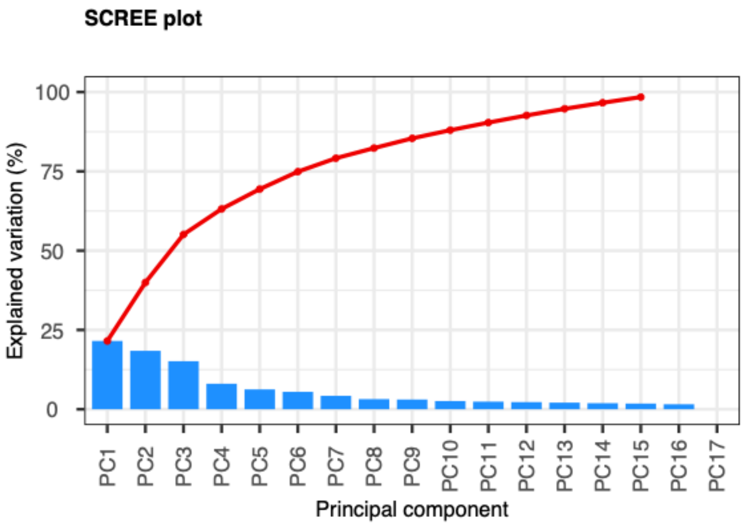


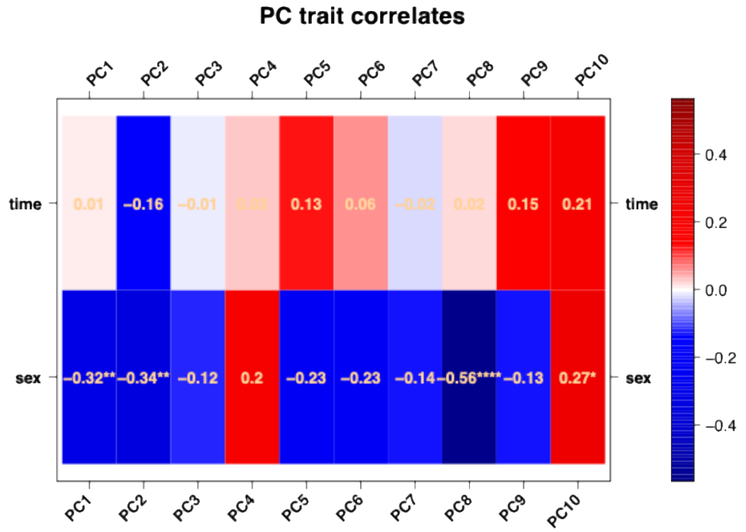

The University of Texas at Austin
Waggoner Center for
Alcohol & Addiction Research

### Karlovich: RLE, NUSE, and pseudoimages

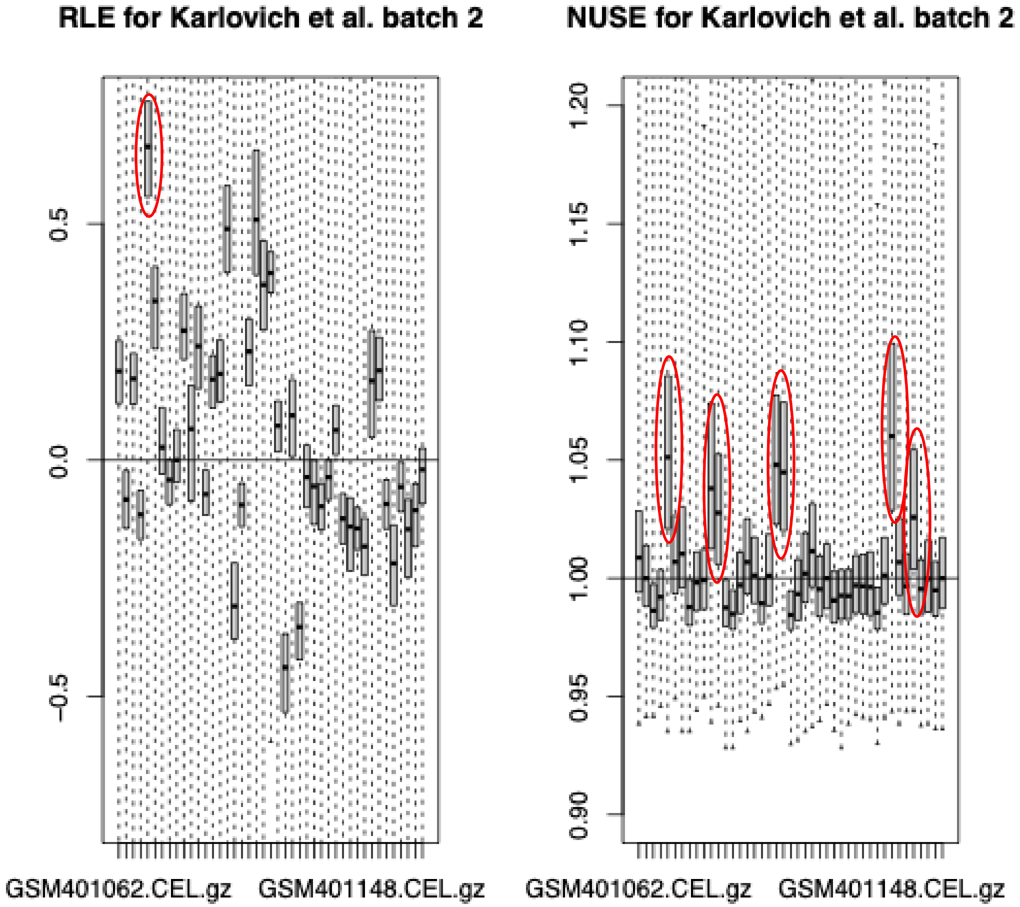


**Batch 2**

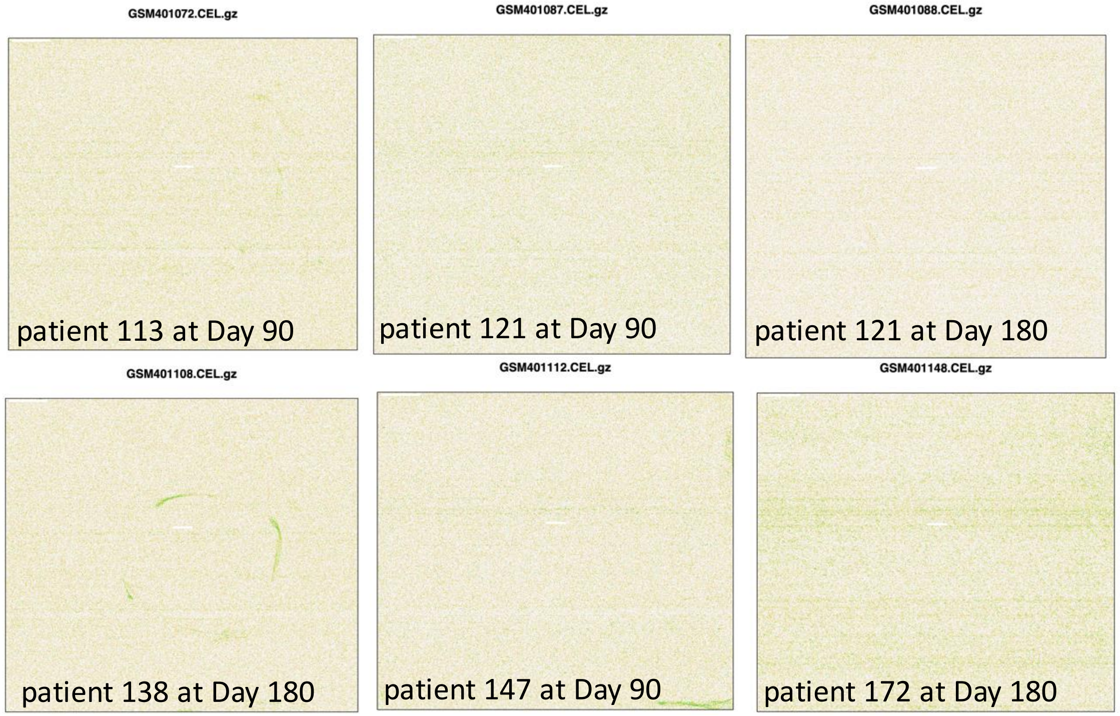


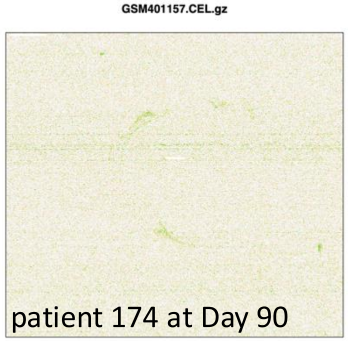


From Karlovich et al: "Whole genome expression profiles were generated for all samples at scheduled time points (Days 1, 14, 28, 90, and 180) in the subset of 22 subjects with the exception of the Day 180 time point for subject 174 who died prior to the end of the study." It is stated later in the study that the cause of death was lung cancer, diagnosed 70 days into the study. **For this reason, we removed GSM401157** from the analysis.

The University of Texas at Austin
Waggoner Center for
Alcohol & Addiction Research

### Karlovich: batch 2 filtered vs. unfiltered PCA

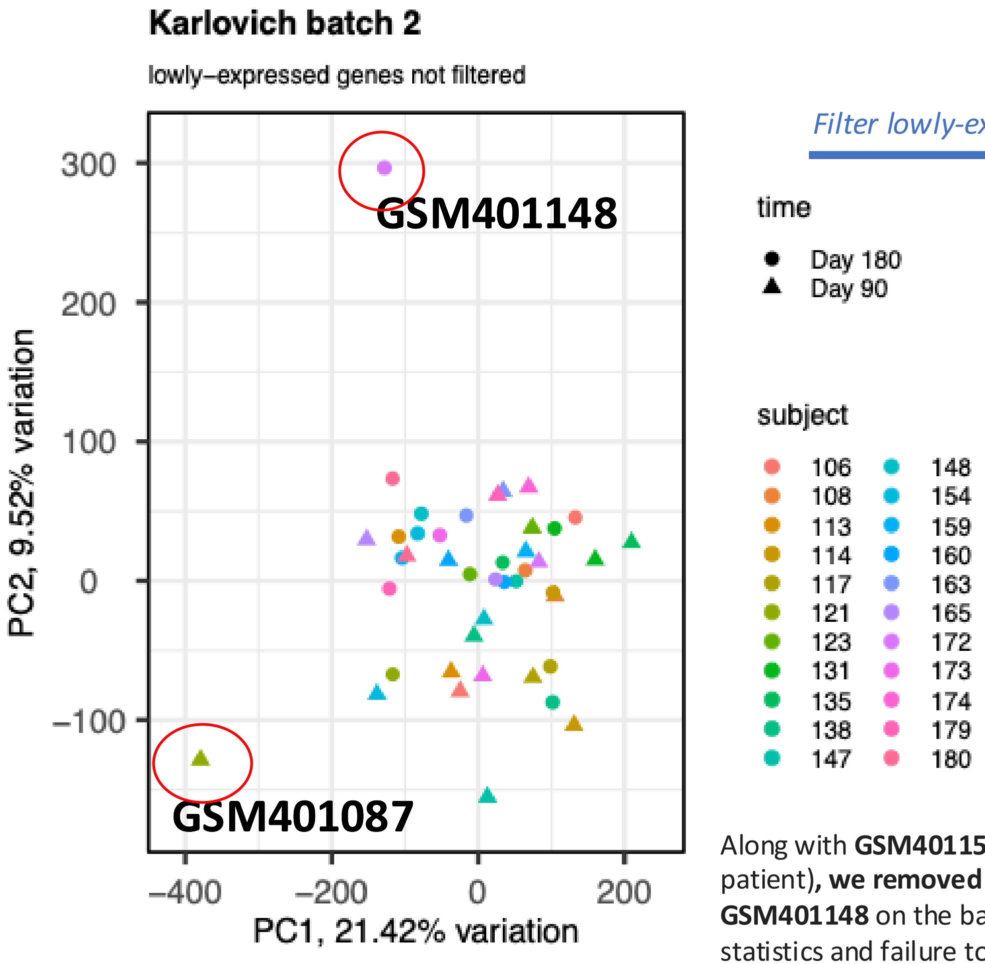


*Filter lowly-expressed genes*

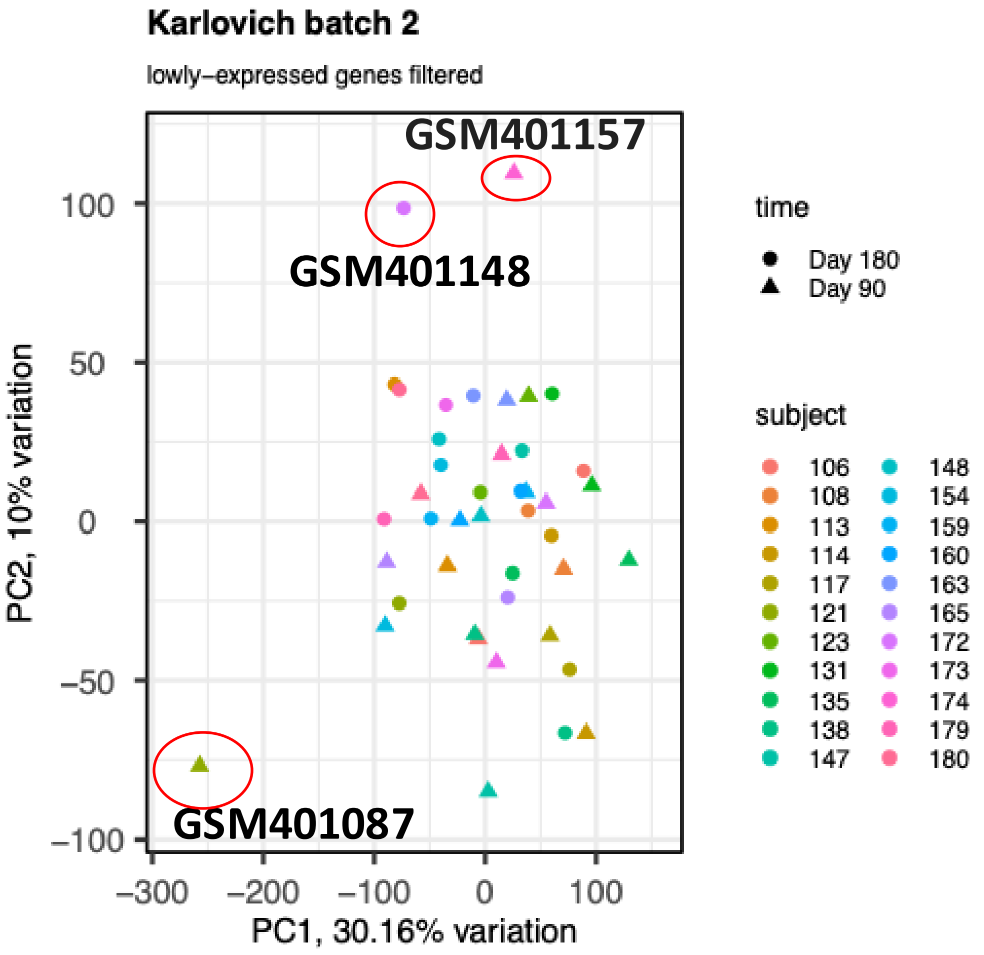


Along with **GSM401157** (lung cancer patient)**, we removed GSM401087 and GSM401148** on the basis of their NUSE statistics and failure to cluster with the experiment on PCA.

The University of Texas at Austin
Waggoner Center for
Alcohol & Addiction Research

### Karlovich: batch 2 PCA after sample removal and filtering

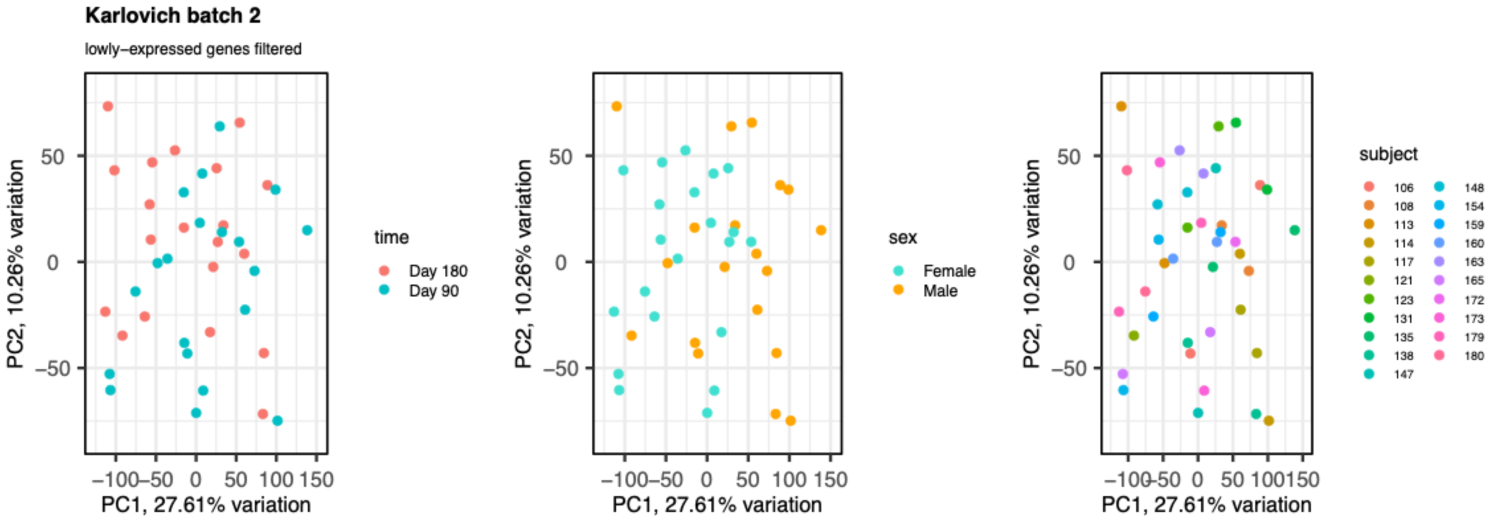


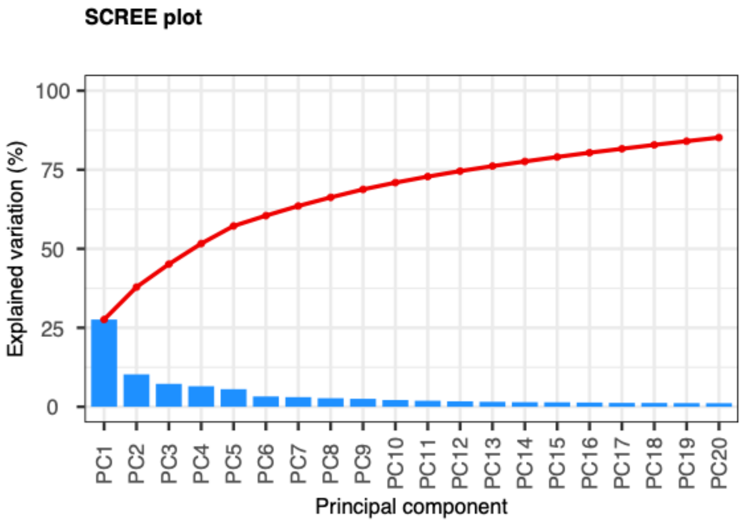


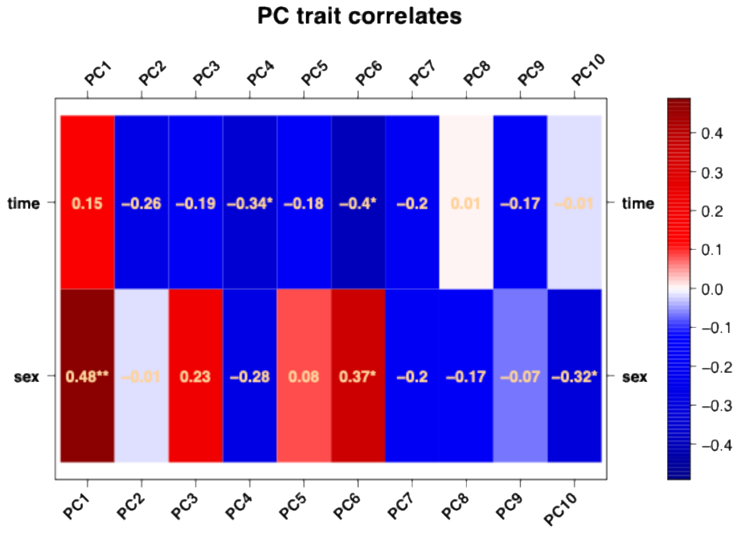

### Karlovich: Plots of features (L) and samples (R) for all samples after removing outlier samples

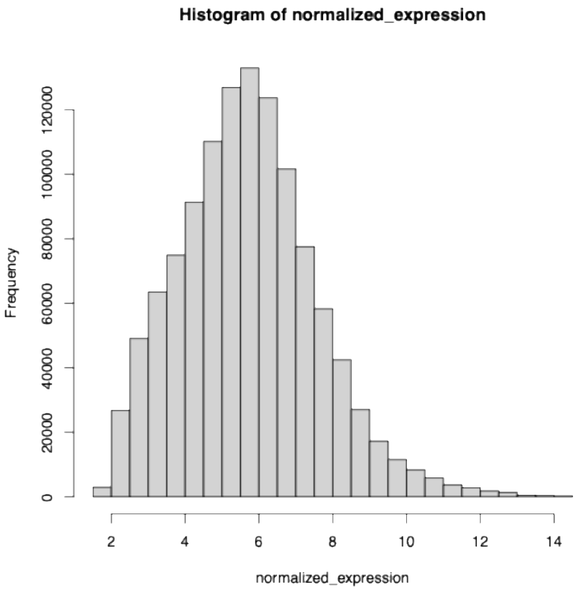


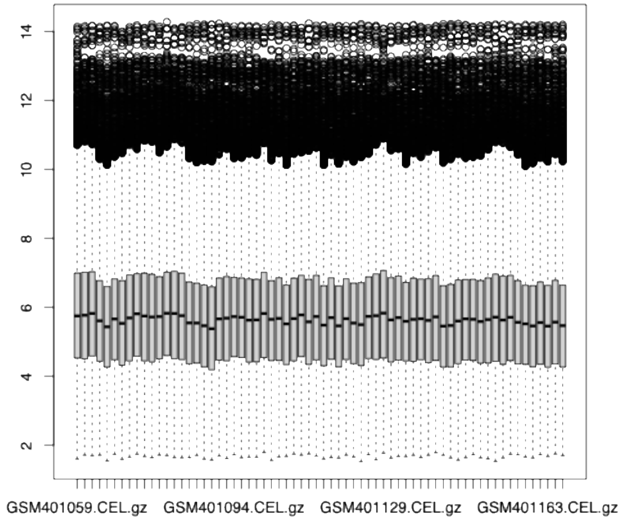


Batch 1

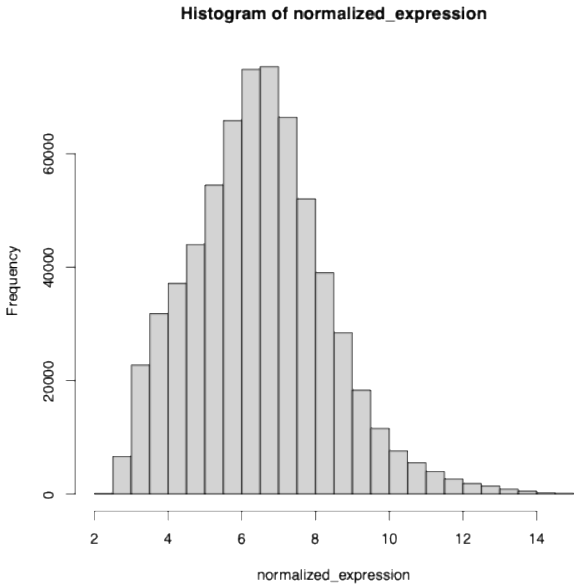


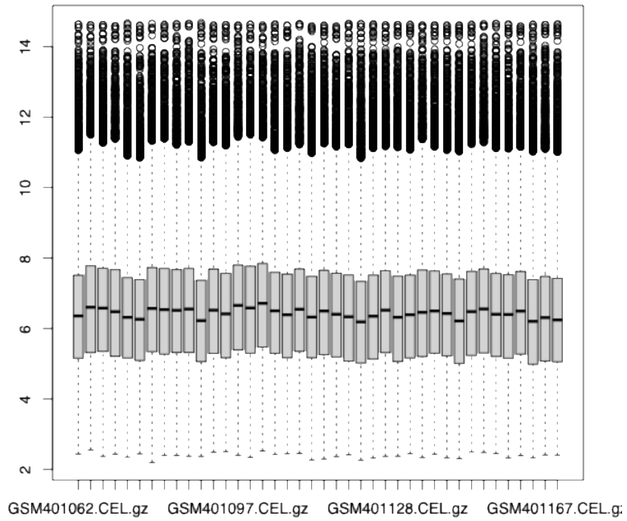


Batch 2
