## Supplementary Figures for "The Blood RNA Stability Atlas: defining temporal structure and trait-state programs in the human whole-blood transcriptome"

### Slide 1
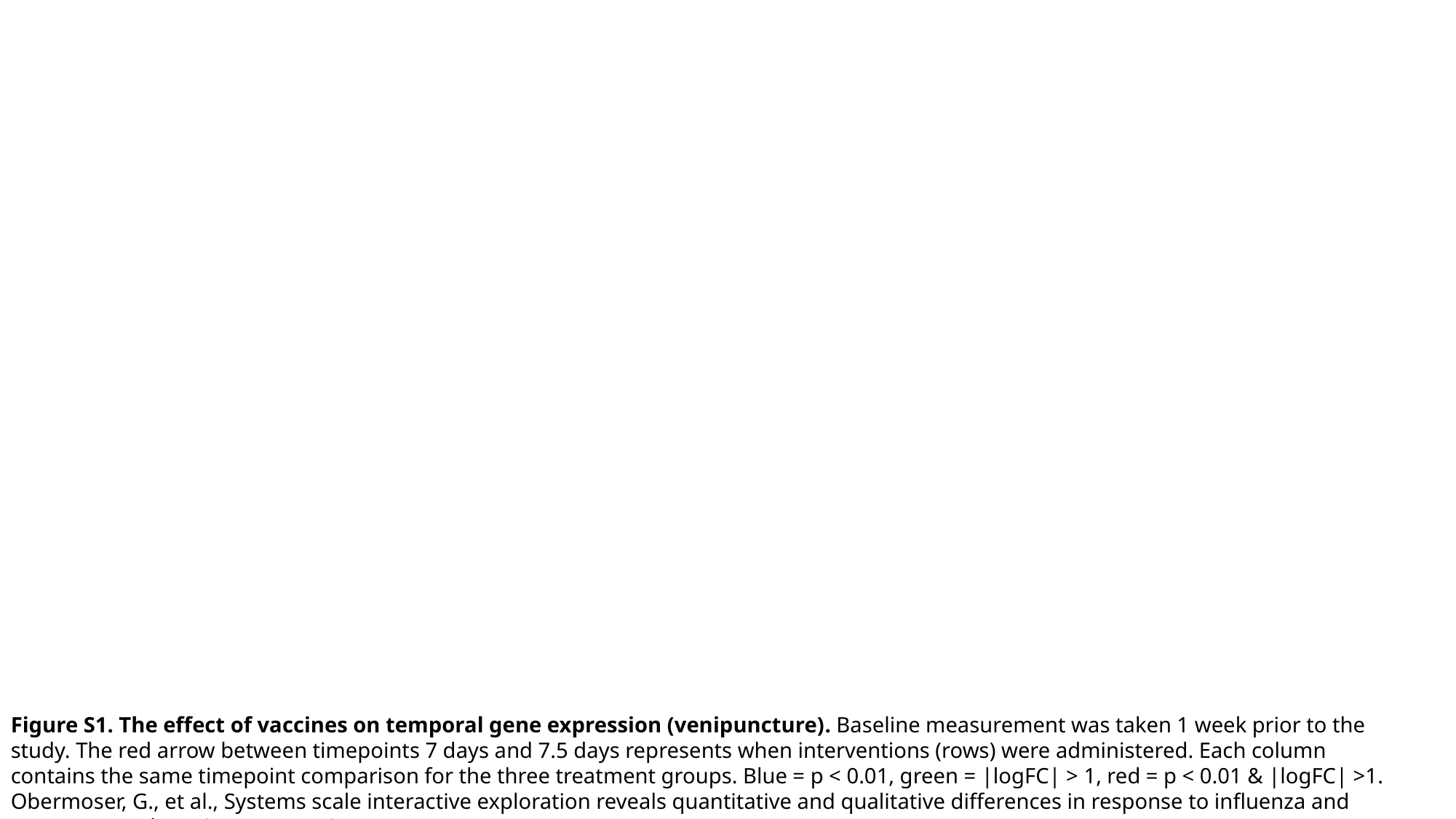

Figure S1. The effect of vaccines on temporal gene expression (venipuncture). Baseline measurement was taken 1 week prior to the study. The red arrow between timepoints 7 days and 7.5 days represents when interventions (rows) were administered. Each column contains the same timepoint comparison for the three treatment groups. Blue = p < 0.01, green = |logFC| > 1, red = p < 0.01 & |logFC| >1. Obermoser, G., et al., Systems scale interactive exploration reveals quantitative and qualitative differences in response to influenza and pneumococcal vaccines. Immunity, 2013. 38(4): p. 831-44.

### Slide 2
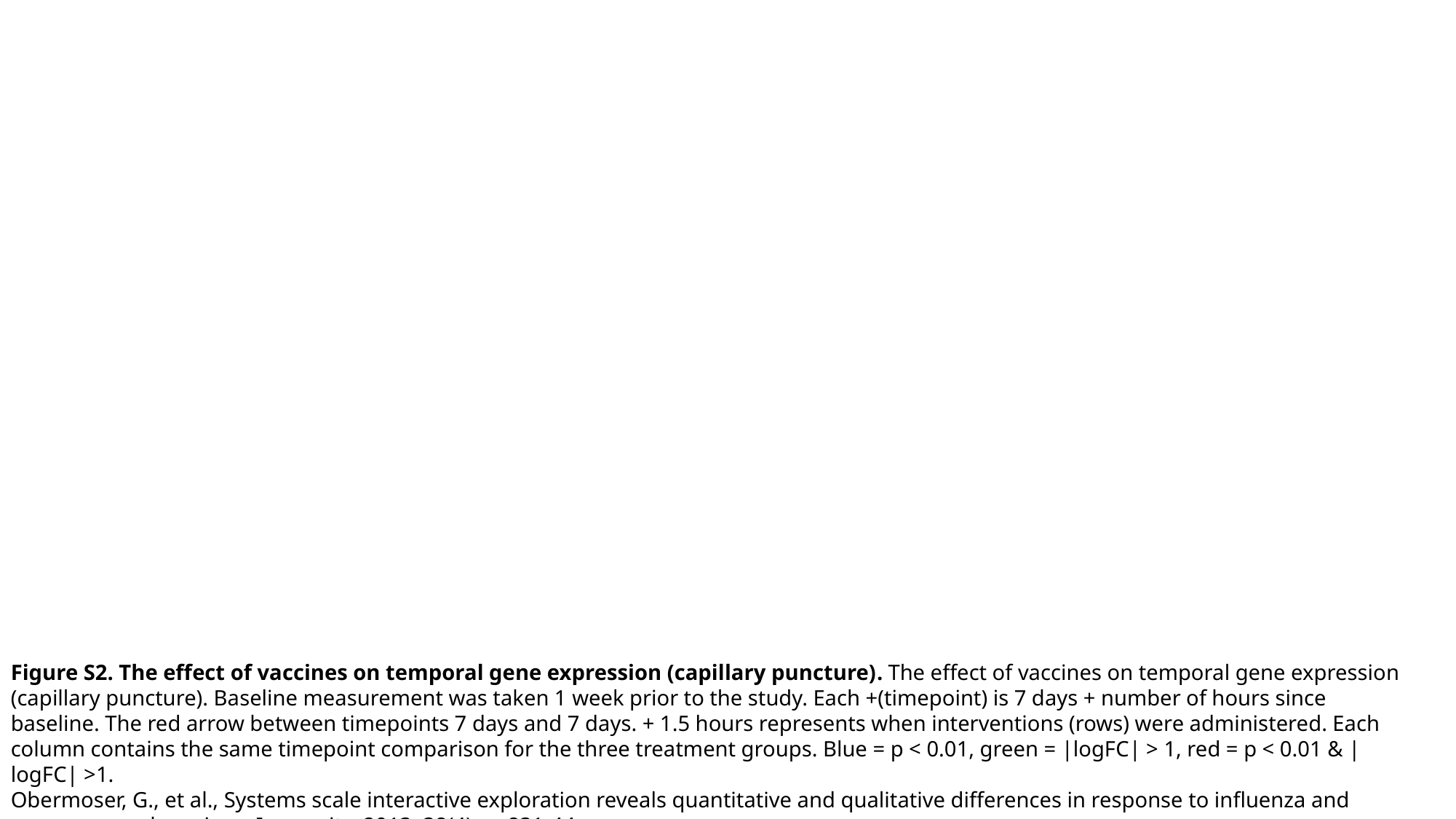

Figure S2. The effect of vaccines on temporal gene expression (capillary puncture). The effect of vaccines on temporal gene expression (capillary puncture). Baseline measurement was taken 1 week prior to the study. Each +(timepoint) is 7 days + number of hours since baseline. The red arrow between timepoints 7 days and 7 days. + 1.5 hours represents when interventions (rows) were administered. Each column contains the same timepoint comparison for the three treatment groups. Blue = p < 0.01, green = |logFC| > 1, red = p < 0.01 & |logFC| >1.
Obermoser, G., et al., Systems scale interactive exploration reveals quantitative and qualitative differences in response to influenza and pneumococcal vaccines. Immunity, 2013. 38(4): p. 831-44.

### Slide 3
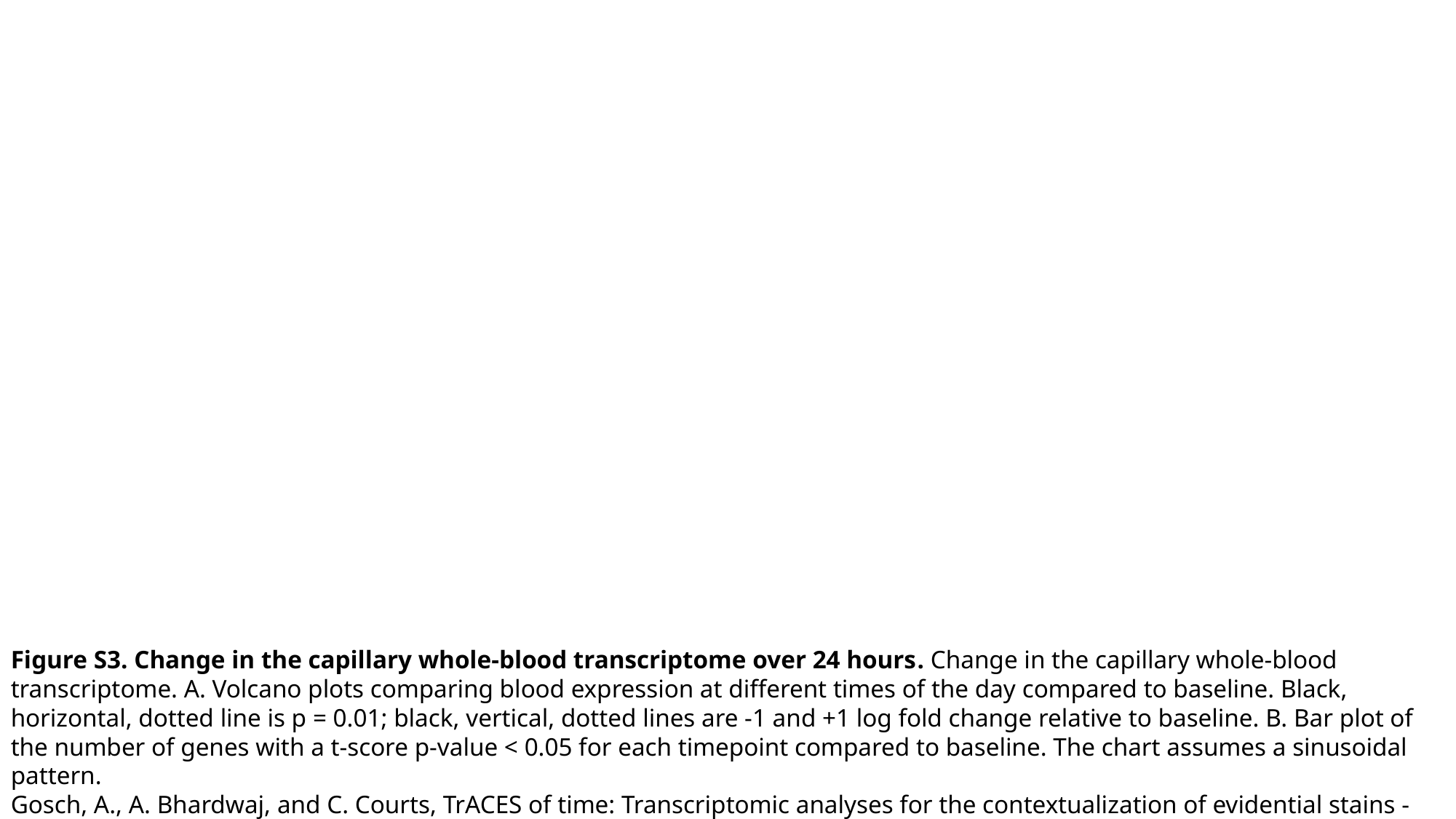

Figure S3. Change in the capillary whole-blood transcriptome over 24 hours. Change in the capillary whole-blood transcriptome. A. Volcano plots comparing blood expression at different times of the day compared to baseline. Black, horizontal, dotted line is p = 0.01; black, vertical, dotted lines are -1 and +1 log fold change relative to baseline. B. Bar plot of the number of genes with a t-score p-value < 0.05 for each timepoint compared to baseline. The chart assumes a sinusoidal pattern.
Gosch, A., A. Bhardwaj, and C. Courts, TrACES of time: Transcriptomic analyses for the contextualization of evidential stains - Identification of RNA markers for estimating time-of-day of bloodstain deposition. Forensic Sci Int Genet, 2023. 67: p. 102915.

### Slide 4
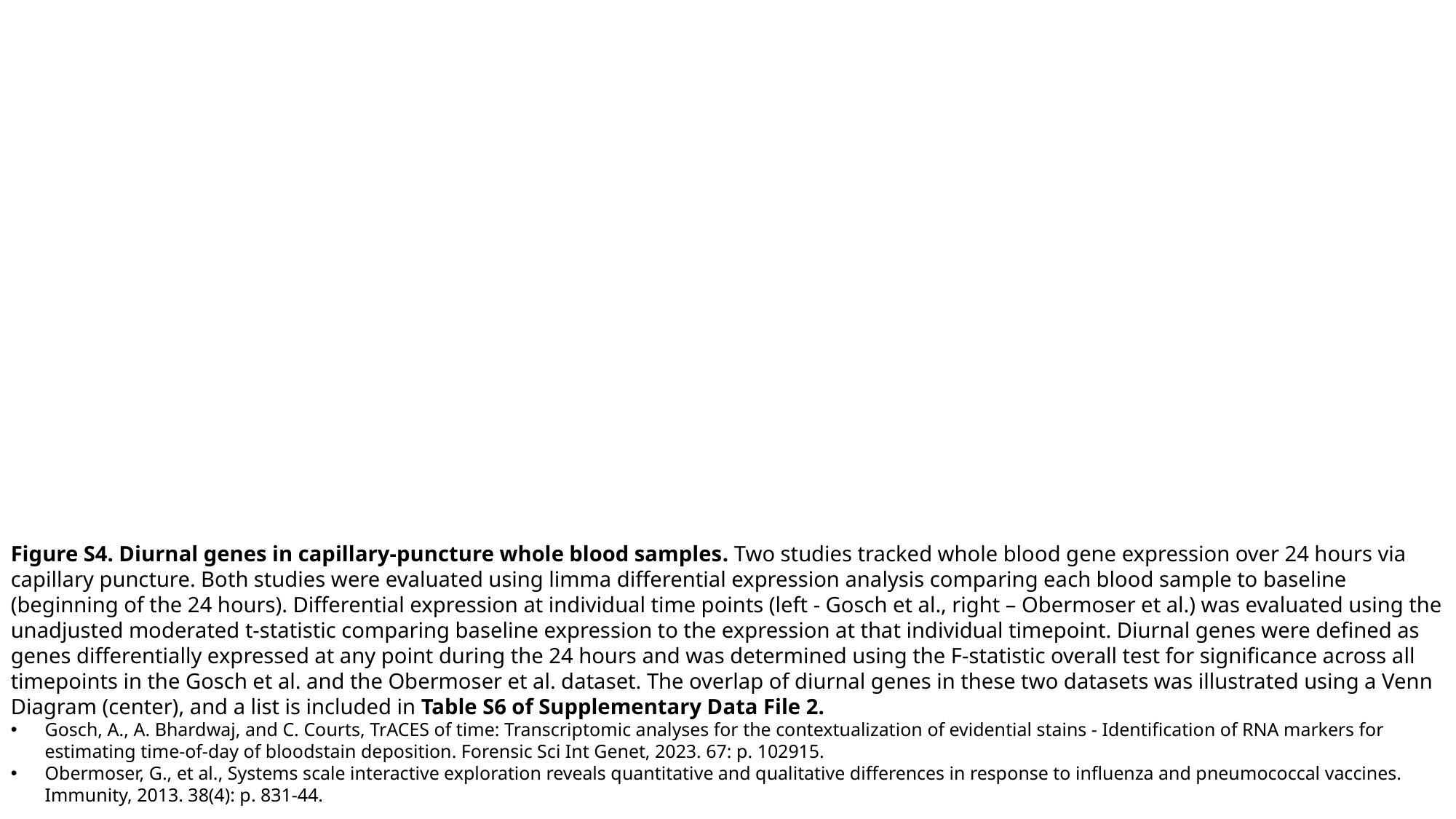

Figure S4. Diurnal genes in capillary-puncture whole blood samples. Two studies tracked whole blood gene expression over 24 hours via capillary puncture. Both studies were evaluated using limma differential expression analysis comparing each blood sample to baseline (beginning of the 24 hours). Differential expression at individual time points (left - Gosch et al., right – Obermoser et al.) was evaluated using the unadjusted moderated t-statistic comparing baseline expression to the expression at that individual timepoint. Diurnal genes were defined as genes differentially expressed at any point during the 24 hours and was determined using the F-statistic overall test for significance across all timepoints in the Gosch et al. and the Obermoser et al. dataset. The overlap of diurnal genes in these two datasets was illustrated using a Venn Diagram (center), and a list is included in Table S6 of Supplementary Data File 2.
Gosch, A., A. Bhardwaj, and C. Courts, TrACES of time: Transcriptomic analyses for the contextualization of evidential stains - Identification of RNA markers for estimating time-of-day of bloodstain deposition. Forensic Sci Int Genet, 2023. 67: p. 102915.
Obermoser, G., et al., Systems scale interactive exploration reveals quantitative and qualitative differences in response to influenza and pneumococcal vaccines. Immunity, 2013. 38(4): p. 831-44.
